# Hormonal induction, quality assessments and the influence of seasonality on male reproductive viability in a long-term managed *ex situ* breeding colony of Southern Rocky Mountain Boreal toads, *Anaxyrus boreas boreas*

**DOI:** 10.1101/2023.08.23.554154

**Authors:** N.E. Calatayud, Leah Jacobs, Gina Della Togna, Cecilia J. Langhorne, Amanda C. Mullen, Rose Upton

## Abstract

The Southern Rocky Mountain Boreal Toad (*Anaxyrus boreas boreas)* is an *ex situ* managed species which relies heavily on the use of assisted reproductive technologies to augment existing *in situ* populations. Despite the use of ARTs to manipulate reproduction of this species, the *ex situ* program continues to face challenges with annual reproduction. Human chorionic gonadotropin (hCG) at 10 IU/gbw singly or in combination with 0.4 ug/gbw GnRH-A have been successfully reported for this species, however, with a number of other available hormones, it is not clear if these are the most effective protocols for this species. Moreover, there is no information on how exogenous hormone administration is affected by other factors such as seasonality. Therefore, in the first part of this study, we compare the efficacy of the 10 IU/g hCG and 0.4 Lg/g Gonadotropin Releasing Hormone agonist (GnRH-A) administered singly or in combination, as well as GnRH-Apd + Amphiplex (0.4 Lg/g GnRH-A + 10 Lg/g Metoclopramide [MET] - a dopamine antagonist), or 10 Lg/g MET alone. Spermiation responses and sperm viability were compared across treatments with hormonal curves then correlated to seasonality. The results of this study suggest that the optimal hormonal stimulation protocol, across all treatments, in terms of sperm quality was 10 IU/g hCG + 0.4 Lg/g GnRH-A. Further optimization is required, in particular the exploration of higher doses of GnRH-A. Lastly, we observed that the effect of seasonality influenced the hormonal efficacy and magnitude of the spermiation response. As was expected, quality and concentration parameters were affected by the month in which hormone administration occurred.

**HIGHLIGHTS:**

- Spermiation in the Southern Rocky Mountain Boreal toad, *Anaxyrus boreas boreas,* is elicited most effectively by hCG singly or in combination with GnRH-A.
- Spermiation responses lasted up to 96 hours post injection (hpi) though quantity and quality parameters were highest in the first 12 hpi.
- Total motility, forward progressive motility and percentage live cells all indicated *A. b. boreas* sperm quality was in a good range.
- Acrosome integrity requires more research as it was comparatively lower than expected.
- Responses to hormone inductions are seasonally affected in this species but suggest semi-discontinuous cycling.
- Ex situ breeding should perform better sperm assessments before designing breeding strategies.

## INTRODUCTION

*Ex situ* breeding programs (ECPs) have been globally recognized as necessary mitigation strategies to safeguard against extinction and loss of biodiversity [1]. Additionally, ECPs play a large role in facilitating ecosystem restoration, and assist in the management of fragmented, dwindling or shrinking populations while combatting growing pandemics that exacerbate species declines [1,2]. As one of its critical roles, *ex situ* management must preserve genetically robust populations that exhibit high levels of fitness, preserve adaptive viability and, where possible, maximize genetic exchange with wild populations [3–5]. A growing number of studies are modeling the optimum targets required for the genetics of a species to be adequately managed *ex situ,* which, when combined with developing assisted reproductive and genomic technologies, augment the impact of ECPs on species conservation:[6–9]. Factors relating to inbreeding, loss of gene flow and reduced genetic variation, require interdependent analysis of a species’ evolution both *in situ* and *ex situ* since viable populations that are required for species survival estimates differ markedly under both conditions [4,10–12].

Despite the urgent need, taxonomic prioritization of amphibians (and reptiles) continues to be overlooked [13]. The last reported assessments of the world’s threatened species estimated that ∼3% are held in zoological institutions [13] while more recent reports by Johnson et al. (2021) recommended 398 species for ECP management, of which372 are recommended for further management with additional genome resource banking (GRB) [1,14,15]. With the increasing need for *ex situ* management, GRBs, along with the development of ARTs, will likely continue to grow financially and logistically, facilitating better strategic preservation of genetically valuable individuals without the need to expand colony numbers or extirpate animals from the wild ([6,7,16].

Particularly in the last decade, the growing implementation of ARTs in amphibian conservation has provided a number of approaches that help with increasing breeding output in ECPs, by exogenous hormone induction of spermiation, ovulation and breeding behaviors, artificial fertilization, refrigerated/cold, short-term storage and gamete cryopreservation [1,8,9,17–19]. Examples of conservation programs that are actively implementing ARTs in their recovery strategies include, the Mountain yellow-legged frog, *Rana muscosa* [20], Puerto Rican Crested Toad, *Peltophryne lemur* [21], the Chinese Giant Salamander, *Andrias davidianus* [22]*, Atelopus zetteki, A. varius, A. limosus, A. certus, A. glyphus, Triprion spinosus* and *Craugastor evanesco* [8,23] and the Oregon spotted frog, *Rana pretiosa* (Hunter, B., 2023, MSc Thesis). However, in order to implement best practices when using ARTs for conservation, it is important to conduct parallel research on the life history, reproductive viability, husbandry requirements, impacts of long-term captivity and genetic diversity of a species as part of any strategic conservation planning [1,15]. Incorporating a multi-faceted approach to preserve and augment genetic diversity, requires interlinking *ex situ* with *in situ* conservation strategies to promote gene flow across wild populations and ECPs [21], Hunter, B., 2023 MSc thesis). These strategies should also aim to augment reproductive output, especially in long-lived ECPs, in the face of loss of relevant genetic diversity, adaptive fitness and reproductive viability due to multi-generational domestication [1,3,4,24,25].

The Southern Rocky Mountain Boreal Toad (*A. boreas boreas)* is representative of an *ex situ* managed species where ARTs are regularly implemented to augment existing *ex situ* populations, in addition to hatching and rearing individuals brought in from the wild. *A. b. boreas* is a subspecies of the North American Western Toad and has a distribution restricted to the montane wetlands of southeastern Idaho, Western and South-Central Wyoming, most of Utah (except western Box Elder County), Colorado, and North-Central New Mexico (USFW, 2017). As with many amphibian species, the current status of *A. b. boreas* ongoing survival is inextricably linked to the prevalence of Chytridiomycosis *(Bd)* and the species or its distinct populations’ resilience to it. Declines of *A. b. boreas* date back to the mid-1980’s in Colorado and South-Central Wyoming but more specifically, within the Southern Rocky Mountains, extirpation from 6 of 20 known sites have been linked to *Bd* invasion between 2003-2013 [26,27]. *Ex situ* breeding, captive rearing and reintroductions have been at the forefront of conservation for *A. b. boreas* since 2000. However, within the *ex situ* population, reproduction has been variable and low with few studies investigating the incorporation of ARTs to increase reproductive success in tandem with naturalistic husbandry practices (e.g. brumation) [28–32].

The Native Aquatic Species Restoration Facility (NASRF) in Colorado, has housed the largest *ex situ* colony of *A. b. Boreas* in existence for over two decades, and serves as an important insurance colony for a number of genetically unique populations which are bred for reintroduction and translocation purposes. In 2012-2013, prior to this study, ARTs were incorporated in an effort to enhance the low reproductive viability that has been characteristic of this and other anuran ECPs. In 2012, a combination of hormone induced breeding, as well as artificial fertilization, led to the production of ∼24,500 and 1,710 tadpoles from 14 pairs, respectively. In 2013, 31,000 eggs, 642 embryos and 582 tadpoles were produced from 13 pairs and reintroduced to the wild and 59 retained for brood stock (Smith, T., 2014, SRM Boreal Toad Annual Report). As in most amphibian ECPs, some factors such as the length of time and adaptation to captivity, genetic diversity, lack of gamete synchronization and/or inappropriate husbandry or environmental cues, i.e. seasonality, in addition to an extended latency to reach sexual maturity could be responsible for poor reproductive viability [29,30]. The development of hormonal stimulation protocols has been at the forefront of NASRF’s breeding efforts and to date, four studies have reported on exogenous hormone protocol development in male and female *A. b. Boreas* [28–30,32]. Langhorne et al., (2021) reported that spermiation could be successfully induced using a single injection of hCG at three concentrations (5, 10 and 15 IU/gbw) and found that 10 IU/gbw was the most efficient in eliciting good sperm concentrations and motility. However, the latter did not report on other quality metrics relating to viability. Additionally, the existence of other available hormones have not been reported, therefore it is not clear if 10 IU/gbw hCG is the optimal protocol for this species, if viability assessments are included, or how its effect is influenced by other factors such as seasonality. Therefore, in this study, we compare the efficacy of the originally reported 10 IU/gbw hCG protocol, to a number of hormones and combinations that have previously been reported as successful for other species. We chose 0.4 Lg/g GnRH-A singly or in combination with 10 IU/g hCG, GnRH-Apd + Amphiplex (0.4 Lg/g GnRH-A + 10 Lg/g Metoclopramide (MET) - a dopamine antagonist), or a single dose of 10 Lg/g MET, which have been reported to work in a number of continental-wide American anurans [23,33–36]. Spermiation responses and one sperm viability metric (proportion of live sperm across all treatments and acrosome integrity) were compared across treatments along with hormone response curves and spermiation peaks, then correlated to seasonality as a factor influencing the efficacy of the treatments.

## 1. METHODS

### 2.1 Animals

The individuals used in this study were maintained at the Native Aquatic Species Restoration Facility in Alamosa, Colorado (NASRF). Males representing 13 source populations across 11 localities in Colorado including: Eagle, Grand, Hinsdale, Jackson, Larimer, Mesa, Mineral, Pitkin, Routt, Chaffee and Clear Creek^1^ were selected for a comparative study that examined: 1) the effects of exogenous hormones on sperm quality parameters including concentration, motility and viability and 2) the effect of hormones on male reproductive seasonality (measured month to month from May - September 2014 and June - August 2015). Over the course of the two years, 2014-2015, 219 males were used for hormone data collection while another 105 males were used as control animals and were injected with saline amphibian ringers (SARs). The minimum time between hormonal stimulation for individual’s that were reused more than once was four weeks, a time frame selected based on McDonough et. al., 2016, [37].

### 2.2 Housing & husbandry

#### Housing during brumation

Males and females were housed together in groups according to the locality of origin. Brumation took place between December and May at temperatures between 2–6°C in an EcoPro G2 1350 liter upright refrigerated cabinet (Foster refrigerator, Corp., Hudson, New York, USA) in plastic boxes (33 x 13 x 15 cm) lined with a layer of activated carbon, moistened sand (3.81 cm deep) and moistened sphagnum moss.

#### Housing during active season (breeding and foraging)

Males and females were housed in multi-generational groups of 5 - 20 according to their locality of origin. During the active season (breeding and foraging periods), toads were housed in rectangular fiberglass tanks (121 x 60 x 30 cm) tilted at a 20° angle to allow constant drainage of free–flowing groundwater (Figure 1). Artificial lighting (Zoo Med 2 Pack of T8 ReptiSun 10.0 UVB, 24 Inch, 17-watt, Reptile Habitat Lighting) was provided and adjusted to coincide with natural photoperiod. Crickets and mealworms were gut–loaded with Bug Burger®, Hydro–load® water replacement gut–load (Allen Repashy’s®, La Jolla, California, USA) and fresh carrots, 24 hours prior to being fed to the toads. Toads were fed *ad libitum* 3 times per week.

**Figure 1.**
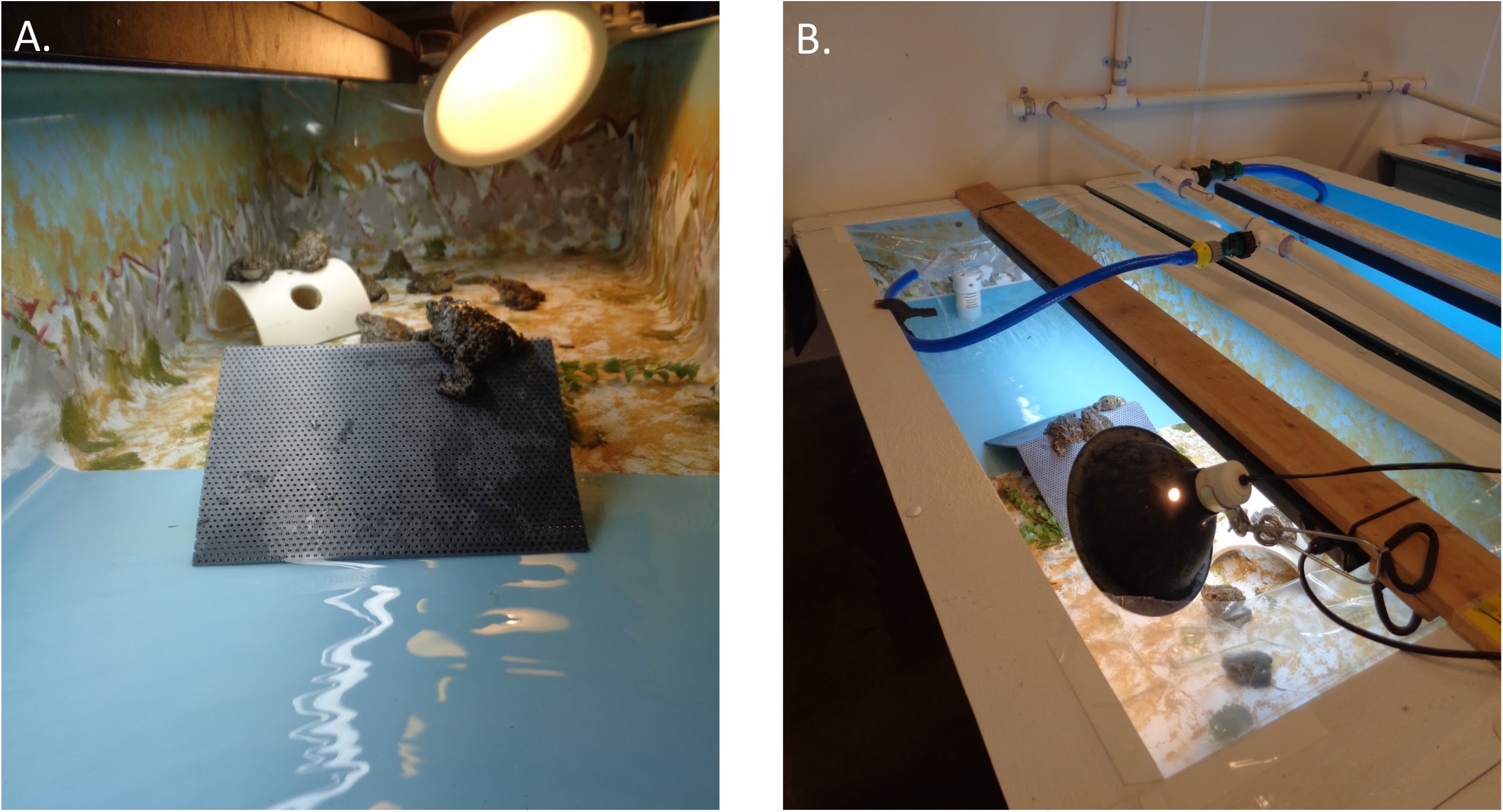
Toads were housed in cohorts according to county of origin. Per large fiberglass tanks, the number of animals varied between 5 - 20 individuals. Artificial light and other enrichment items were offered for movement and basking purposes.

### 2.3 Hormonal stimulation

Human chorionic gonadotropin (hCG) and Gonadotropin-Releasing Hormone agonist (GnRH-A) individually or in combination (hCG + GnRH-A), GnRH-A priming dose + GnRH-Apd + Amphiplex (GnRH-A + Metoclopramide) and Metoclopramide (MET) alone, were tested (Table 1). All hormones were reconstituted in sterile saline amphibian ringer’s (SAR) and injected intra–peritoneally (IP) using 27–gauge needles in a gram per body weight (gbw) basis [38]. Stock hormone solutions were stored in aliquots at –20°C and thawed prior to use.

**Table 1.** Exogenous hormones administered with their dosage based on body weight of the frog.

| Hormone treatment | Dose | Manufacturer | Reference |
| --- | --- | --- | --- |
| Human chorionic gonadotropin<br>hCG<br>**Total n = 77** | 10 IU/gbw | Sigma Aldrich, (cat # C1063) | Langhorne, 2016;<br>Langhorne et al., 2021 |
| Des-Gly10,D-Ala6,Pro-NHEt9)-GnRH acetate salt<br>GnRH-A<br>**Total n = 105** | 0.4 µg/gbw | BACHEM (cat #H-4070) | Calatayud et al., 2017 |
| hCG - GnRH-A<br>**Total n = 44** | 10 IU/gbw + 0.4 µg/gbw | Sigma Aldrich, (cat # C1063) + BACHEM (cat #H-4070) | Calatayud et al., 2017 |
| GnRH-Apd + Amphiplex<br>**Total n = 30** | 0.4 µg/gbw GnRH-A + GnRH-Apd + Amphiplex (0.4 µg/gbw GnRH-A + 10 µg/gbw MET ) | GnRH-A (BACHEM, cat #H-4070) + MET (Sigma Aldrich, cat# M0763) | Trudeau et al., 2010;<br>Della Tonga et al., 2017 |
| Metoclopramide (MET)<br>**Total n = 44** | 10 µg/gbw MET | Sigma Aldrich, cat# M0763 | No prior report on singular use. |

In 2014, hormonal stimulations consisted of four treatment groups (hCG alone, hCG + GnRH-A, GnRH-A and GnRH-Apd + Amphiplex). In 2015, an additional treatment of MET at 10 Lg/gbw was added for a total of five treatment groups. Hormonal stimulations were conducted using a random block design, twice a week. During each collection day, five (2014) or six (2015) individuals were injected with each treatment, including a control male. Prior to injection, a time zero (t=0) was also collected from each of male, both hormonally treated and the control, and checked for the presence of sperm. The additional number of males in the hCG and GnRH-A treatment groups were added to this dataset from a separate artificial fertilization experiment.

### 2.4 Collection, quantity and quality parameters of hormone-induced spermatozoa: Concentration, motility and viability

During the study, male frogs were held separately in a plastic container (25.4 cm x 33.5 cm x 54.9 cm) with 1 inch of water to ensure continuous water absorption for frequent urination. To collect spermic urine, male frogs were held above a petri dish using the cradling method outlined in Calatayud et al., (2019). On the rare occasion where urination could not be manually stimulated as described above, spermic urine was obtained by inserting a small disposable vinyl catheter (0.34mm x 0.052mm) (Scientific Commodities cat #: BB31785-V/5) into the cloaca (as described in Calatayud et al., 2019).

To determine the best hormone protocol, spermic urine was collected after males were administered one of four or five (depending on the year) different hormone treatments (Table 1), across several time-points 0, 2, 3, 5, 7, 9, 12, 24-, 48-, 72- and 96-hours post injection (hpi). Data was collected in order to determine 1) the most effective hormone treatment, 2) the response duration (response curve) and 3) spermiation peaks. Sperm concentration peaks were calculated as previously described in [23] and Mouttham, et al., 2011, [39] and used to illustrate the best concentration peaks but were not statistically analyzed.

To define the spermiation peaks, cutoffs measurements were calculated using the average sperm concentration for each time point. Then, an average of all time points was calculated and added to 1x standard deviation from such calculation. Time points with sperm concentration equal or higher than the cutoff were selected and removed. This calculation was repeated with the remaining time points with a new cutoff calculated until there were no values equal to or higher than the cutoff. All time points that were equal to or higher than the cutoffs were recorded as part of the peak or peaks [23].

### 2.5 Sperm concentration

Sperm concentration was analyzed using a Bright Line hemocytometer (Millipore-Sigma, USA cat: #Z359629) and a phase contrast microscope (Nikon Eclipse 80i) (Calatayud, 2015 Supplementary Document). Briefly, samples of 10 μL of urine were loaded onto the haemocytometer either as a straight volume or, for ease of counting where samples had a high density of sperm as determined visually by viscosity, at a dilution of 1:5 (1 μL of sperm to 4 μL of water) or 1:10 (1 μL of sperm to 9 μL of water). In diluted samples, final sperm concentrations were calculated by multiplying the final concentration by 5 or 10 accordingly.

### 2.6 Sperm motility and forward progression

Sperm motility was assessed (as described in Langhorne et al., 2021) and recorded for every time-point collected (2, 3, 5, 7, 9, 12, 24-, 48-, 72- and 96-hpi): (1) sperm concentration, (2) percent motile sperm, and (3) forward progressive motility (Appendix 1).

### 2.7 Sperm viability and morphological assessment

In addition to motility and concentration, sperm were further evaluated for, 1) viability, or the proportion of live sperm in the sample and 2) acrosome integrity. For the former, sperm were stained with eosin-nigrosin and a modified Pope solution which could be used in the field for acrosome integrity assessment (see protocol below) to ascertain information on sperm plasma membrane and acrosome integrity, respectively. These assessments were completed using a microscope equipped with a camera (Nikon Eclipse 80i). Though not part of the analysis, acrosome integrity was only tested on sperm collected from the hCG + GnRH-A treatment (most effective treatment). Additionally, samples were prepared for scanning Electron Microscopy (SEM) to characterize and measure the spermatozoa.

### 2.8 Live staining with Eosin-nigrosin

Briefly, eosin-nigrosin was added to fresh sperm samples at a 1:1 dilution (10 μL sperm:10 μL eosin-nigrosin) on a clean slide, gently mixed together and incubated at room temperature for 1 minute. After 1 minute, the sample was gently spread across the slide and allowed to dry completely before conducting the final evaluation. Once dry, 100 sperm cells were counted and the number of live (cells with intact membranes that appeared white and did not absorb stain, Figure 7a) were recorded as proportion of those sperm that were dead (had absorbed the dye). Counts and photography were conducted at x40 (2.5 μm) and x100 (1 μm) magnification using a (Nikon Eclipse 80i) microscope.

### 2.9 Acrosome integrity

*2.10* Sperm samples were fixed in 4% paraformaldehyde (PFA) made up in phosphate-buffered solution (PBS), at a ratio of 1:4 (spermic urine: PFA) and incubated overnight at 4L. Samples were mounted with Permount™ Mounting Medium (Fisher Scientific, Pittsburgh, PA, USA) on Superfrost slides (Superfrost Plus ™, Fisher Scientific, cat#:12-550-15). Sperm were stained with a modified Pope Stain (50ml PBS, 0.5g Rose Bengal, 0.4975g Fast green) for 90 sec, air dried and mounted. Morphological assessments were carried out describing normal morphology broadly by the presence of an intact acrosome, head, and tail versus abnormal acrosome (missing or not intact) and tail (coiled, broken, with or without mitochondrial vesicle). One hundred sperm cells were counted and the proportion of sperm with an intact acrosome recorded (Figure 7b).

### 2.10 Statistical Analysis of the comparative Assessments of sperm concentration, motility and viability

To determine the effect of the hormone (hCG, hCG + GnRH-A, GnRH-A, MET and GnRH-Apd + Amphiplex) and time post-injection (0, 2, 3, 5, 7, 9, 12, 24, 48, 72 and 96 hpi) on sperm concentration (cells/mL) we used zero-inflated negative binomial models, using hormone type as explanation for excess zeroes in the model. Month was included as a random effect to explain overdispersion in the data. To determine the effect of hormone type/dose and time post-injection on total motility, forward-progressive motility and viability, we used beta binomial logistic regressions - interpreted as the proportion of successful cases (i.e., motile or membrane intact sperm) - with the total number of sperm counted equaling the weights for the model. Both Frog ID and month were used as random effects to explain overdispersion in these models. For motility and viability assessments, only data where at least 90 sperm were counted were included and thus the timepoints for 48 h, 72 h and 96 h, as well as all data for MET was excluded from the model due to not enough replication across all treatments.

To determine the effect of the month on sperm concentration (cells/mL) we used zero-inflated negative binomial models, using hormones as explanation for excess zeroes in the model. Frog ID was included as a random effect to explain overdispersion in the data and the months May and October were dropped from the analysis due to not enough replication across all treatments. To determine the effect of month on total motility, forward-progressive motility, and viability, we used we used beta binomial logistic regressions - interpreted as the proportion of successful cases (i.e., motile or membrane intact sperm) - with the total number of sperm counted equaling the weights for the model. Observation-level random effects were used to explain overdispersion in these models. For motility and viability assessments, only data where at least 90 sperm were counted were included and thus all data for MET, as well as the months May and October, was excluded from the model due to not enough replication across all treatments.

All GLMM modeling in this study was run in R version 4.2.1 (R Foundation for Statistical Computing) package glmmTMB [40]. QQ plot residuals, distribution, dispersion, and uniformity was assessed using ‘DHARMa’ [41]. As described in Upton et al. 2021, model estimated marginal means (EMMS) and 95% confidence limits were determined for each condition and back transformed to proportions. Odds ratios (ORs) comparing treatments were also generated using the package emmeans (version 1.4.8) [42] *Available at* https://CRAN.R-project.org/package=emmeans *[verified 10 March 2021]*). All graphing was completed using Prism Graphpad Version 10.0 [released July 2023].

## 2. RESULTS

### 3.1 Effects of hormone and collection time points on sperm concentration

Sperm concentration ranged from a bottom average of 3.521 ×10^5^ (GnRH-Apd + Amphiplex) up to a top 2.38 x 10^6^ sperm cells/ mL (hCG + GnRH-A) when combined across all treatments and timepoints of collections. Concentration was significantly affected by both hormone treatment (LRT χ^2^(4) = 141.05, (DF 4, (LRT) L^05/2^, p< 2.2e-16) and the different time points [hours post-injection (hpi)] at which it was collected (LRT χ^2^(9) = 256.63, (DF 9, ^63/2,^ p< 2.2e-16). Sperm concentration ranged from a bottom average of 3.521 ×10^5^ (GnRH-Apd + Amphiplex) up to a top 2.38 x 10^6^ sperm cells/ mL (hCG + GnRH-A) when combined across all treatments and months of collection.The most effective treatment was hCG + GnRH-A followed by single administration of hCG, MET, GnRH-A and GnRH-Apd + Amphiplex respectively (Table 2).

**Table 2.** Sperm concentrations averaged across treatments and including all time points collected. CI = lower confidence limit = upper confidence limit.

| Hormone | EMM | LCI | UCI |
| --- | --- | --- | --- |
| hCG + GnRH-A | 1,631,171 | 1,330,519 | 1,999,760 |
| hCG | 1,101,581 | 887,229 | 1,367,721 |
| MET | 875,874 | 436,796 | 1,756,323 |
| GnRH-A | 494,506 | 356,625 | 685,697 |
| GnRH-Apd + Amphiplex | 240,351 | 179,842 | 321,218 |

Odds Ratios (OR) were used to further compare results between treatments. Comparatively, when averaged across all timepoints, hCG + GnRH-A produced the highest concentration of sperm, when compared to a singular dose of hCG (1.5 fold, Odds Ratio (OR) 1.481, 95% confidence interval (CI) 1.27 - 1.73), GnRH-Apd + Amphiplex (6 fold, OR 6.787, CI 5.00 - 9.21) and lastly GnRH-A alone (3 fold, OR 3.30, CI 2.27 - 4.79) (Table 3). When comparing hCG-GnRH-A to MET there was no statistically significant difference, despite the mean being 2-fold higher (2 fold, OR 1.862, CI 0.932 - 3.72)When administered singly, hCG was the next best treatment and was resulted in significantly higher sperm concentration when compared to GnRH-Apd + Amphiplex (OR 2.81, CI 1.55 - 5.08) (Table 2, Figure 2), GnRH-A alone (2 fold, OR 2.228, CI 1.526 - 3.25), but not significantly higher than from MET (OR 1.258, CI 0.633 - 2.499). MET the next best treatment, eliciting a significantly higher concentration than GnRH-Apd + Amphiplex (3-fold, OR 3.644, CI 1.757 - 7.56), and a higher but not significant difference when compared to GnRH-A alone (OR 1.77, CI 0.825 - 3.8). Finally, GnRH-A elicited a significantly higher concentration than GnRH-Apd + Amphiplex (2-fold, OR 2.057, CI 1.354-3.13) (Appendix 1 - 2).

**Table 3.** Table classifying hormone efficacy by statistical outcome in order of 1 - 4. Index is calculated based on a scoring value totaling 5. The order of efficacy is converted to a scale 1 = 1. 2 = 0.75, 3 = 0.5 and 4 = 0.25.

|  | **hCG +**<br>**GnRH-A** | **hCG** | **GnRH-A** | **GnRH-Apd + Amphiplex** |
| --- | --- | --- | --- | --- |
| Concentration (cells / mL) | 1 | 2 | 3 | 4 |
| Total motility (TM) | 2 | 1 | 4 | 3 |
| Forward progressive motility (FPM) | 1 | 1 | 2 | 3 |
| % Live cells | 1 | 1 | 1 | 1 |
| Response duration | 1 | 2 | 4 | 4 |
| Index | 4.75 | 4 | 2.75 | 2.5 |

**Figure 2.**
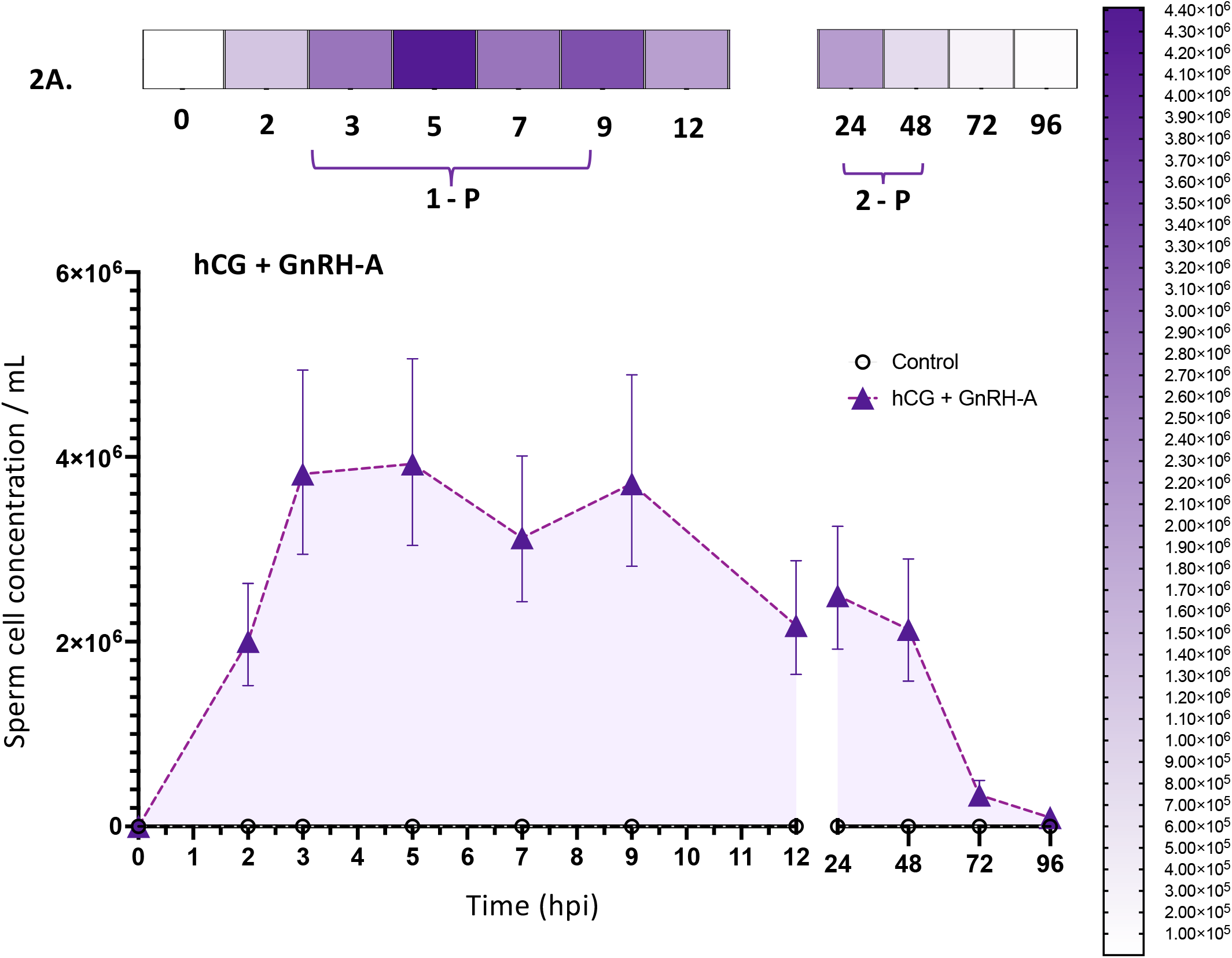
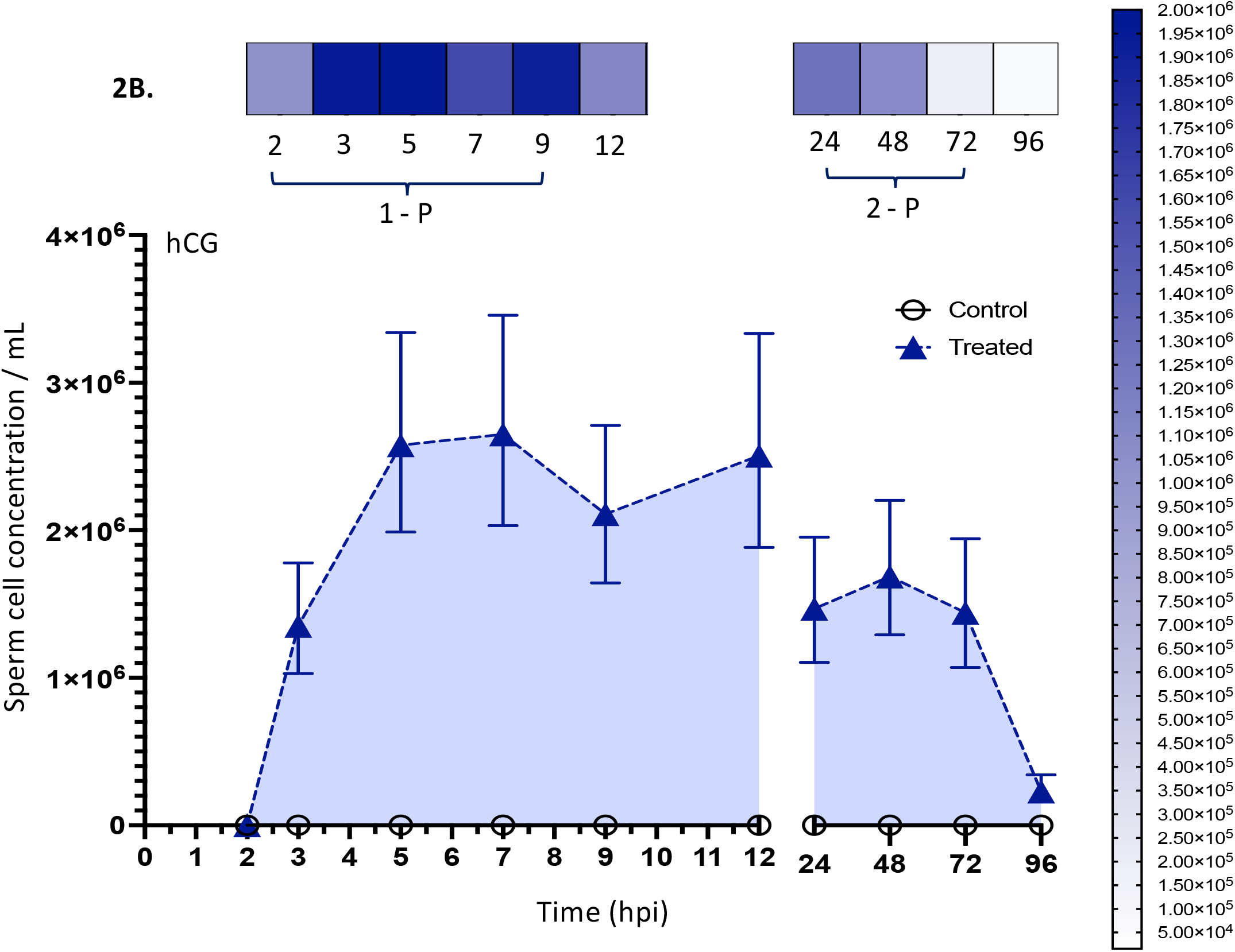
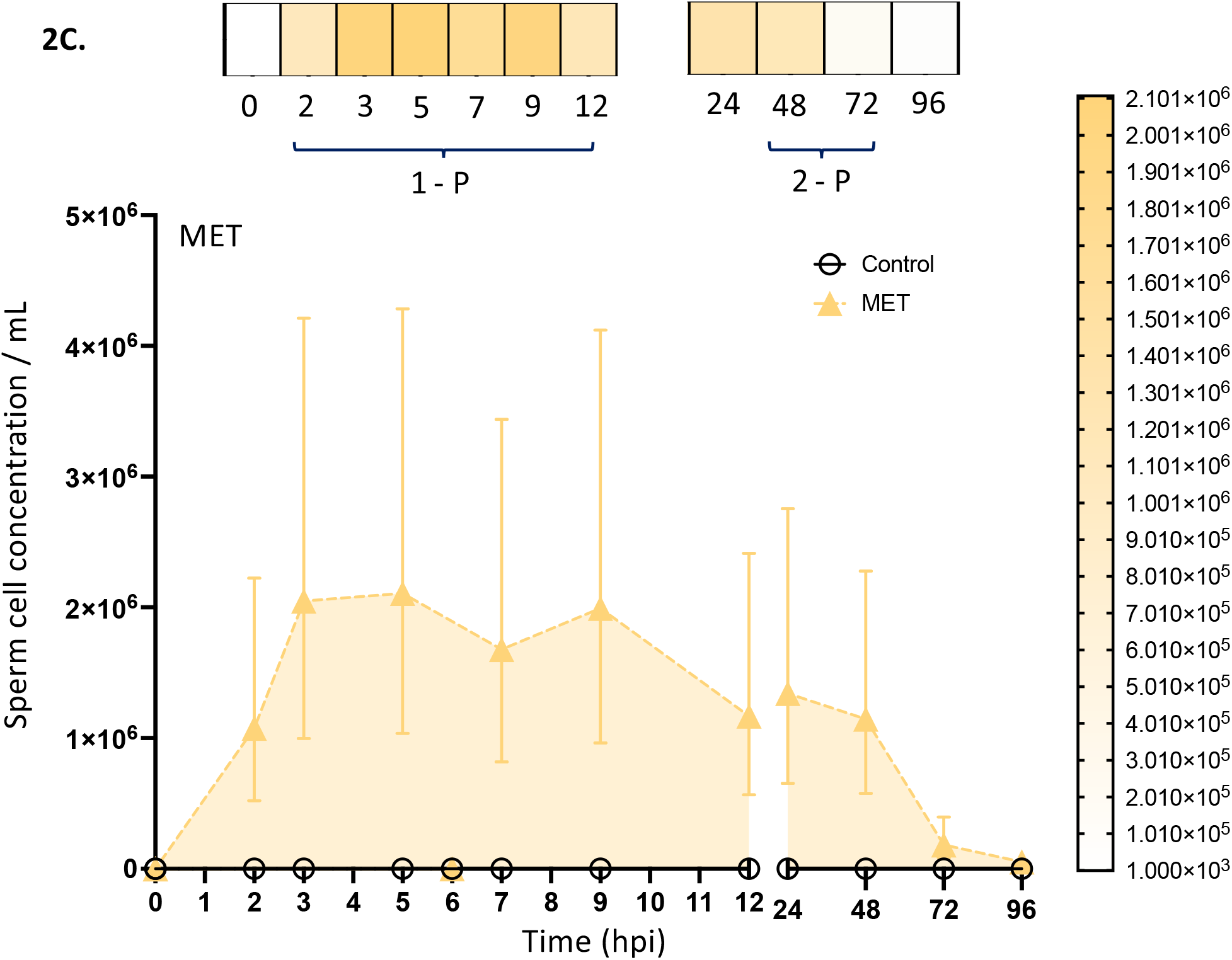
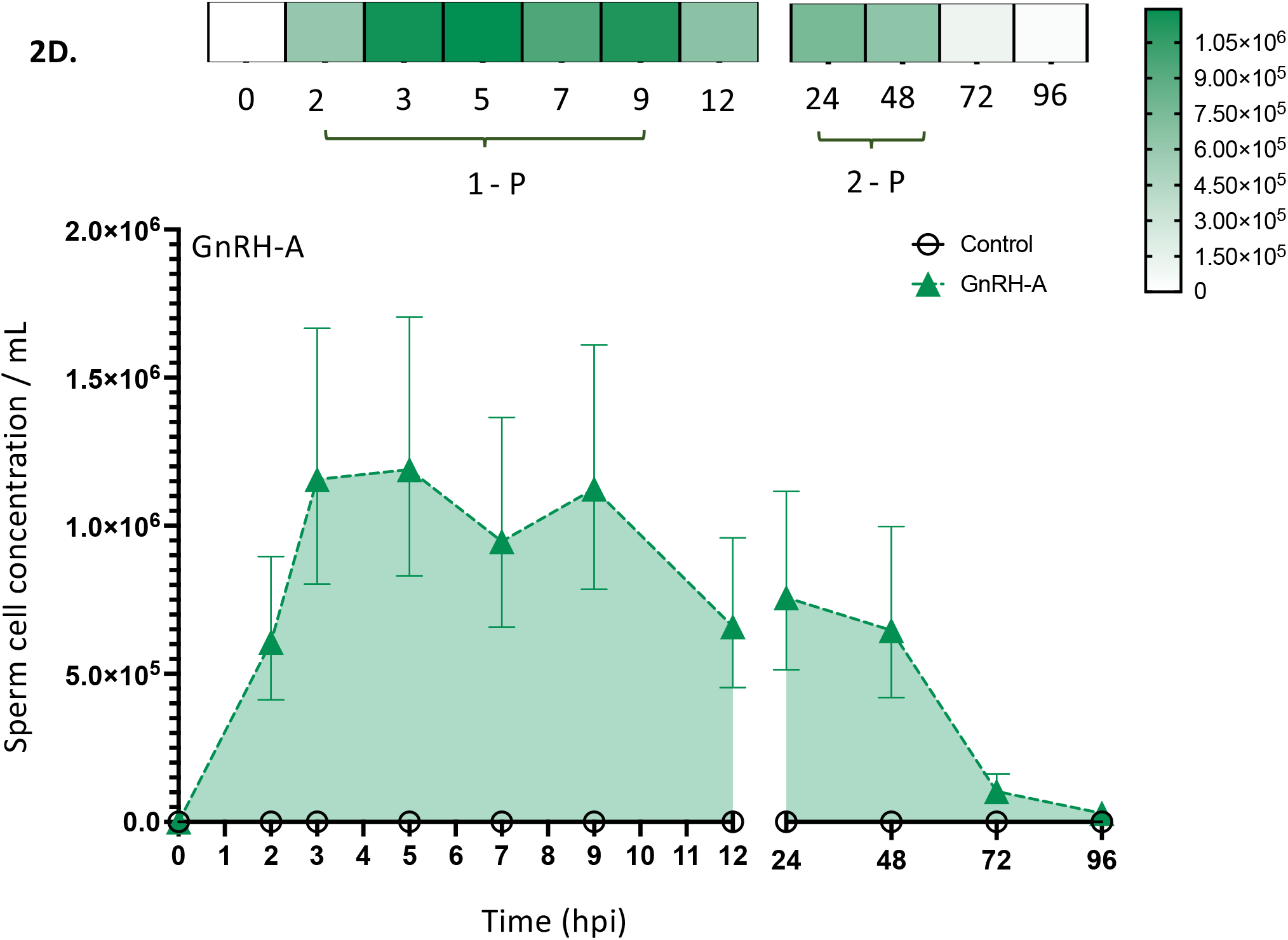
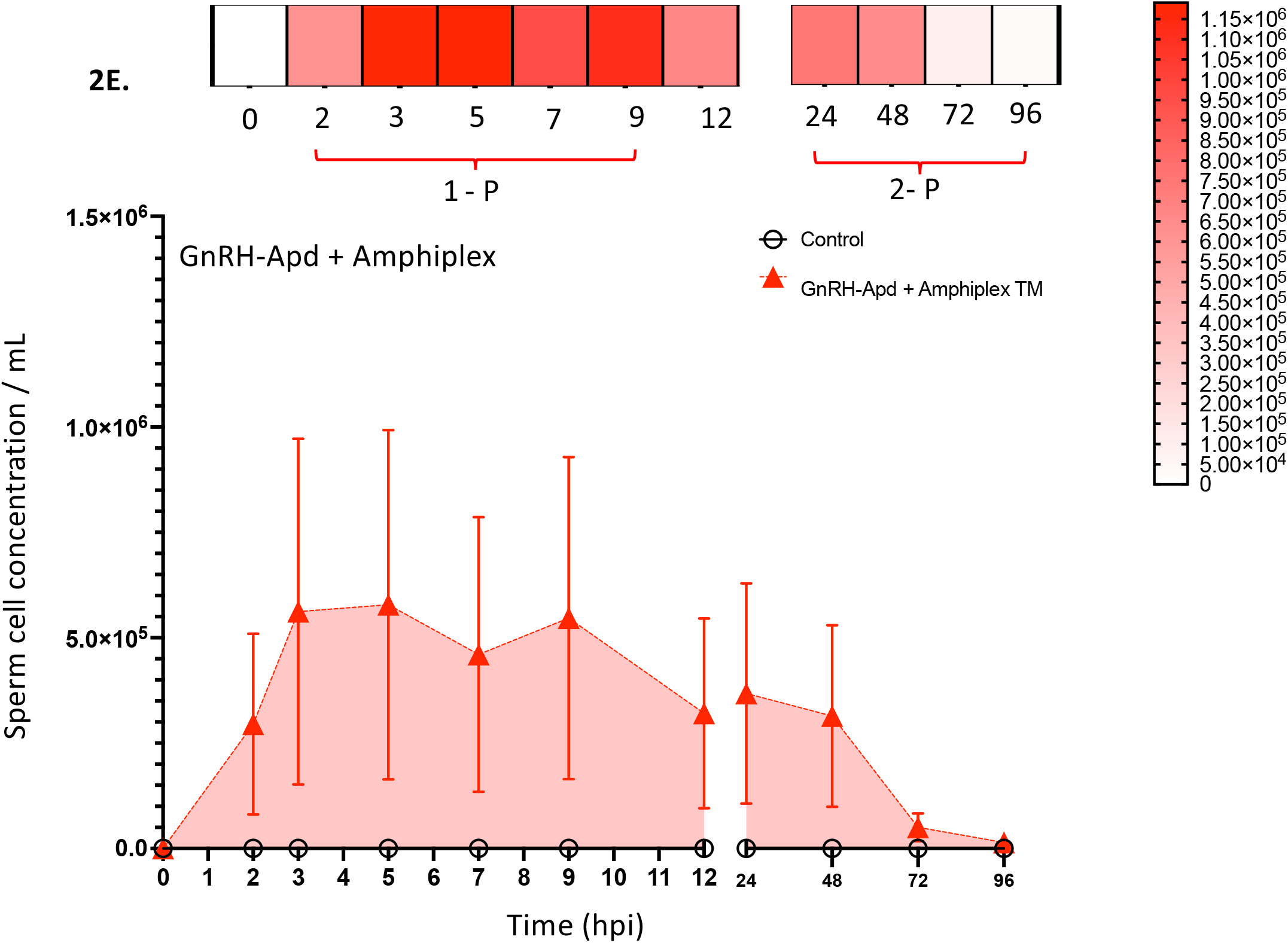
**A.** The response to hCG + GnRH-A led to a long release of spermatozoa with an initial peak of 7 h from, 3 hpi till 9 hpi. Later, at 24 hpi, an increase in sperm production occurred compared to 12 hpi signifying a probable second sperm peak occurring between these times and then steadily decreasing thereafter to 48 hpi. A steep drop in sperm production was observed at 72 – 96 hpi with the latter showing an almost complete return to zero. The greatest concentrations were noted at 5 and 9, 3 and 7 hpi respectively though relatively high concentrations, averaging 2.3 million cells were observed consistently across all time points with this treatment. Sperm concentrations are represented per milliliter with error bars representing 95% confidence intervals (CIs). **B.** The response to hCG led to an initial peak of 7 h starting at 3 hpi till 9 hpi as observed in the hCG + GnRH-A combination. The greatest concentrations were noted at 3, 5 and 9 hpi. Relatively high concentrations, (>2 million cells), were observed consistently across time with this treatment. A 24 hpi, an increase in sperm production occurred from 24 hpi till 48 hpi, decreasing thereafter. Sperm concentrations are represented per milliliter with error bars representing 95% confidence intervals (CIs). **C.** The response to MET led to least one peak that began at 3 hpi and lasted till 12 hpi. Total sperm concentrations exceeded >2 milion sperm for the first 10 h and remained relatively constant from 3 hpi till 9 hpi indicative of a long first 7 h peak. A second peak occurred from 24 hpi and from thereon concentrations declined from 48 hpi until reaching almost baseline at 96 hpi. Sperm concentrations are represented per milliliter with error bars representing 95% confidence intervals (CIs). **D.** The response to GnRH-A led induced at least one peak that lasted 10 hpi and a second one at 24 hpi. As with other treatments sperm concentrations declined from 48 hpi and reached almost zero by 96 hpi. Total sperm concentrations, >1million for the first 6 h declining from 9 to 12 hpi. As with other treatments, another peak was probable after 12 hpi with a decline from 24 hpi to 48 hpi. Sperm concentrations are represented per milliliter with error bars representing 95% confidence intervals (CIs). **E.** The response to Amphiplex led to at least one peak of 7 h from 3 to 9 hpi. Like GnRH-A, a second peak probably occurred, and sperm concentrations then declined from 24 - 48 hpi and reached almost zero by 96 hpi. Like most treatments, Amphiplex elicited sperm production for up to 72 hpi, though concentrations past 48 hpi were very low. Sperm concentrations are represented per milliliter with error bars representing 95% confidence intervals (CIs).

In frogs that responded, hormone treatments elicited spermiation responses at all time points collected, regardless of hormone treatment. Statistically, when averaged across all hormone treatments, sperm concentrations rose significantly and there was a significant almost two-fold increase in concentration when comparing 3 to 2 hpi (2-fold, OR 1.9, CI 1.44 −2.52). When comparing between 5 and 3 hpi there was no change (OR 1.0, CI 0.79 −1.3). There was a decrease in concentration when comparing 5 to 7 hpi (OR 1.3, CI 1.0 −1.6). Concentrations then followed a sinusoidal pattern, with sperm numbers increasing, though not significantly, from at 9 hpi compared to 7 hpi (OR 1.2, CI 0.898 - 1.56) but decreasing again, with significantly high sperm concentrations at 9hpi when compared to 12 hpi (OR 1.7, CI 1.287 - 2.261). In subsequent time-points, there was a small rise in concentration at 24 hpi but it was not significantly different in concentration to that observed at 12 hpi and for all subsequent time-points, concentrations decreased consistently until the final collection at 96 hpi with a significant 21.97-fold difference between 48 and 96 hpi (21.97-fold, SE±6.348, CI 15.845 - 41.72).

hCG + GnRH-A had the highest responses starting at 2 hpi (Estimated Marginal Mean [EMM] 2×10^6^, ^CI^ 1.5×10^6^ - 2.6×10^6^, Figure 2A), followed by hCG administered alone (2 hpi - EMMS 1.355×10^6^, CI 1.02×10^6^ - 1.78×10^6^, Figure 2B) and MET (2 hpi - EMMS 1.075×10^6^, CI 5.20×10^5^ - 2.22×10^6^, Figure 2C. While all treatments showed initial increases in concentrations, the above mentioned three treatments showed increasing sperm concentrations from 2 - 5 hpi that surpassed the millions (×10^6^), while other treatments such as GnRH-Apd + Amphiplex never surpassed concentrations of ×10^5^ and remained relatively stable from 3 - 9 hpi (∼5×10^5^) (Figure 2E). GnRH-A was slightly better than GnRH-Apd + Amphiplex at inducing sperm, reaching its highest concentrations early between 3 - 5 hpi (∼1.1×10^6^) (Figure 2D). All hormone treatments, remaining in the same order of efficacy, showed slight increases in concentrations at 24 and 48 hpi before declining substantially at 72 and 96 hpi (Figure 2A-E). The peaks in sperm concentration for each treatment were analyzed in greater detail and discussed below.

When combined, hCG + GnRH-A elicited the highest concentration of sperm above ∼2×10^6^ up to 48 hpi and remained the treatment with the highest sperm output up to 96 hpi (EMM 9.7×10^4^, CI 5.87×10^4^ - 1.60×10^5^). hCG followed by MET were the next best treatments, maintaining concentrations above 1×10^6^ until 48 hpi and dropping to ∼6.5 and ∼5.2×10^4^ by 96 hpi respectively. The hCG-GnRH-A combination was significantly better at inducing spermiation than all treatments. Compared to GnRH-Apd + Amphiplex, hCG-GnRH-A was 6.80-fold more efficient (SE±1.06, CI 5.00 - 9.21), 3.30-fold more efficient than GnRH-A alone (SE±1.356, CI 2.272 - 4.79), 1.86-fold and 1.26-fold more efficient than MET and hCG respectively (MET SE±0.66, CI 0.932 - 3.72; hCG SE±0.117, CI 1.27 - 1.73). (Appendix 3).

Sperm peaks were not analyzed statistically but were calculated to reflect the overall efficacy of the above-mentioned treatments. Sperm peaks occurred at similar times in treatments containing hCG, GnRH-A or a combination thereof, with a single long peak beginning at 3 hpi and continuing till 7 or 9 hpi (Figure 2A-C) and a subsequent peak observed at 24 hpi. Due to the lack of time points between 12 and 24 hpi, it is hard to determine whether an additional peak occurred between these two time points. GnRH-Apd + Amphiplex and MET also had two peaks and, though the concentrations were an order of magnitude lower, the first peak was longer than that observed for hCG + GnRH-A, hCG and GnRH-A (Figure 2D-E). As with hCG + GnRH-A, hCG and GnRH-A, a second peak was observed at 24 hpi.

### 3.2 Motility and viability

Sperm motility was measured as total and forward progressive motility (FPM) and ranged across treatments between 32 - 90%. We found significant effects of hormone treatment and timepoint (hpi) on total motility (LRT χ^2^(3) = 12.8, p=0.005 and LRT χ^2^(6) = 59.4, p = 5.872×10^-11^ respectively) and FPM (LRT χ^2^(3) = 25.3, p=1.366 x 10^-05^ and LRT χ^2^(6) = 59.3, p = 6.171×10^-11^ respectively).

Administration of hCG had the highest total motility and FPM and regardless of treatment, total motility and FPM decreased as time point of collection increased.

Average viability was high, with a range of 73 - 94 % live sperm. We found no significant effect of hormone treatments on viability (LRT χ^2^(3) = 1.9, p=0.6), however, there was a significant effect of collection time (hpi) (LRT χ^2^(6) = 24.6, p=0.0004), with motility decreasing after 12 hpi. The proportion of sperm collected from males injected with hCG + GnRH-A (deemed the best treatment), had a mean number of intact acrosomes of 60% (SEM ±2%).

### 3.3 Total motility

Overall, sperm collected by hCG administered singularly had the highest proportion of motile sperm, compared to hCG + GnRH-A (1.279-fold; CI 1.006 - 1.625), GnRH-A + GnRH-Apd + Amphiplex (1.358-fold; CI 0.871 - 2.118) and GnRH-A (2.198-fold; CI 1.376 - 3.511). hCG + GnRH-A was the next most effective treatment compared to GnRH-Apd + Amphiplex (1.062-fold, CI 0.691 - 1.632) and GnRH-A (1.718-fold, ; CI 0.1063 - 2.777). GnRH-Apd + Amphiplex had a higher proportion of GnRH-A alone (1.618-fold; CI 0.883 - 2.965). (Appendix 4 - 5.

Total sperm motility was significantly greater during the first 12 hpi compared to 24 hpi irrespective of hormone treatment. From the time of the first collection, a gradual increase in motility was noted up to 9 hpi. The highest proportions of motile sperm were detected between 7 and 9 hpi and were higher than those recorded at to 2, 3, 5 and 12 hpi. By 24 hpi motility had significantly decreased compared to all other time points (Figure 3, Appendix 6).

**Figure 3.**
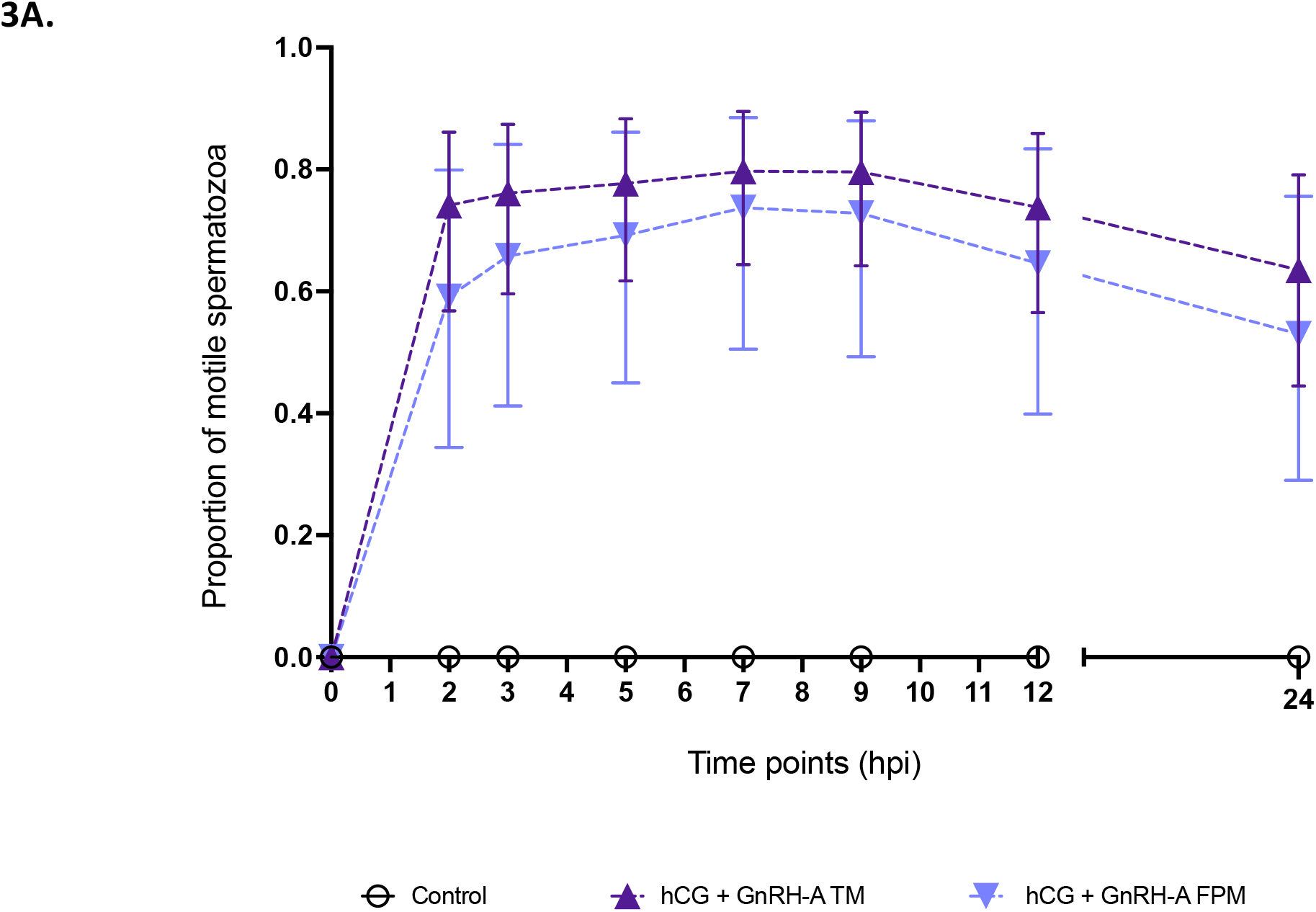
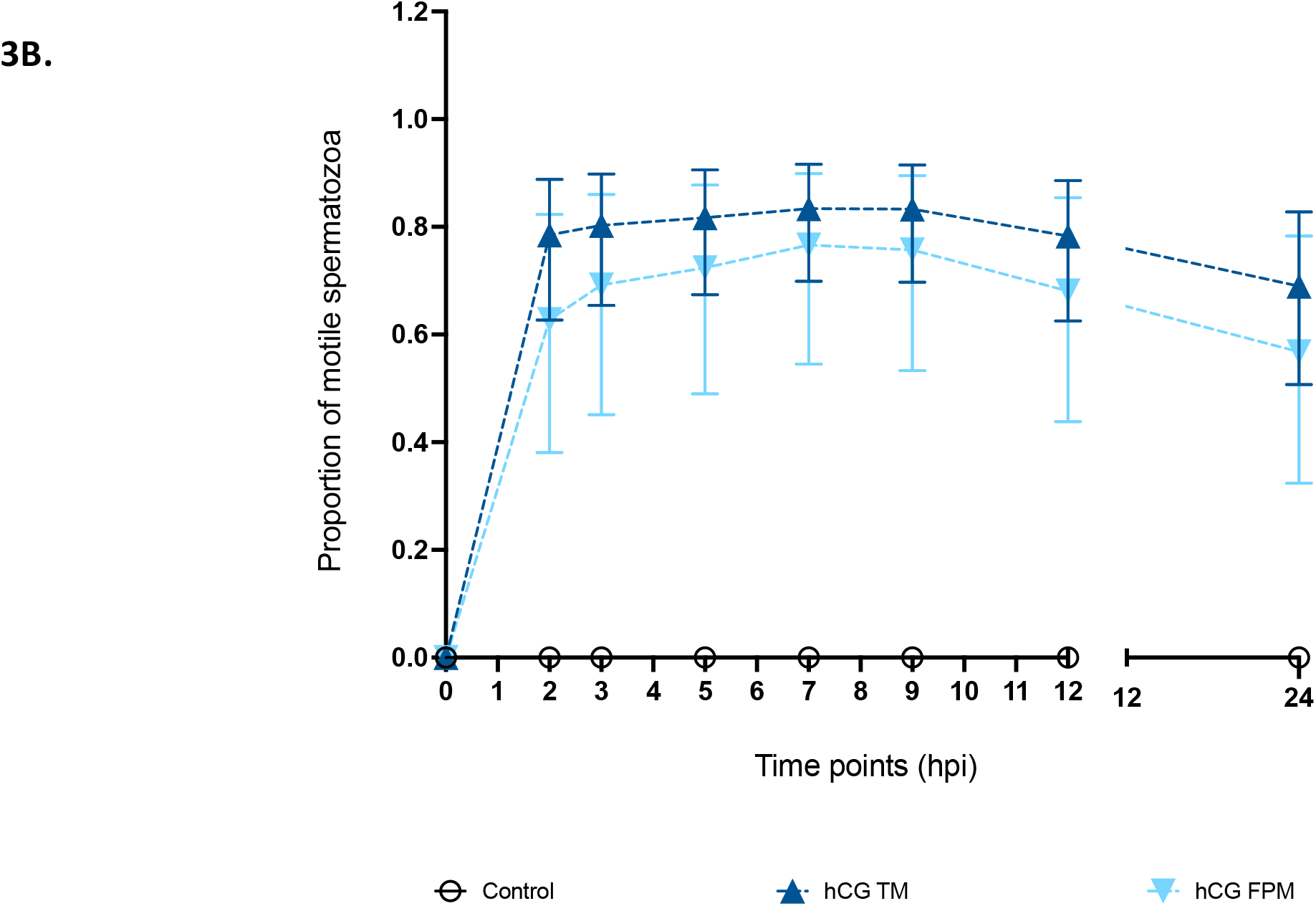
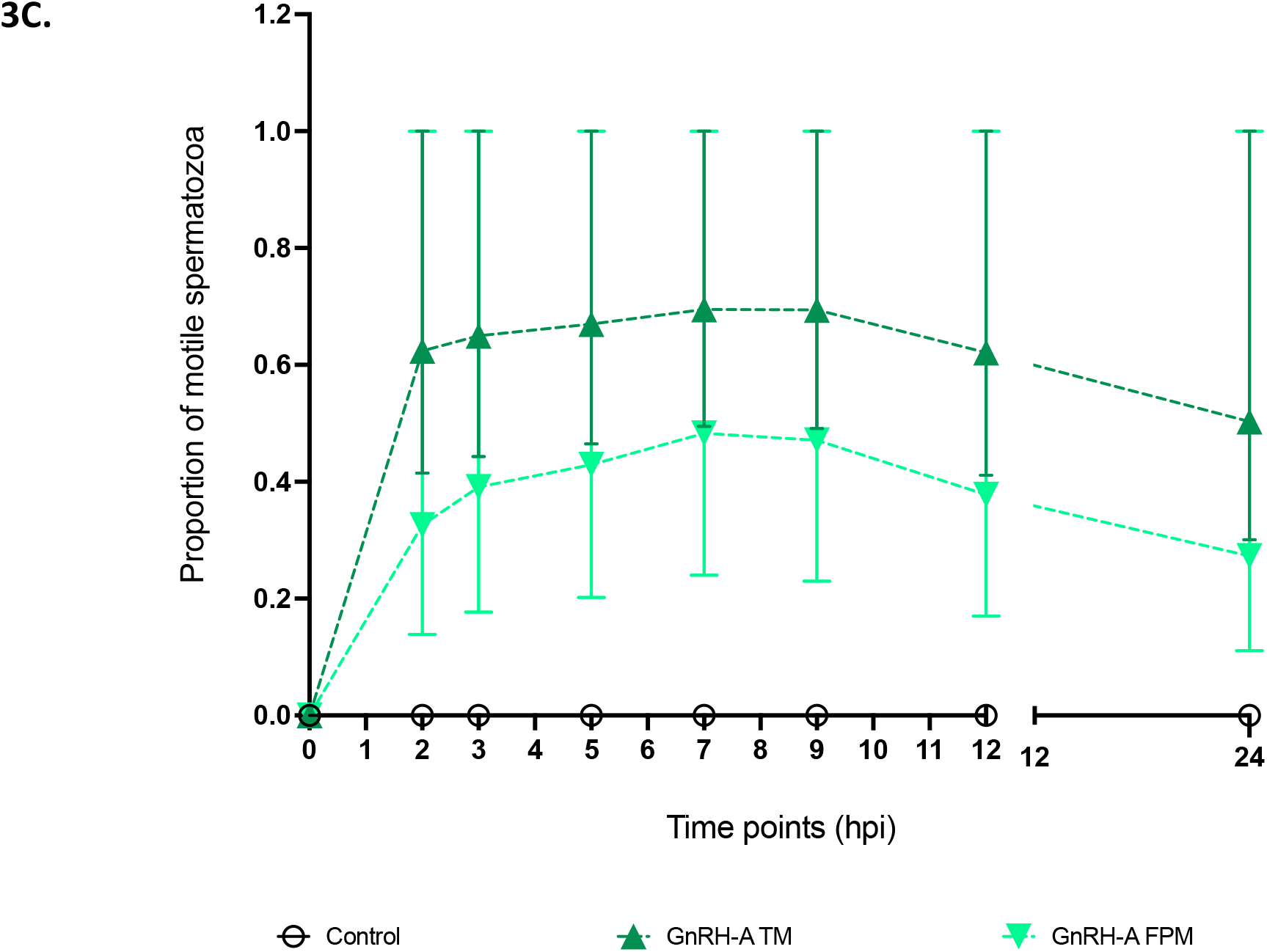
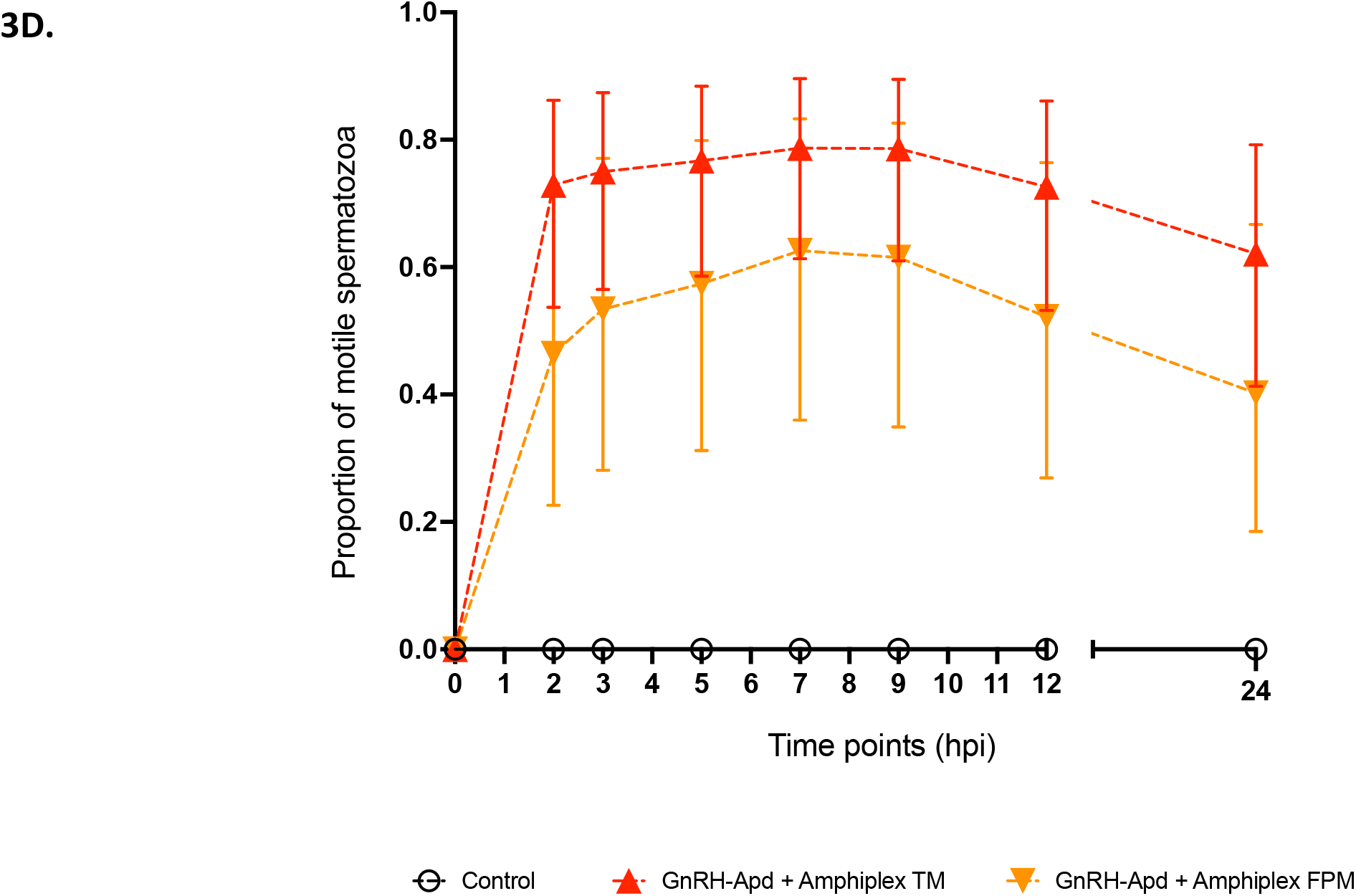
**A.** Points represent time points collected over a 24 h period for hCG + GnRH-A. TM was high between ∼70 - 80% while FPM was also quite high at ∼60 - 75%. The proportion of total spermatozoa (TM, darker upright triangles) and forward progressive motile (FPM, lighter inverted triangles) plotted as estimated marginal means (EMMS) with error bars representing 95% confidence intervals (CIs). **B.** Points represent time points collected over a 24 h period for hCG. TM was high between ∼75 - 80% while FPM was also quite high at ∼60 - 75%. The proportion of total spermatozoa (TM, darker upright triangles) and forward progressive motile (FPM, lighter inverted triangles) plotted as estimated marginal means (EMMS) with error bars representing 95% confidence intervals (CIs). **C.** Points represent time points collected over a 24 h period for GnRH-A. TM was lower than hCG-based treatments, between ∼60 −65% while FPM was also lower at ∼25 - 40%. The proportion of total spermatozoa (TM, darker upright triangles) and forward progressive motile (FPM, lighter inverted triangles) plotted as estimated marginal means (EMMS) with error bars representing 95% confidence intervals (CIs). **D.** Points represent time points collected over a 24 h period for GnRH-Apd + Amphiplex TM was high between ∼75 - 80% while FPM was also quite high at ∼45 - 60%. The proportion of total spermatozoa (TM, darker upright triangles) and forward progressive motile (FPM, lighter inverted triangles) plotted as estimated marginal means (EMMS) with error bars representing 95% confidence intervals (CIs).

### 3.4 Forward progressive motility

As with overall motility, hCG and hCG + GnRH-A, had the highest proportion of motile sperm but were not significantly different to each other. hCG had a higher proportion of FPM compared to GnRH-A, (3.5-fold, SE±0.959; CI 2.044 - 5.99) and GnRH-Apd + Amphiplex (1.954-fold; CI 1.167-3.27). The combination of hCG + GnRH-A had a significantly higher FPM than GnRH-Apd + Amphiplex (2.997-fold; CI 1.012-2.77) and GnRH-A (1.674-fold; CI 1.722-5.22) respectively. (Appendix 7 - 8).

Across time points, significant differences in FPM were noted between time points in the first 12 hpi. Specifically, a steady but insignificant increase at 5 compared to 3 hpi (OR: 1.2, CI 0.9 - 1.5). Increases in fpm at 7 hpi compared to 3 hpi were significant (OR: 1.5; CI 1.112-1.914). Although proportions continued to increase from 3 to 5 until 7 hpi, differences were not significant between 5 and 7 hpi nor was there any statistical difference between 7 and 9 hpi though the proportions of FPM sperm began to decrease by 9 hpi. By 12 hpi there was a lower likelihood of detecting FPM sperm than at 7 (1.535-fold; CI 1.168-2.018) and 9 hpi (1.465-fold; CI 1.112-1.929) respectively. By 12 and 24 hpi, the proportion of FPM sperm in the sample was similar to that detected at the first time of collection (2 hpi). Therefore, by 24 hpi, FPM sperm had decreased significantly when compared to 3 (OR: 1.7; CI 1.29-2.252), 5 (OR: 2.0; CI 1.514-2.637), 7 (OR:2.5; CI1.879 −3.298) and 9 h pi (OR: 2.4; CI 1.787-3.152) respectively (Figure 3 A-D). (Appendix 9).

### 3.5 Sperm viability

Overall, viability was relatively high throughout the duration of the study with a range of 82 – 89% across all treatments in the first 12 h pi (Figure 4A-D). Unlike the other parameters tested, the only influential factor acting on the proportion of live sperm was time of collection (hpi). No significant differences in the proportion of live sperm were detected between time points in the first 12 hpi. It was only when earlier timepoints (2,3,5,7,9,12) were compared to 24 hpi that significant differences could be observed with ORs ranging from 1.5-1.7 (Appendix 10 - 12). (See Figure 5A. for live/dead eosin-nigrosin staining of boreal toad spermatozoa).

**Figure 4.**
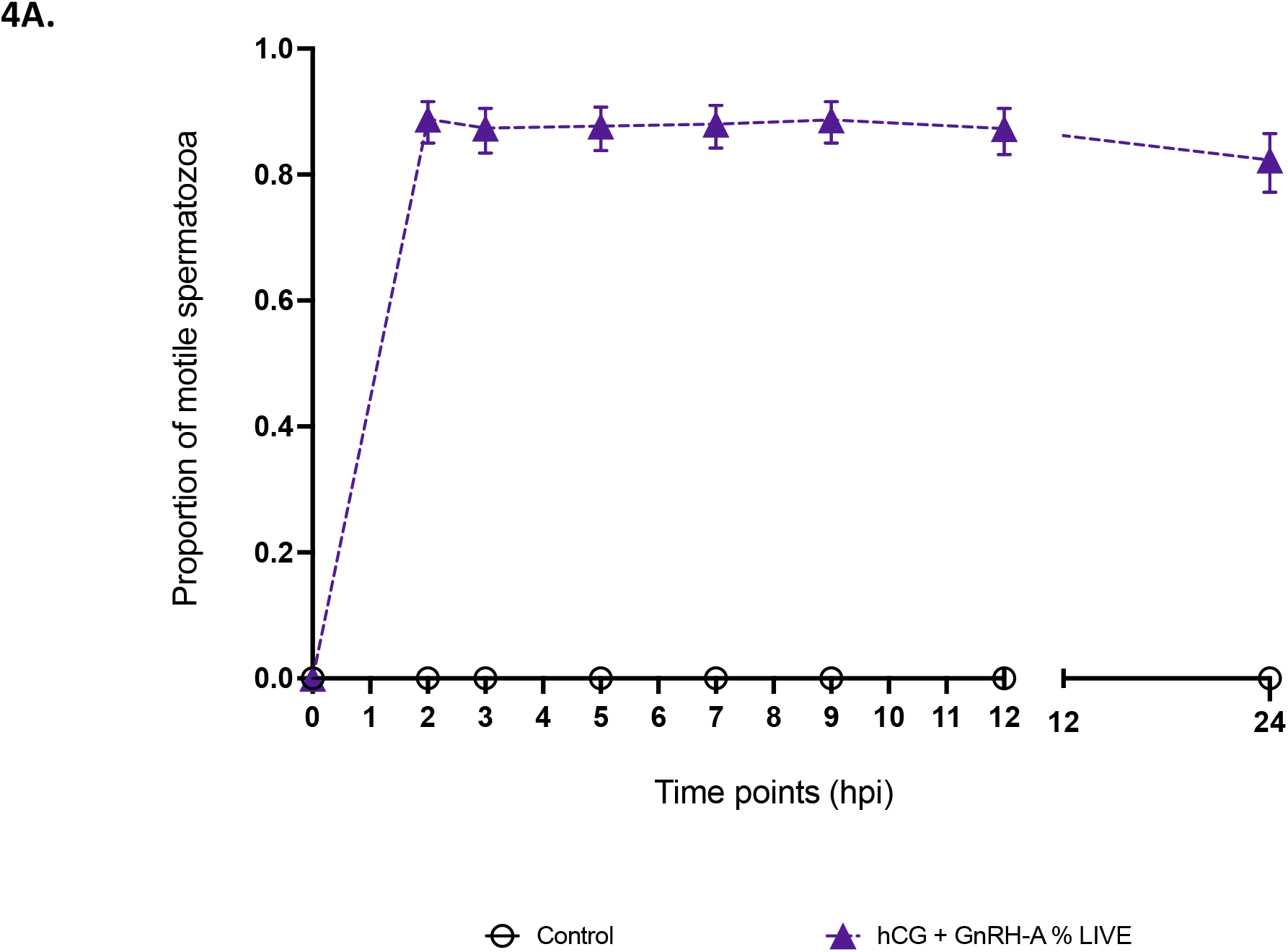
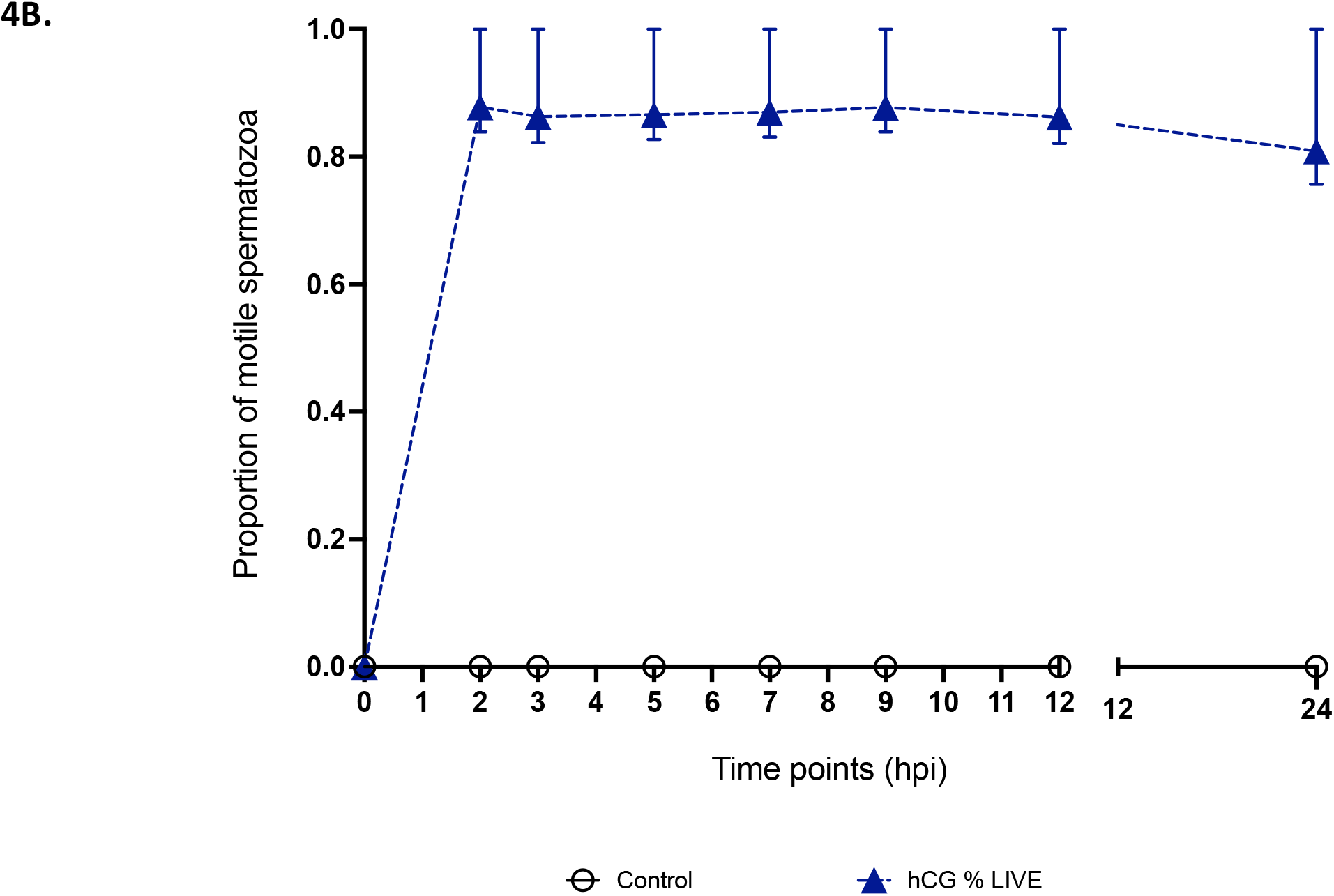
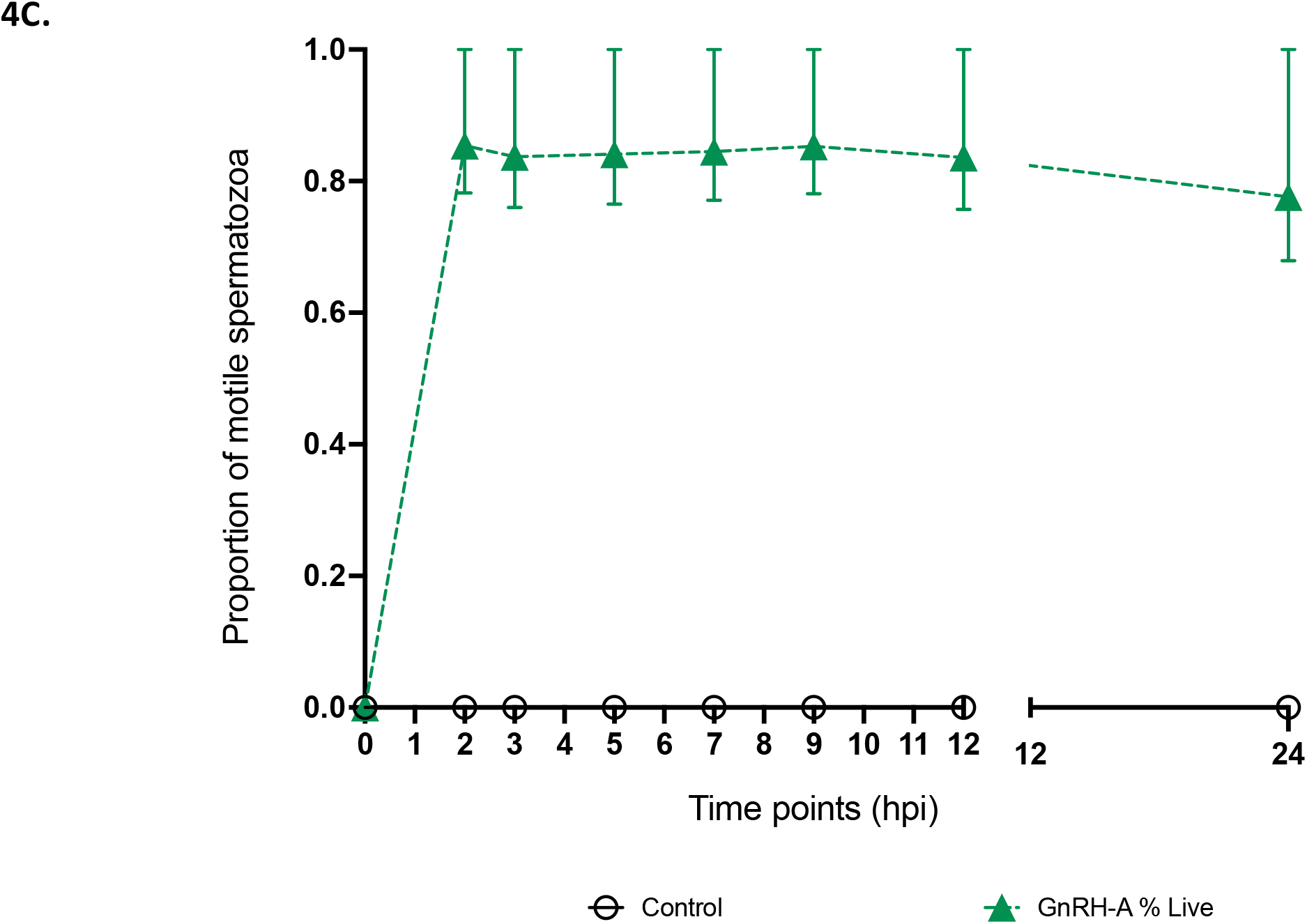
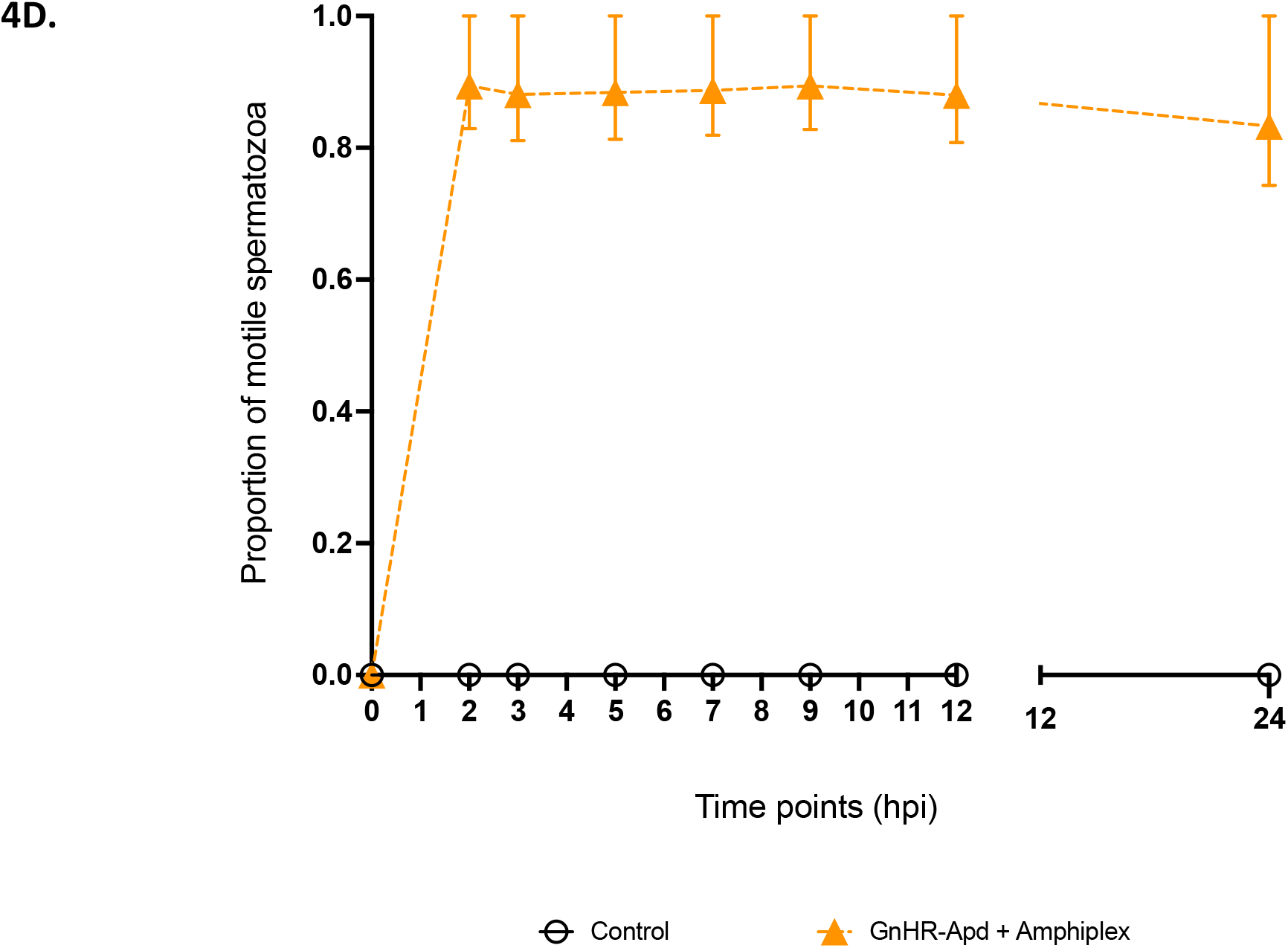
**A.** Vitality staining with eosin-nigrosin indicated that the proportion of live sperm cells in hCG + GnRH-A were not significantly different with all-time points yielding above ∼80% live cells during the first 24 hpi or significantly different to those observed in the other hormone treatments. The proportion of live cells plotted as estimated marginal means (EMMS) with error bars representing 95% confidence intervals (CIs). **B.** Vitality staining with eosin-nigrosin inidcated that the proportion of live sperm cells in hCG were not significantly different with all-time points yielding above ∼80% live cells during the first 24 hpi or significantly different to those observed in the other hormone treatments. The proportion of live cells plotted as estimated marginal means (EMMS) with error bars representing 95% confidence intervals (CIs). **C.** Vitality staining with eosin-nigrosin inidcated that the proportion of live sperm cells in GnRH-A were not significantly different with all-time points yielding above ∼80% live cells during the first 24 hpi or significantly different to those observed in the other hormone treatments. The proportion of live cells plotted as estimated marginal means (EMMS) with error bars representing 95% confidence intervals (CIs). **D.** Vitality staining with eosin/nigrosin inidcated that the proportion of live sperm cells in GnRH-Apd + Amphiplex were not significantly different with all-time points yielding above ∼80% live cells during the first 24 hpi or significantly different to those observed in the other hormone treatments. The proportion of live cells plotted as estimated marginal means (EMMS) with error bars representing 95% confidence intervals (CIs).

**Figure 5.**
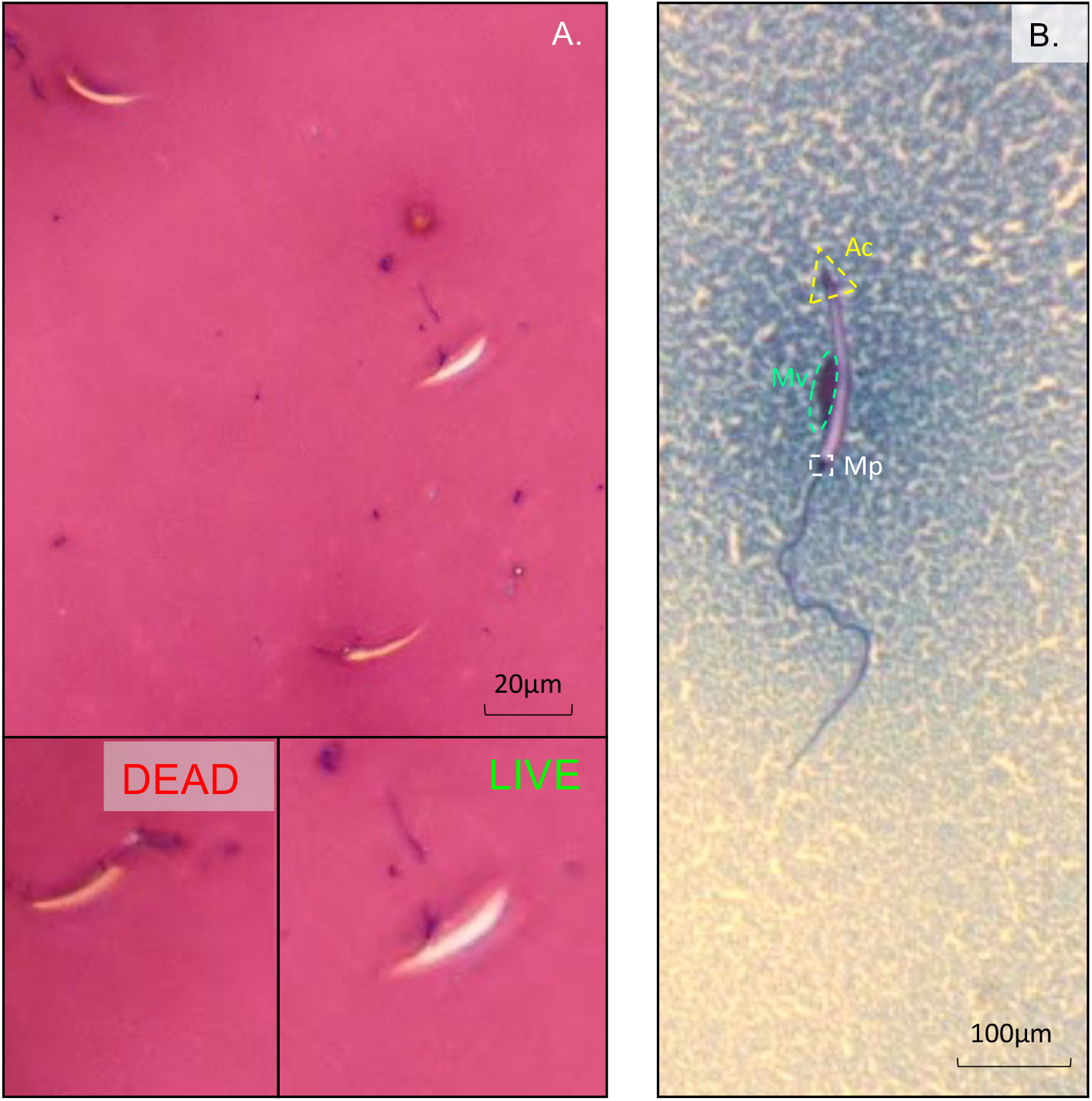
**A)** Eosin-nigrosine staining of boreal toad sperm with both live and dead spermatozoa. Staining was performed on sperm acquired from all hormone treatments tested. **B)** Coomasie blue staining the mitochondrial vesicle and intact acrosome of boreal toad sperm. Coomasie staining was only performed on sperm recovered from hCG – GnRH-A treatments.

### 3.6 Acrosome integrity

Due to logistical constraints, acrosome integrity was only analyzed in the hCG + GnRH-A since this was determined to be the best treatment out of the four tested (Figure 5B). Results indicate an average acrosome integrity of 60 % (SD ± 10.31) in this treatment. Notably, we also observed that integrity in this treatment group was different across different populations (manuscript in preparation), suggesting that fertilization capabilities may be variable across genetic populations and generations within those populations. These preliminary results have management impacts for future breeding efforts. Future research should compare this parameter across all treatments to elucidate the relationship between hormone treatment and acrosome integrity thereby adding to sperm viability scores but also developing *ex situ* breeding strategies to understand and improve quality not just quantity.

### 3.7 Effects of hormone and month on sperm concentration

Both the type of hormone treatment (LRT χ^2^(3) = 83.1, p<2.2×10^-16^) and the month in which it was administered (LRT χ^2^(3) = 55.5, p=5.5×10^-12^) affected sperm concentration significantly (Figure 6A). Irrespective of hormone type, when analyzing hormone efficacy across months overall decreases were observed. Odds Ratios (OR) were used to further compare results between treatments. Higher concentrations were obtained in June and decreased thereafter following a downward trend to July (1.4-fold, OR 1.66; CI 1.257 - 2,191), August (3-fold, OR 3.119; CI 2.324 - 4,184) and until the last collections in September (2-fold, OR 1.917; CI 1.322 - 2.781). July was the next best month in which to obtain higher concentrations of sperm compared to August (OR 1.879; CI 1.451 - 2.434) and September (OR 1.155; CI 0.836 - 1.596) (Figure 7A).

**Figure 6.**
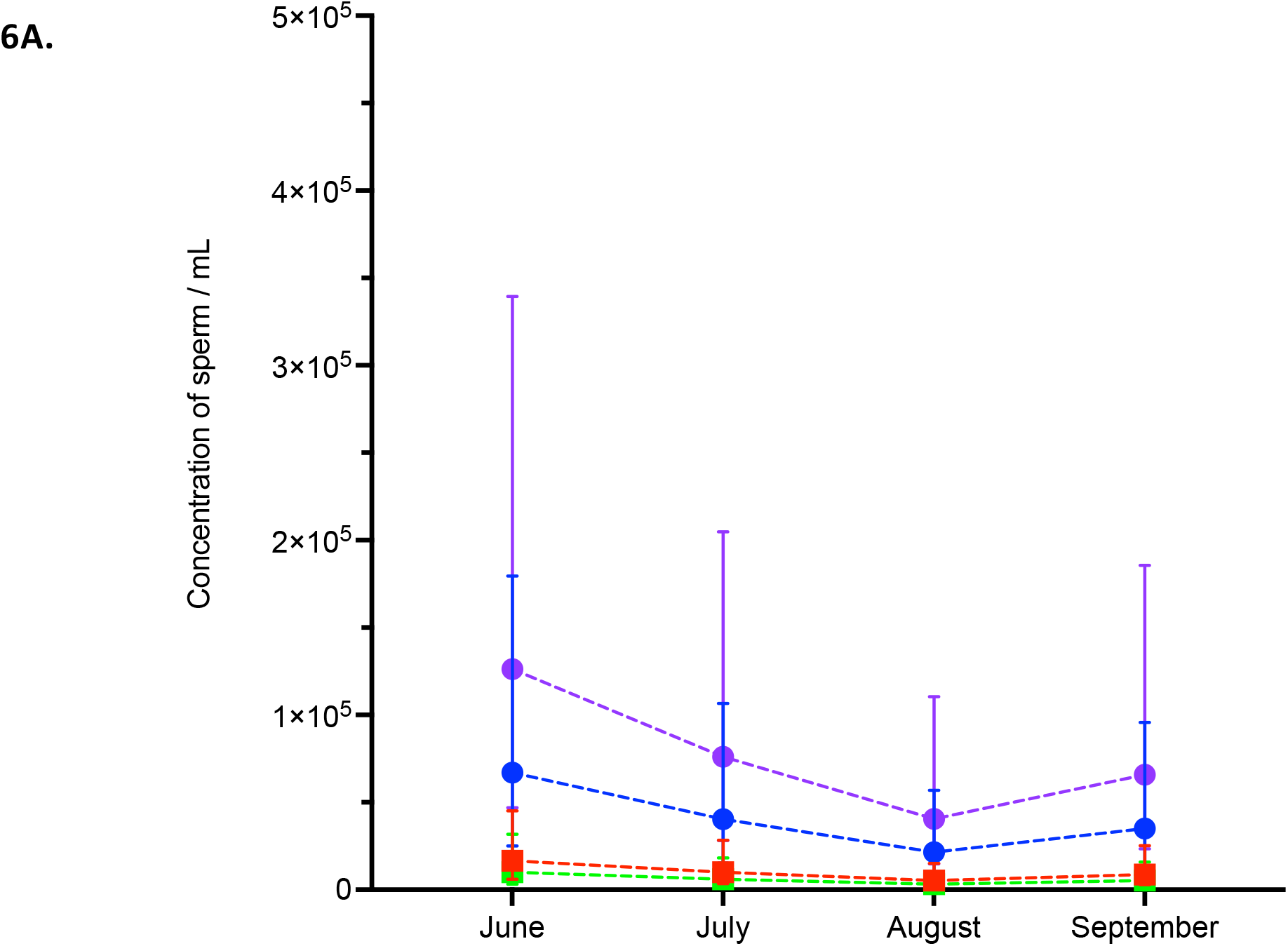
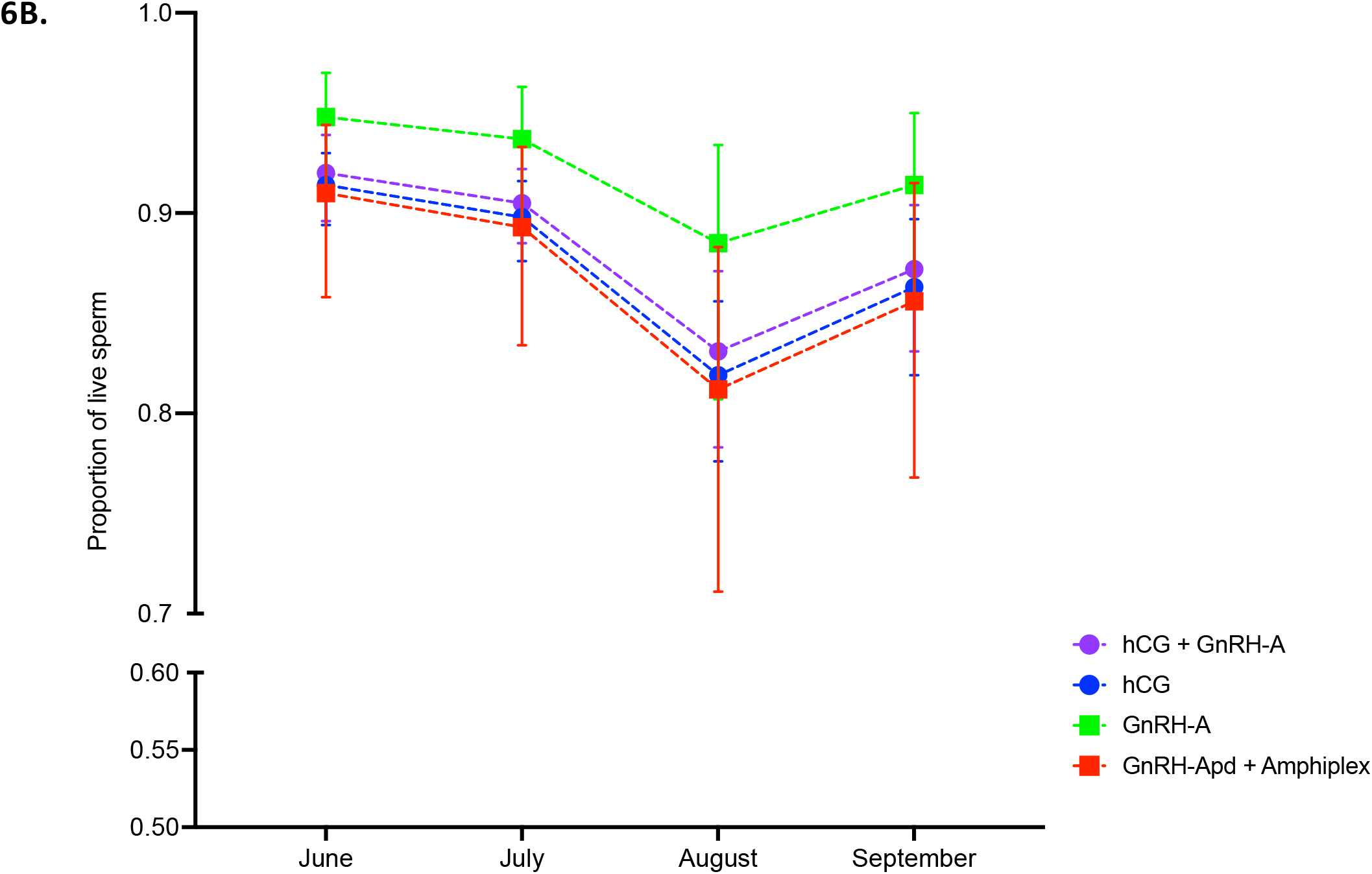
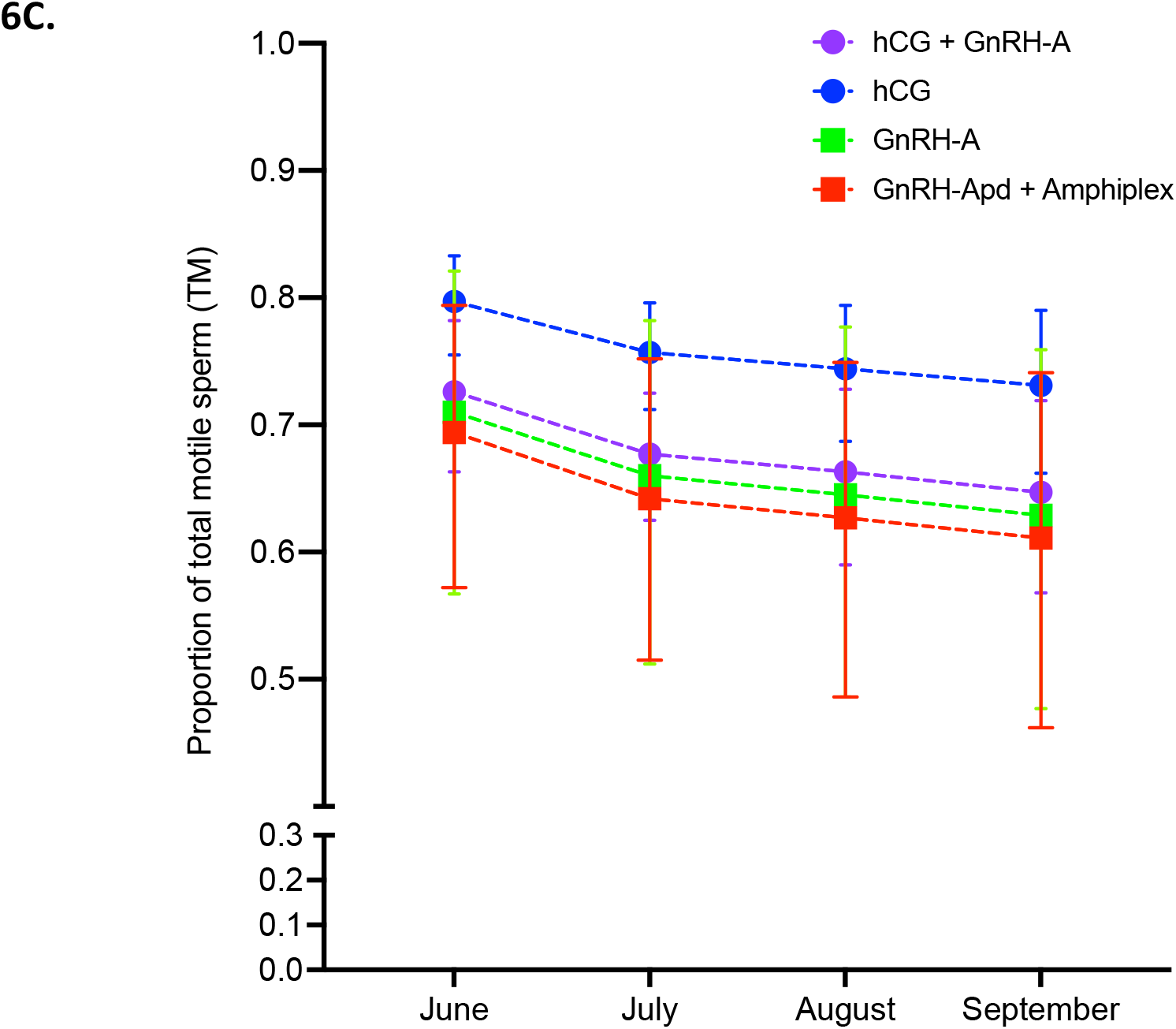
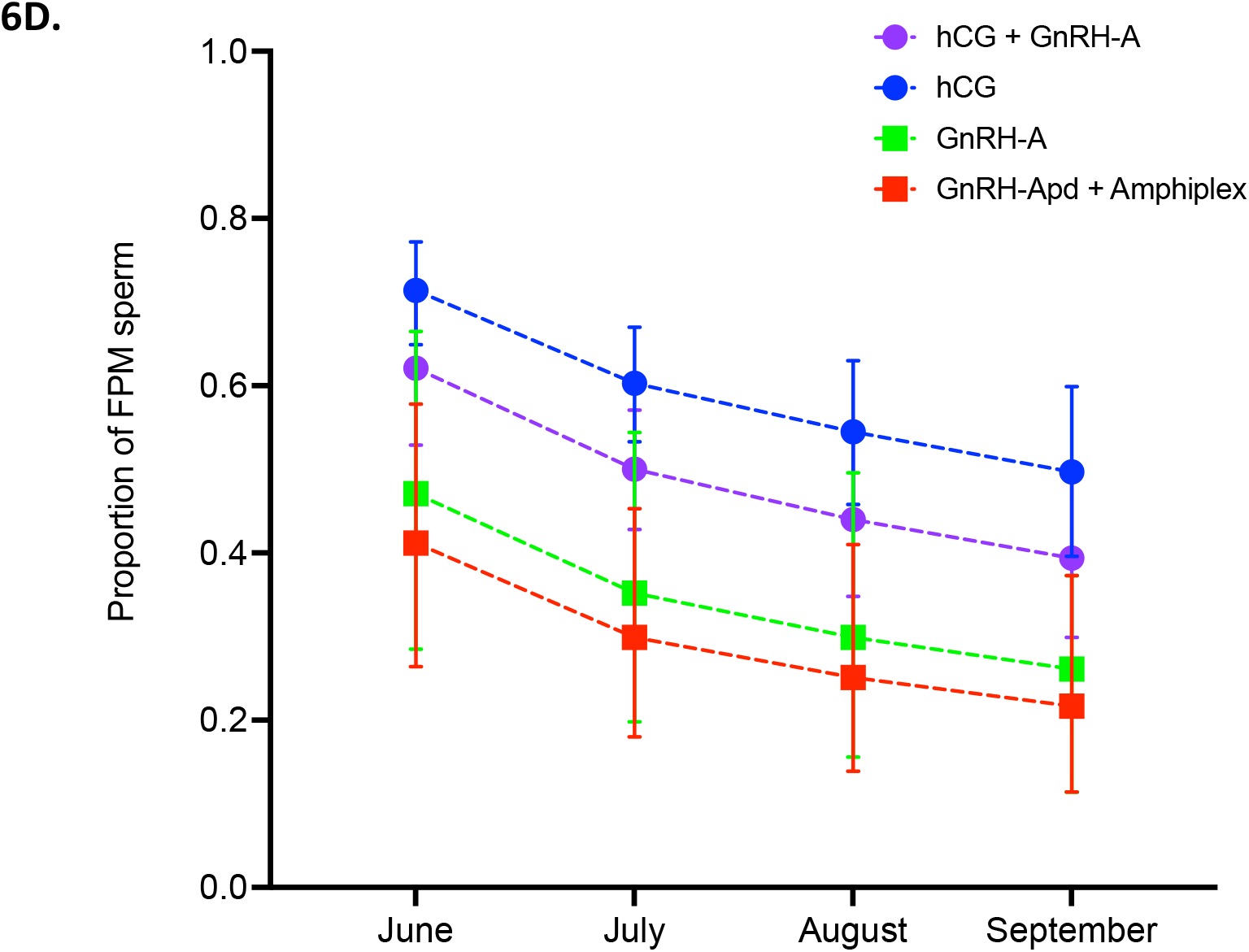
**A.** The proportion of live cells across treatments for June, July, August and September 2014-15. Proportion of hormones plotted as estimated marginal means (EMMS) with error bars representing 95% confidence intervals (CIs). **B.** The proportion of live cells across treatments for June, July, August and September 2014-15. Proportion of live sperm plotted as estimated marginal means (EMMS) with error bars representing 95% confidence intervals (CIs). **C.** The proportion of total motile cells across treatments for June, July, August and September. Proportion of total motility was not significantly affected by the month of administration. TM plotted as estimated marginal means (EMMS) with error bars representing 95% confidence intervals (CIs). **D.** The proportion of FMP cells treatments for June, July, August and September. FMP was significantly affected by the month and the hormone treatment. Samples are plotted as estimated marginal means (EMMS) with error bars representing 95% confidence intervals (CIs).

**Figure 7.**
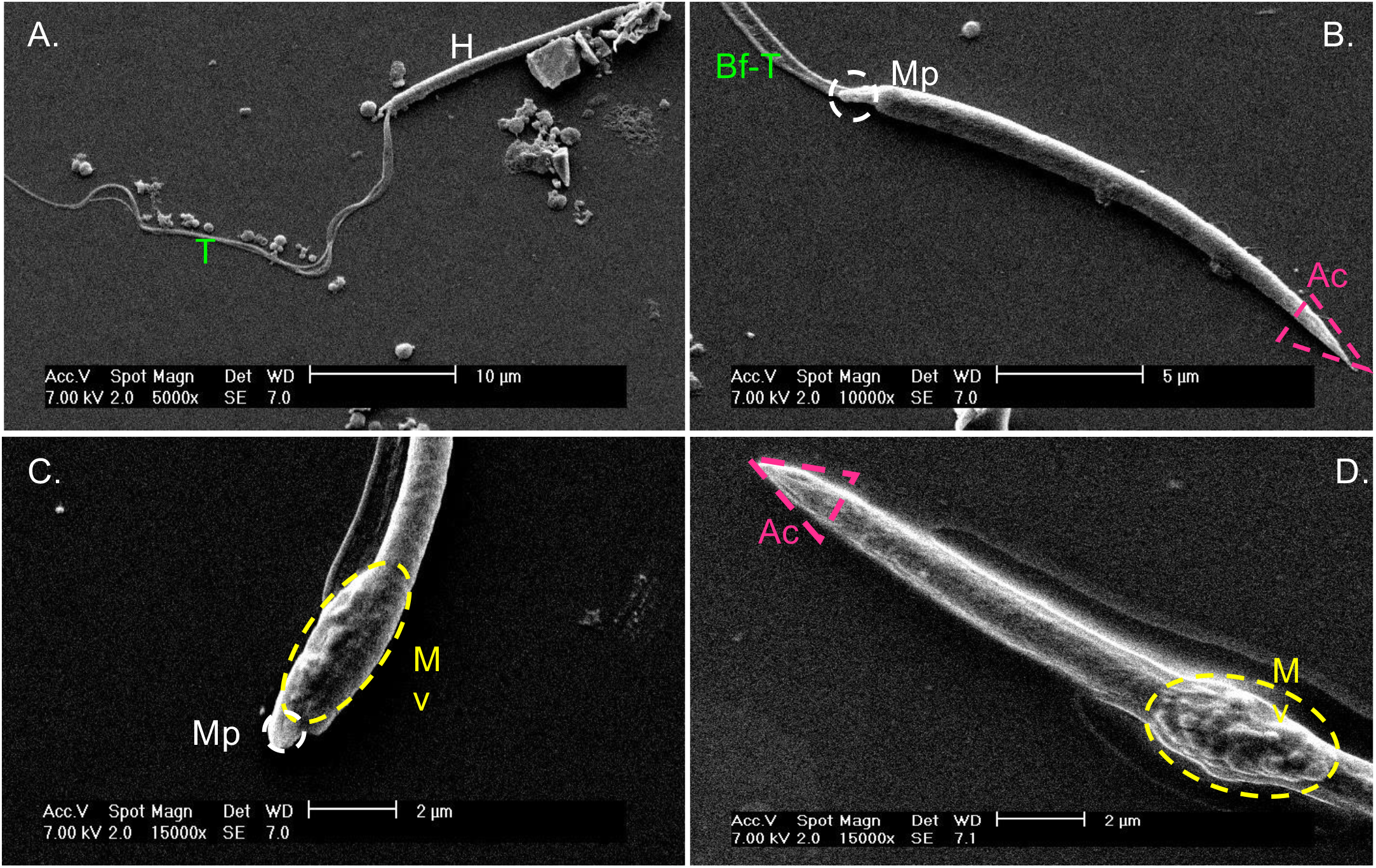
Scanning Electron Microscopy of *Anaxyrus boreas boreas* sperm collected after hormonal induction. **A-B)** Spermatozoa without mitochondrial vesicle; **C)** Spermatozoa with mitochondrial vesicle. H = head structure with additional structures as shown on, B - D: Ac - acrosome, Mitochondrial vesicle - MV, Mp = mid- piece (mitochondrial collar) highlighted in B; T = Tail with additional denotation as shown in FA - B: Bf-T - bi-flagellar tail.

When analyzed across months, the hCG-based treatments outperformed treatments containing GnRH-A with respect to concentration. Once again, hCG + GnRH-A was the best across all the months examined compared to GnRH-A (12-fold, OR 12.61, CI 5.005 - 31.77), GnRH-Apd + Amphiplex (7-fold, OR 7.62; CI 4.549 - 12.76) and hCG (2-fold, OR1.88; CI 1.328 - 2.67). Administered singly, hCG was the second-best treatment compared to GnRH-A (7-fold, OR 6.68; CI 2.84 - 15.78) and GnRH-Apd + Amphiplex (4-fold, OR 4.045; CI 2.359 - 6.94) (Figure 7A).

### 3.8 Effects of hormone and month on sperm quality

Sperm motility was measured as total and forward progressive motility (FPM) and ranged across treatments between 61 - 80% for total motility and 22 - 71% for FPM (Figure 6B and 6C). We found significant effects of hormone treatment but not month on total motility (LRT χ^2^(3) = 12.1, p=0.007 and LRT ^2^(3) = 4.7, p = 0.2 respectively), with treatments administered in June yielding the highest proportion of total motility. We found a significant effect of both hormone and month on FPM (LRT χ (3) = 21.3, p=9.3 x 10 and LRT χ (3) = 17.7, p = 0.0005 respectively), with hCG administered in June yielding highest FPM. Average viability was high, with a range of 81 - 95% live sperm (Figure 6C). We found no significant effect of hormone treatments on viability (LRT χ^2^(3) = 3.6, p=0.3), however, there was a significant effect of month (LRT χ^2^(3) = 27.7, p=4.2×10^-6^), with viability of sperm collected in June being highest (Figure 6D, Appendix 13).

For total motility, regardless of month administered, we found hCG administered alone was the significantly better treatment compared to hCG-GnRH-A (OR 1.5, 95% CIs 1.1-1.9) and GnRH-Apd + Amphiplex (OR 1.7, 95% CIs 1.0-2.9) and was not significantly different to GnRH-A alone (OR 1.6, 95% CIs 0.9-3.0). There were no other significant differences, however hCG-GnRH-A had a higher total motility than GnRH-Apd + Amphiplex (OR 1.2, 95% CIs 0.7-2.0) (Figure 6B, Appendix 14).

For collecting sperm with FPM, we found hCG administered alone was the best treatment compared to hCG + GnRH-A (1.5-fold, OR 1.523; CI 1.106 - 2.097), GnRH-A (2-fold, OR1.842; CI 1.277 - 6.167), GnRH-Apd + Amphiplex (3-fold, OR 3.57; CI 1.819 - 6.167). Overall, June was the best month with significantly greater FPM than compared to sperm collected in July (2-fold, OR1.643; CI 1.128 - 2.39), August (2-fold, OR2.086; CI 1.343 - 3.24) and September (2-fold, OR2.529; CI 1.563 - 4.092). Results showed an interesting trend in the hormones across months. Though not significant due to low replication, particularly in the case of GnRH-A and GnRH-Apd + Amphiplex, in August and September, hCG alone appeared to maintain the highest proportion of FMP sperm followed by hCG + GnRH-A. (Figure 6C, Appendix 15).

For collecting viable sperm, regardless of hormone administered, June had a greater proportion of live sperm than either August (2-fold, OR2.349; CI 1.671 - 3.301) or September (2-fold, OR1.696; CI 1.165 - 2.471) but not July (1-fold, OR 1.209; CI 0.91 - 1.608). July had a greater proportion of live sperm than August (2-fold, OR1.942; CI 1.414 - 2.667) but no significant differences were noted between August and September (Figure 6D, Appendix 16 -18).

### 3.9 Sperm morphology

With scanning electron microscopy (SEM), we examined the ultrastructure of sperm released post-hormone induction. Averaged from five photographed sperm, the head of boreal toad sperm measured approximately 18.96 ± SD 1.97 μm and its midpiece about 1.61 ± SD 1.14 μm midpiece (Figure 8A-B). Tail measurements showed 30.24 ± SD 12.79 μm. As with many anurans, the tail was an undulated membrane with paraxonemal rods attached to a filiform head as described by Van Der Horst (2022) (Figure 8A) [17,23,43]. During hormonal induction spermatozoa without (Figure 1A-B) and with mitochondrial vesicles were collected (Figure 7C-D). The mitochondrial vesicle was composed of an oval, granular mass on the central side of the head (Figure 7C-D).

**Figure 8.**
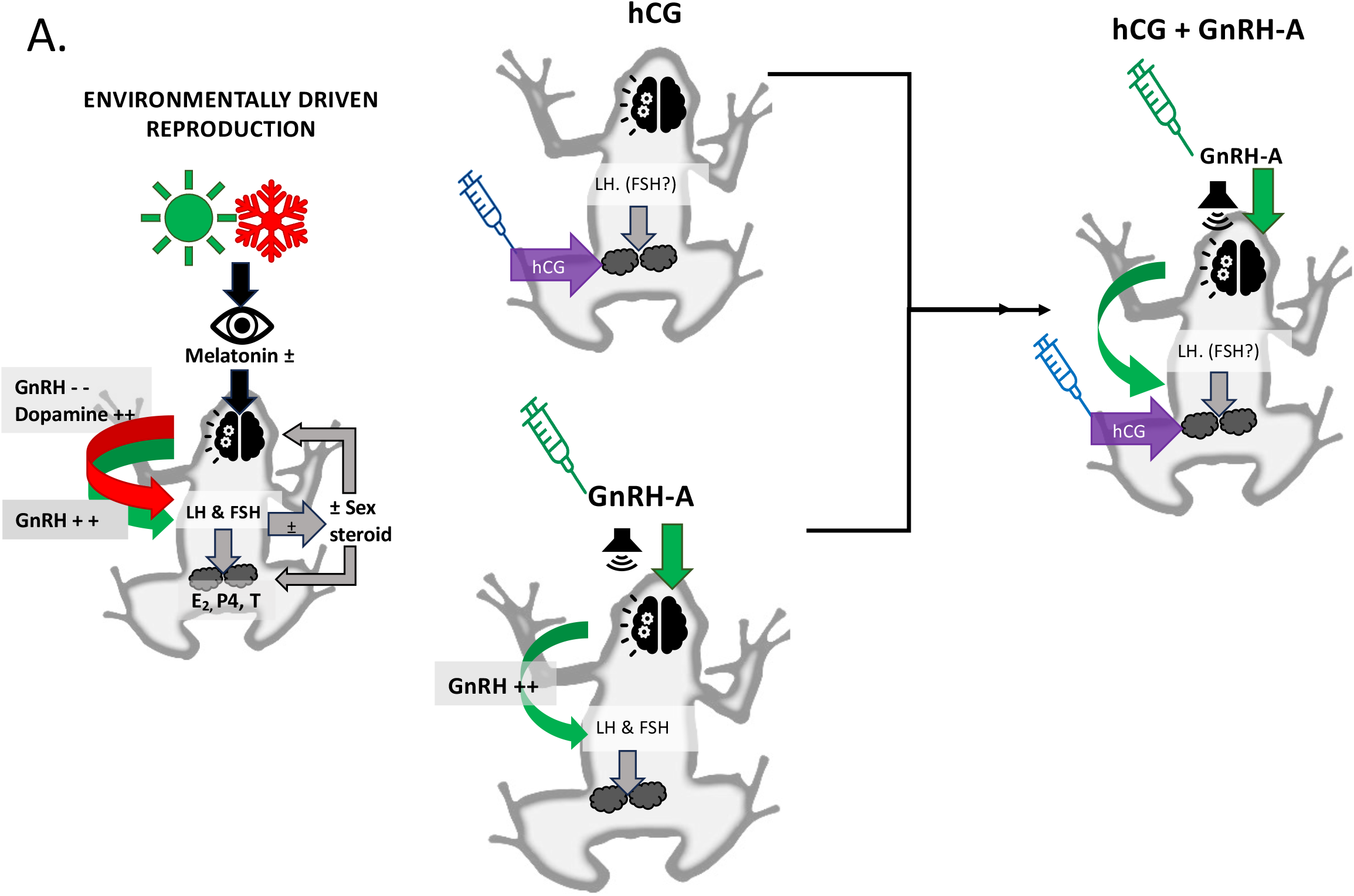
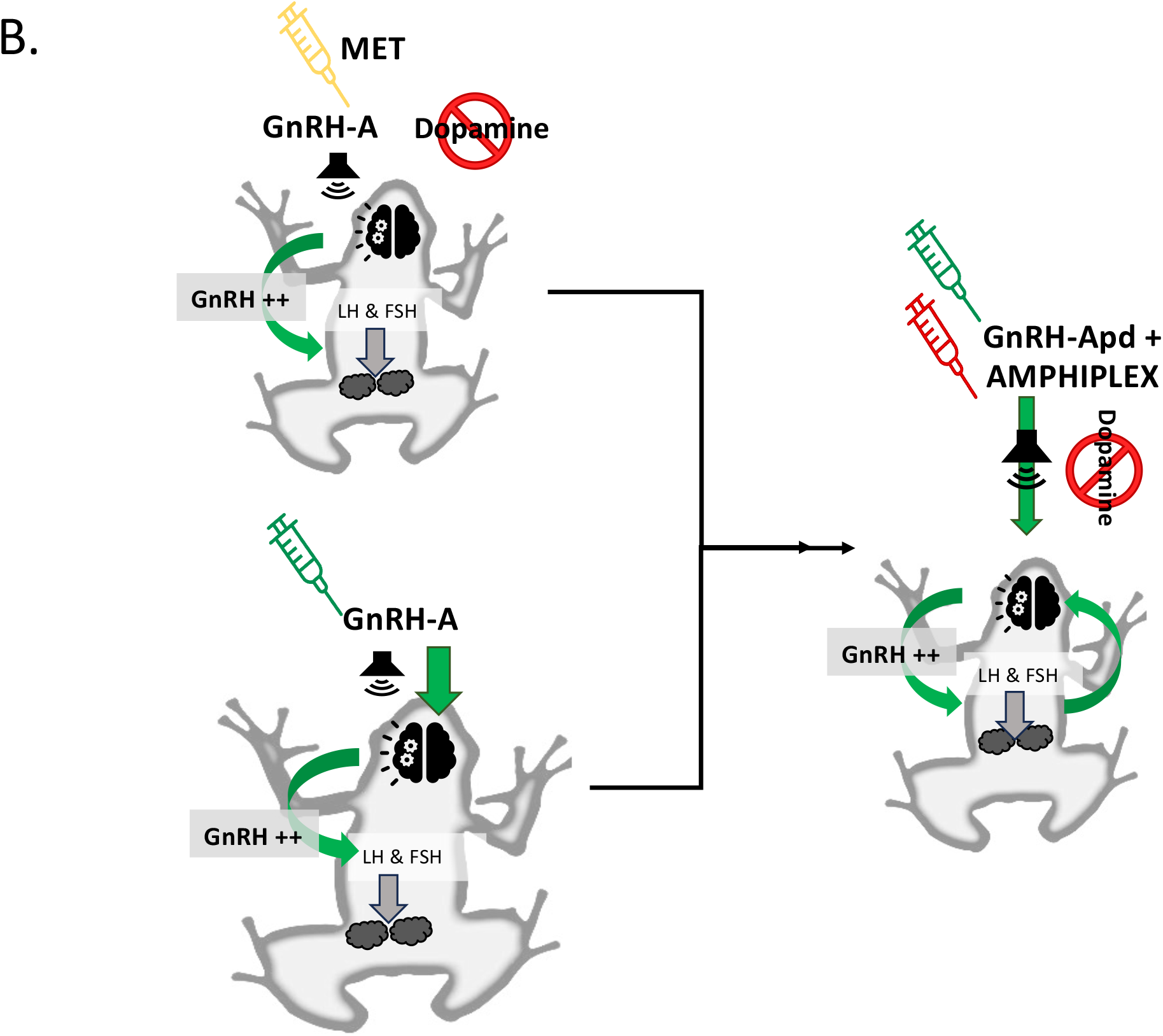
Simple schematic of environmental and hormonal control of reproduction in discontinuous breeders. During winter, temperature and photoperiod suppress GnRH-A and downstream, LH and FSH through melatonin and dopamine control. Spring increase in temperatures and photoperiod suppressing melatonin and dopamine and releasing GnRH. Increases in GnRH lead to LH, FSH or an increase in both hormones, eventually leading to an LH surge leading and sperm release. A combination of hCG + GnRH-A could provide double stimulatory actions at the brain and testicular levels to elicit hormonal changes which release spermatozoa. hCG can also stimulate the testes directly producing the second highest sperm concentrations in this study. As hCG is the common hormone, in the two best treatments it is likely that in *A. boreas boreas* the mechanisms of sperm production are mediated via LH-mediated pathways. GnRH-A was most successful if administered in combination with hCG rather than alone. **B.** Poor response for GnRH-A indicates that it should be administered at sufficient dosages to be sufficient to extinguish the influence of dopamine in the system particularly early on in the summer whilst eliciting a strong LH-surge. MET alone stimulated sperm production when administered early in summer through a dopamine-inhibiting mechanism which allows an increase in endogenous GnRH. Amphiplex could both reduce dopamine and enhance endogenous GnRH leading to a downstream LH surge.

## 3. DISCUSSION

Expanding on previous studies in this and other anuran species, we compared the efficacy of four hormone treatments, 10 IU/g hCG + 0.4 µg/g GnRH, and 10µg/g MET administered singly or in combination, to induce sperm production. Three parameters were used as indicators of good quality sperm: concentration, motility, and proportion of live sperm [23,33,44]. Overall, our findings indicate that, 1) spermiation was successfully induced using every hormone treatment tested with varying degrees of efficacy; 2) the variability with which the efficacy each hormone treatment can have on inducing sperm of good quality should be determined by multiple quantity and quality metrics, which should be added cumulatively across multiple parameters; 3) the most effective treatment may vary depending on the month of administration is indicative of seasonal variations in hormone actions. According to the above-mentioned point we therefore found that administration of a combination of hCG with GnRH-A was the most successful protocol in promoting the release of sperm that, across all parameters tested, produced the highest concentrations of sperm, in addition to good motility and viability. Below we discuss the results of this study in relation to the above-mentioned points and explore some of the physiological mechanisms affecting concentration, motility and viability.

### 4.1 Concentration

Overall, 10IU hCG/gbw, administered in combination with 0.4 _µ_g/gbw GnRH-A or alone, induced the highest concentrations of sperm in comparison to 10_µ_g/gbw MET, 0.4 _µ_g/gbw GnRH and GnRH-Apd + Amphiplex (10_µ_g/gbw MET, 0.4 _µ_g/gbw GnRH combined), which were decreasingly effective in that order. In comparison, the highest motility and viability metrics were elicited by a single injection of hCG and had significantly higher TM, but not FPM when injected in combination with GnRH-A. For all hormones assessed, sperm concentration curves illustrate responses of at least 48 h whereafter, rapid decreases were noted from 72 - 96 hpi. Last, viability showed no significant differences between treatments up to 12 hpi with a consistent decrease in all treatments by 24 hpi. The differences in efficacy between a single hCG injection, or its combination with GnRH-A, make it difficult to favor one treatment over another but both were clearly more effective than GnRH-A-based treatments.

Higher sperm concentrations recorded for hCG + GnRH-A compared to hCG alone, could be the result of a close interplay between these two hormones (Figure 9). Under normal circumstances, GnRH would be the causative inductor of LH-surges in the amphibian hypothalamic-pituitary axis (HPA) [19,45]; however, synergistic action with hCG may have increased testicular sensitivity, explaining the enhanced concentration and viability metrics observed compared to GnRH-A-based treatments. A coupling of hCG with GnRH-A could promote b the formation and multiplication of spermatogonia at earlier stages of development which, in turn, increase the number of spermatozoa being released. This theory aligns with a study in *R. arenarum,* which reported that a combination of hCG and PMSG had comparable effect on inducing spermiation as did pituitary extracts. While PMSG was responsible for promoting only spermatogenesis, when combined with hCG increases in spermatogenesis, spermiation and interstitial cell activity occurred [46,47].

Nevertheless, differences between hCG administered alone or in combination with GnRH-A, compared to GnRH-A alone, suggest that it was predominantly hCG which was physiologically, responsible for the elevated concentration of sperm, longevity of sperm release and the number of TM and FPM sperm. When added to hCG, GnRH-A may have an additive effect in increasing concentration, longevity of sperm release and the number of TM sperm but not proportion of live cells nor sperm FPM (Table 3). The mechanisms for the above-mentioned results clearly require further investigation. However, from an endocrine perspective, the increased response to hCG also lends support to the theory that the induction of spermiation in *A. b boreas* is likely to be predominantly an LH-mediated mechanism [48]. Although the role of FSH cannot be excluded at this time, LH- and FSH-mediated actions reported in other amphibians highlight the variability with which these hormones can affect testicular activity. For example, in some species like *Bufo marinus* and *Pseudoeurycea smithii*, LH is more potent than FSH. Comparatively in other species such as *Taricha granulosa* and *Rana arenarum,* FSH is more potent than LH [49,50].

From a broader perspective, exogenous hormone efficacy has been correlated to phylogeny [19,51] and the competing efficacy of hCG compared to GnRH-A is variable and specificity could be genus or species-specific. The increased efficacy of hCG over GnRH-A in *A. b. boreas* coincides with a number of other amphibian species. In some North and Central American anurans, hCG has been favorably reported as a suitable sperm-inducing hormone in a number of toad species including *A. b. boreas* [44], *Anaxyrus fowleri* [37], *Anaxyrus baxteri* [52], *Rhaebo guttatus* [53], *Anaxyrus houstonensis* and *Peltophryne lemur* [21], suggesting a common mechanism of action in toad species.

Although GnRH-A-based treatments were lacking compared to hCG-based treatments with respect to concentration, it is worth noting that the doses of GnRH-A administered in this study induced relatively good sperm TM and FPM, albeit significantly lower than those observed for hCG-based treatments. These differences could be due to the lower efficacy of GnRH-A-based treatments, or to the low dose administered. Future studies should test higher GnRH-A concentrations in *A. boreas boreas,* as increased doses could potentially induce higher quantity and quality sperm in this species. This theory is substantiated by reports in several species (*Hyla regilla, Craugastor evanesco, Atelopus zetekki, Crinia georginia, Heleioporus albopunctatus and Anaxyrus americanus*) where concentrations of 1 - 6_µ_g/gbw, five to fifteen times greater than those reported in this study, are required to increase sperm quantity and quality [34,54–56] (Otero et al., 2023 (*in press*).

Physiologically, we hypothesize that low doses of GnRH-A administered singly were insufficient (i.e., sub physiological) to activate the LH-receptors or promote endogenous LH. As a result, a significantly lower output of sperm was observed and might also correlate to the lowered response observed for GnRH-Apd + Amphiplex sperm outputs. The dose-dependent nature of GnRH-receptor activation may require higher doses of GnRH agonists to increase neuronal firing rates and the resulting LH-surge which could not be elicited by low dose used in this study. If so, the administration of 0.4 Lg/gbw GnRH-A could instead lead to an attenuating negative-feedback loop rather than the required LH-surge. [57–59].

Despite the lower output observed in GnRH-A and GnRH-Apd + Amphiplex-treated animals, long recorded peaks (5 h for peak one and a second peak at 24 hpi) and response duration (up to 72 hpi) were not dissimilar to those observed for hCG-based treatments. The GnRH-A doses used in this study were selected based on Trudeau et al., 2010 [33], while the priming dose of GnRH-A administered 24 h prior to GnRH-Apd + Amphiplex, was chosen to test potential seasonal testicular stimulation by GnRH-Apd + Amphiplex and be inclusive of breeding and post-breeding months (discussed further in the seasonality section) [60]. GnRH-Apd + Amphiplex induced higher sperm concentrations, TM and FPM than GnRH-A alone, perhaps indicating that a priming dose of GnRH-A added to the down regulation of dopamine by MET had a boosting effect. Since GnRH-Apd + Amphiplex alone was not administered as a single dose we cannot discern whether it was an accumulation of the priming dose of GnRH-A along with GnRH-Apd + Amphiplex or whether GnRH-Apd + Amphiplex alone would have resulted in a similar outcome.

However, the prominent effect of MET injected singly at the beginning of the season highlights the involvement of the dopamine pathway associated with reproductive control in this species as in other over wintering amphibians (Figure 9). This is further supported by our results which showed that after July, MET was unreliable when injected independently, and its loss of efficacy was most likely related to the downregulation of dopamine. If this is the case then MET’s dopamine-suppressing properties decreased GnRH inhibition during the early part of the reproductive season allowing GnRH to cycle naturally, rebuilding its endogenous levels leading to sperm release. Furthermore, dopamine down-regulation may only be relevant in the beginning of the season, as it is tightly correlated with specific temperature and photoperiod thresholds, after which the increasing levels of circulating GnRH and steroids feeding back and forth between the brain and the testes negate the influence of dopamine and MET [33,61,62]. Future studies should re-examine this neurotransmitter and proceed more diligently across the months to clarify the loss of effect seen after July, and whether dopamine pathways could be suppressed by MET even in the presence of decreasing temperatures [35,60,63]. Nevertheless, these preliminary results indicate that MET did elicit a greater concentration of sperm in May and June compared to GnRH-A alone or GnRH-Apd + Amphiplex [60].

### 4.2 Time-points and peaks

The initial six time-points selected for this study were based on those reported by Langhorne et al., 2021, but with the addition of three more, totaling 11 (0, 2, 3 5, 7, 9, 12, 24, 48, 72, 96). The additional time-points were added to elucidate the total duration of hormonally induced responses across multiple hormone protocols, information not yet reported in the literature, but of immense importance to EBP breeding management strategies. All hormones tested showed responses of at least 72 hpi (hCG + GnRH-A, GnRH-A, GnRH-Apd + Amphiplex and MET) and up to 96 hpi (hCG) in the top two treatments. Even though sperm concentrations differed in orders of magnitude between treatments and time-points, the longevity of response curves was relatively equal supporting the theory that spermiation in this species is dose-dependent and favoring of LH-mediated pathways. Although the molecular mechanisms involved in these prolonged responses are unknown, in a study by Vu et al., (2017) genetic mechanisms in *Lithobates pipiens* post-MET and GnRH-Apd + Amphiplex injections illustrated regulation of at least three genes, *lhb, fshb* and *gnhr1* in a time and dose-dependent manner [35]. Further investigation into the molecular mechanisms driving spermiation in *A. b. boreas* warrants comparison between hCG and GnRH-A-based treatments to clarify genetic regulation and hormonal pathways that led to such a difference in sperm production.

In *A. b. boreas*, hormonal induction elicited at least two concentration peaks lasting between 5 and 7 h, followed by at least one other detectable peak at 24 or 48 hpi. The time gaps between the subsequent days of collection, 24 - 96 hpi obviously hinder the author’s ability to determine exactly how many additional peaks occurred over the course of the experiments. Nevertheless, it seems likely that at least one more peak was detected in all treatments and, together with the extended length of the response curves, demonstrates that an extended spermiation response can be induced in this species following any of the treatments administered in this study.

### 4.3 Motility

As was the case for concentration, treatments that had 10 IU/gbw hCG as a component had the highest number of total motility (TM) and forward progression (FPM). The increased efficacy of hCG influence on sperm motility in *A. boreas boreas* could be mediated through gonadotropin regulation of Sertoli or Leydig cells LH/hCG more specifically, influencing germ cell mitosis and mitochondrial energy production for flagellar movement [64]. Once again, the specificity with which a particular hormone can influence sperm quality parameters is evident. *A. b. boreas,* similar to other North American toad species, shows a greater proportion of motile and forward progressing sperm when injected with hCG compared to GnRH-A. Among the species that have been reported to respond positively to hCG are *A. baxteri* (80%) [52], *Anaxyrus fowleri* (60 - 90%) [37], *Atelopus zeteki* (90%) [23] and *Rhaebo guttatus* (100%) [53], all of which had motility comparable to the percentages reported in our study. Therefore, the motility parameters measured for *A. b. boreas* in this study, by comparison, were of good quality (60 - 90% TM). Setting an acceptable quality baseline for amphibians may be difficult due to the vast array of reproductive adaptations however, in the case of *A. b. Boreas,* suggests that standards reflecting that of acceptable human quality assessments, as set by the World Health Organization (WHO), could be used [65]. In this case, acceptable motility parameters for this species would require that the proportion of immotile spermatozoa exceeded 40%. In instances where this is not the case then further clarification on the quality of the sperm sample should examine the proportion of live spermatozoa as an additional metric.

### 4.4 Live/Dead Analysis

In determining the quality of a sperm sample, the percentage of immotile spermatozoa should not be higher than the percentage of dead spermatozoa [66]. Viability in this study was measured as the proportion of live cells obtained from all hormonal treatments. Our results indicated that vitality was under the influence of time (hpi), not treatment type. Viable sperm was detected at consistently high levels in the first 12 hpi (80-90%) decreasing significantly but equally by 24 hpi in all treatments. Interestingly, the trend in viability scores was opposite to that observed for concentration and motility. hCG alone had the lowest proportion of live sperm, followed by the hCG + GnRH-A cocktail, GnRH-Apd + Amphiplex and GnRH-A; however, there were no significant differences between the treatments. As a measurable indicator of fertility viability metrics such as DNA fragmentation [67], the proportion of live cells and acrosome integrity [68] can have profound implications for processes such as successful syngamy and embryonic development.

Although only analyzed in one treatment type, acrosome integrity was not as high as expected. Moreover, when examined, results indicated differences in the proportion of sperm with intact acrosomes differed across a number of distinct populations and generations, suggesting that fertilization capabilities may be variable. To date, poor fertilization and survival in the *ex situ* populations has been assumed to be related to poor sperm concentrations or gamete synchronization. However, these results suggest that sperm production is concentrated and long lasting and that issues relating to fertility may lie with poor viability of the sperm produced. Although the proportion of live sperm was good across all treatments, live cells devoid of an acrosome would be unable to fertilize regardless of high concentrations and motility. These preliminary results have management impacts for future breeding efforts, and future research should compare acrosome integrity across all hormone treatments, populations, source (captive born versus wild caught) and age ranges to elucidate appropriate viability scores for this CBP. Additionally, other factors such as pH, osmolarity and sperm morphology are necessary.

### 4.5 Duration of responses across all quality assessments

A last observation that warrants further investigation and is especially applicable to breeding management, is the duration of response observed in response to exogenous hormones. In this study we found that all hormones elicited responses of 24 - 96 hpi, despite a consistent reduction in sperm concentrations, motility, and viability after the initial 24 hpi period. Like *Bombina bombina,* which exhibits a long response time to hormones and long response time across months, *A. b. boreas* not only showed longevity in their responses over days, but also across months of testing. In *B. bombina*, the extended duration of the spermiation was attributed to pituitary cells remaining high throughout the summer months, not decreasing monotonically. If true to *A. b. boreas,* this could be reflective of a partially continuous spermiation strategy and breeding could occur over an extended period, having important captive breeding management implications [49,55]. If females are available and testes remain responsive to exogenous hormones, then captive breeding would not necessarily have to proceed directly following emergence from brumation. These types of managerial decisions would require a better understanding of female reproduction and readiness to breed. The consistent 24 - 48 h responses followed by rapid decreases observed in this study are in line with those reported in Tungara frogs, where hCG and GnRH-A cause spikes in detectable Testosterone (T) metabolites with hCG causing larger elevations in T than GnRH-A. In *A. b. boreas* we observed an almost complete cessation of spermiation by 72 and 96 hpi, which could be correlated to spiking T initiating a feedback loop. Although it is unknown in this species whether steroids are inhibitory or stimulatory (they can be either in different species), as the extended response over time and months observed in *A. b. boreas,* it is probable that stimulation of spermatogonial proliferation and rate of maturation of spermatids into spermatozoa occurs in a similar fashion to that reported in *R. esculenta, R. pipiens, R. catesbiana. R. sylvatica, B. marinus and X. laevis* [49,69] and this requires further investigation. The Boreal toad is a high elevation, long-lived species (up to 19 years in captivity and 15 years in the wild, T. Smith pers comm) (Figure 9). Spending long periods of the year under extreme cold conditions this species has evolved overwintering (brumation) as a survival adaptation. Emerging from winter once the snow melts and temperatures begin to warm, the species’ begins its activities with breeding commencing immediately [70–73]. In overwintering anurans, growth and reproduction are therefore tightly linked to abiotic factors. When environmental cues are translated in the eye and brain, causative circadian, genetic, and endocrine pathways drive spermatogenesis [19,63,74–76]. What is not widely reported are the effects of seasonality on sperm quantity and quality parameters. Unsurprisingly, in this study, we found a significant correlation between sperm production and viability depending on the month that hormones were administered. While the number of sperm with TM and FPM decreased steadily throughout the summer months, concentration showed a slight recovery in September as did viability.

In June and July, the corresponding months in which this species breeds, all parameters were at their highest, correlating with the species’ reproductive period. As previously described, hCG administered singly or in combination with GnRH-A were the most successful protocols in eliciting higher sperm concentrations compared to GnRH-A or GnRH-Apd + Amphiplex. The efficacy of hCG-based treatments could relate to the hormone’s ability to maintain high levels of spermatogenesis in toads and further lends support, as previously mentioned, to the mechanisms of spermiation in the *A. b. boreas* being LH-mediated. Existing evidence of hCG-action on testicular and interstitium function has been demonstrated in *R. temporaria* and *Bufo bufo,* where spermatogenesis in hypophysectomized males was recoverable through injections of hCG which mirrored plasma levels of LH during breeding months [63,77].

Though there was a consistent decrease in sperm concentration from June to July the most noticeable dip occurred in August, the onset of autumn. Though it is not clear if *A. b. boreas* is a strictly discontinuous or a partially continuous breeder, the onset of August may be reminiscent of testicular refractoriness influenced by a number of mechanisms. Endocrinologically, increases in interstitium development would act in opposition to the spermatogenic cycle with changes in abiotic factors such as temperature, promoting androgen secretion in the Leydig cells [78–80]. As with *R. catesbiana* it is possible that in *A. b. boreas,* hCG promotes LH-mediated spermatogenesis in June and July but as temperatures drop in Autumn, hCG’s influence changes, increasing the interstitium’s sensitivity to LH and androgen production. Increases in the latter, would also explain the dip in sperm concentration and viability observed in August. As changes in steroid secretion impact Leydig cell morphology and, androgen and estrogen levels increase, estrogen receptor mediated interactions with LH-receptors in Leydig cells could create a feedback loop on androgen biosynthesis. These results correlate with those reported in other brumating species such as *Bufo japonicus*, where the mechanisms of GnRH are seasonally immunoreactive and show greater immunoreactivity during the spring and autumn months [81]. In *Xenopus laevis,* GnRH _c_oncentration is higher in reproductively active frogs in the breeding season (spring) compared to winter months [82]. Unique to amphibians is the presence of multiple forms of GnRH which may play roles other than gonadotropin regulation. Nevertheless, isolation of GnRH from the testes in *Rana esculenta* [83] showed higher levels of immunostaining during June and September in the interstitial and Sertoli cells compared to December. These seasonal variations highlight, amongst other factors, the involvement of changing photoperiod and temperatures as regulators of testicular productivity. The monthly patterns observed for concentration were also observed in the proportion of TM, FPM and live cells. A resurgence of viable cells in September compared to August are indicative of a refractory period in *A. b. boreas* while the higher levels of TM, FPM and live spermatozoa in June and July reiterate these as being the peak breeding months compared to later in the year.

Thus, it is possible that as the breeding months come to an end, differentiation of pre-meiotic germ cells and primary spermatogonia reported in other anurans increases but the availability of mature sperm becomes limited [84]. Although the complexity of steroidogenesis on promotion and inhibition of spermiation is beyond the scope of this discussion, a significant increase in sperm concentration was observed in September compared to August reminiscent of testicular recrudescence [78]. Injections of hCG in September may interact with steroid-mediated spermatogenesis by increasing differentiation of seminiferous tubules leading to renewed spermatogenic potential. In *Rana pipiens* and *Rana temporaria,* the appearance of early sperm stages undergoing the first meiotic division clearly indicates that in the testes of seasonal anurans spermatogenesis can continue cycling even if sperm do not have to be released [79,80]. This seems likely in *A. b. boreas,* however, this requires further investigation.

GnRH-A and GnRH-Apd + Amphiplex showed a markedly lower elicitation of sperm production compared to hCG alone across all months tested, but when combined with a priming dose of GnRH-A, GnRH-Apd + Amphiplex performed better than GnRH-A, indicative of the potentiative effects of a priming dose and/or the benefits of including MET into the injection. The reasons for this must be multifactorial but could relate to dosages or the effects of MET on dopamine inhibition.

Unfortunately, not enough data on MET administration alone was collected as this might have better explained the independent role of this neurotransmitter especially in the context of breeding versus non-breeding season. Trudeau et al. (2013) showed that a priming dose of GnRH-A administered 24 h prior to GnRH-Apd + Amphiplex helped potentiate the effects on sperm production when administered out of season [60]. Since we set out to have a uniform treatment we included the GnRH-A priming dose in all months of treatment whether breeding or non-breeding. In *A. b. boreas* a priming dose of GnRH-A given 24 h before GnRH-Apd + Amphiplex did not drastically change sperm output in the reproductive, compared to non-reproductive months. Interestingly, though it was not a particularly effective inducer of spermiation in this species at any point during the study from a concentration perspective it did produce reasonable motility and a high proportion of live cells.

### 4.6 Hormone protocol and EBP reproductive management

Understanding the longevity and magnitude of spermiation output of these treatments is of the utmost importance for EBP managers when making decisions to pair and inject animals for breeding or artificial fertilization. This study clarifies the timing and duration of spermiation peaks for at least five treatments and combined with the ovulation protocols outlined in Calatayud et al., 2015 and 2017, indicates that males paired with females should have at least a 24 h window in which to fertilize egg [29,30]. More specifically, this study adds to the existing literature which would advise pairing males with females close to the time of oviposition and within the first 12 hpi of administering hormones to males [30]. Based on the seasonal nature of this species further recommendations would be to acclimatize animals for a period following emergence from brumation before pairing for breeding. Given the artificial nature of the brumating protocols for this EBP, we recommend more research be conducted on the naturalization of over wintering protocols.

This study also provides new information on sperm quality metrics data that are rarely recorded by EBP’s. Wide screening of quality metrics is needed to ascertain the viability of the male population. Although the proportion of live sperm in all treatments tested was relatively high +80%, the proportion of live cells, compared to cells with intact acrosome integrity, in one of the hormone groups (hCG + GnRH-A), indicated that at least 40% of sperm were devoid of this structure. If similar across other hormone treatments, this result could point to a significant loss in viability in the captive population, since the acrosomal reaction (AR) is necessary for sperm fertilization. In humans, males are considered infertile when sperm viability falls below 75 −80% sperm regardless of whether sperm are motile [85]. Therefore, assessments that do not include these metrics could be underestimating male infertility [86]. For external fertilizers such as *A. boreas boreas,* viability could be an important measure of the longevity of inheritable genetic material when implementing amphibian ARTs especially if the induction of sperm is to be used to genetically manage or for long-term storage of the species’ genetic diversity. Additionally, other considerations for future research are recommended for example:

- Age at time of breeding or gamete collection
- Genetic population
- The source (captive versus wild) where animals were hatched.
- Whether generational domestication has played a role in loss of reproductive viability.

Further to these findings, our results serve as important information that can be used to plan informed breeding management strategies for this species. To this we recommend the following:

a. Informed hormone strategies with which *ex situ* breeding management protocols can be applied are now available for this species and should be strongly based on natural history-based research to enhance the animals’ natural reproductive cycles instead of relying on the application of ARTs to overwrite reproduction.
b. Greater understanding that stimulation of sperm production does not necessarily reflect an increased quality of gamete release and protocol optimizations should be explored in conjunction with environmental quality, genetic diversity, husbandry and generational captivity.
c. That well researched protocols should be properly tested and replicated as a discerning strategy for enhancing reproductive strategies.

## 4. Conclusion

Our data indicated that when developing spermiation protocols, best practices should strive to incorporate more sperm quality parameters than just concentration and motility. With additive assessments such as viability and morphology, it is clear, that higher sperm concentration outputs are not necessarily reflective of sperm with fertilizing capabilities. Particularly, in instances where captive management requires the development of more standardized exogenous hormone protocols, several quality metrics should be used to find a best fit approach. Furthermore, the goals of exogenous hormone use should be carefully considered. A hormone protocol that is developed for the induction of breeding could look very differently to a protocol that attempts to retrieve maximum quality sperm for genetically aimed management such as artificial fertilization or sperm cryopreservation. Finally, the use of hormone protocols may require careful coordination with other factors to work optimally, such as temperature, water quality, time of day hormones are administered, month, photoperiod and latency following a period of brumation. More emphasis on the goals for which these protocols are being developed is necessary as is the consistency with which their application and data is collected by ECPs. Moreover, a better understanding of the natural history, evolution and reproductive modes must be considered, when using hormones to induce gamete release to enhance the efficacy, not bypass poorly understood physiological mechanisms.

The results of this study suggest that there is still more scope to optimize hormone protocols for this species including:

1. The exploration of higher doses of GnRH-A (1 - 6μg/g BW) as alternatives to the low dose tested herein (0.4 μg/g BW).
2. The month of administration affects spermiation in response to hormonal induction regardless of treatment indicative of the seasonality of the species.
3. That optimal hormone concentrations must elicit higher sperm quantities without compromising motility or the proportion of live sperm.
4. Preliminary results suggest a population difference in acrosome integrity that could affect the fertilizing capabilities of some of the populations held at NASRF, accounting for the poor fecundity observed in the *ex situ* population.

## 5. ABBREVIATIONS

(SRM): Southern Rocky Mountain boreal toad population
(hCG): Human chorionic gonadotropin
(GnRH-A): Gonadotropin Releasing Hormone agonist also referred to as GnRH analog [des–Gly10], D–Ala6 ethylamide acetate]
(MET): Metoclopramide
(µg / g BW): micrograms per gram of body weight
(IU): international unit
(mL): milliliter
(µL): microliter
(hpi): hours post injection
(n=): number of
(°C): degrees centigrade
(cm): centimeters

## Supporting information

Supplementary Protocols

Supplementary Odds Ratio Tables

## 6. ACKNOWLEDGEMENTS

Natalie Calatayud would like to thank first and foremost, Exploradora de Inmuebles, S.A. (EISA) for awarding financial aid for her first two years as a postdoctoral fellow. We thank our collaborators at the Colorado Division of Parks and Wildlife, H. Crockett and M. Nicholl, as well as T. Smith, (Hatchery Manager) and the staff at the Native Aquatic Species Restoration Facility in Alamosa, Colorado, for allowing me to live and perform these studies at NASRF. In particular, NEC would like to thank Mr. T. Mix, Mr. J. Houghtaling, Mr. D. Westerman and Mr. N. Heredia for assisting me with my experiments during my 2014 and 2015 seasons. A final thanks to Debra Shier and Dr. Barbara Durrant for their support during my postdoctoral fellowship. Finally, and most importantly, we would like to respectfully acknowledge the Ute Indians, Capote and Mouaches, traditional custodians of the land on which this work was conducted and pay our respect to Elders both past, present and emerging.

## 8. AUTHOR’S CONTRIBUTIONS

NEC obtained the funding, experimental design and conceptualization, data collection and analysis and data interpretation, manuscript writing and editing. LJ data analysis and interpretation, manuscript writing and editing. GDT was involved in data analysis and interpretation, manuscript writing and editing. CJL was involved in experimental design, data collection, data analysis and interpretation, manuscript writing and editing. AM was involved in data collection. RU was involved in experimental design, data analysis and interpretation, manuscript writing and editing.

## 9. ETHICS APPROVAL AND CONSENT TO PARTICIPATE

The Native Aquatic Species Restoration Facility (NASRF) Alamosa, Colorado Animal management and research studies reported here were reviewed and approved by the San Diego Zoo Institutional Animal Care and Use Committee (IACUC # 14-005).

1 Clear Creek males were the only 2 founders remaining from the original group collected in 2000.

