## Supplementary Protocols for "Hormonal induction, quality assessments and the influence of seasonality on male reproductive viability in a long-term managed *ex situ* breeding colony of Southern Rocky Mountain Boreal toads, *Anaxyrus boreas boreas*"

### ASSISTED REPRODUCTIVE TECHNIQUES (ART) FOR BOREAL TOADS

#### INTRODUCTION

*Gametogenesis and the endocrine function in amphibians*

Control of endocrine function and gametogenesis in amphibians is regulated through the hypothalamic-pituitary axis (HPA) and is carefully synchronized through a series of processes incorporating endogenous and exogenous (environmental) signaling. Exogenous cues such as temperature, light and humidity can be powerful regulators of amphibian reproduction, particularly in species that inhabit higher latitudes and altitudes (Morrison, 2003). However, in captivity the optimum environmental conditions and nutrition requirements cannot always be properly provided and in this instance, health and reproduction can become compromised.

The administration of exogenous hormones promotes physiological signaling mechanisms that may be inhibited in captivity by a lack of appropriate environmental cues (Kouba and Vance, 2009). However, hormones should only be administered if and when husbandry procedures have not been sufficient to elicit natural breeding or the exact requirements to provide the correct environment for breeding are unknown. In addition, hormones should be administered during the breeding season only, as evidence suggests that induction of reproductive behaviors and gamete production out of season can be unreliable (Szabo et al., 2000; Trudeau et al., 2010; 2013). Detailed records of the doses and number of injections given to each individual should be carefully logged to avoid over stimulation. To the researcher's knowledge, there is no evidence in the literature that suggests the use of exogenous hormones can have long lasting deleterious reproductive or epigenetic effects on amphibian health. However, ovarian overstimulation by exogenous hormone injection has been well documented in mammals and can have serious deleterious effects on the reproductive health of an individual that can be long lasting. Therefore, anyone considering artificial stimulation should take a conservative approach to the administration of hormones, particularly when using those that are not normally found in amphibians, ie Human chorionic gonadotropin (hCG).

The hypothalamic-pituitary complex in amphibians is composed of several identifiable hormones and neuropeptides that are similar to or have analogues of those found in fish, reptiles and mammals. The group of pituitary hormones that are directly involved in gametogenesis, steroidogenesis and spawning are termed gonadotropins. Oocyte maturation, egg yolk deposition, testicular androgen production and spermatogenesis are controlled by gonadotropins (Licht, 1979; Polzonetti-Magni et al., 1998). As in other vertebrates, the LH surge is critical to ovulation and sperm release (Herbison et al., 2008; Tischkau et al., 2011).

#### HORMONES FREQUENTLY USED FOR ARTIFICIAL STIMULATION OF OVIPOSITION, SPERMIATION AND BREEDING BEHAVIORS IN BOREAL TOADS

*Gonadotropin releasing hormone (GnRH)*

93 Perry Street Error! **Bookmark not defined.**, Error! **Bookmark not defined.** Phone: Error! Bookmark **not defined.** Fax: Error! Bookmark **not defined.**
Error! Bookmark **not defined.**
E-Mail: Error! Bookmark **not defined.** Web: Error! Bookmark **not defined.**

The most common forms of GnRH found in amphibians are similar to the mammalian (mGnRH) and chicken GnRH (cGnRH) (King and Millar, 1986; Sherwood et al., 1986; Licht et al., 1994). In amphibians, the mGnRH is localized to the pre-optic area and cGnRH is found more widely distributed in the brain and spinal cord. It has been suggested that mGnRH is involved in the synthesis of goandotropins, Lutenizing hormone (LH) and follicle stimulating hormone (FSH) while cGnRH acts as a modulator of sexual behaviors (Collin et al., 1995; Licht et al., 1994).

In the boreal toad, GnRH (Des-Gly$^{10}$,D-Ala$^{6}$,Pro-NHEt$^{9}$)-GNRH acetate salt, BACHEM, USA) alone can induce spermiation and amplectic behaviors (Calatayud et al., 2015; Calatayud et al., unpublished) and has been commonly used in the past to induce oviposition (Calatayud et al., 20915). Recorded doses of GnRH used to induce oviposition, spermiation and reproductive behaviors are shown in table 1.

**Table 1.** GnRH hormone protocols for inducing reproductive behaviors and gamete deposition in the boreal toad (see table 4 for product information)

| **Physiological response** | **Application** | **Number of doses (per body weight)** | **Time between injections** | **Response time** |
|---|---|---|---|---|
| Spermiation& amplexus | Priming dose 1 | ^0.2-0.4 µg/g | 24 hr | 3-6 hr** |
| Oviposition | Secondary dose* | 0.4-0.6 µg/g | | |
| | Priming dose 1 | ^0.2-0.4 µg/g | 24 hr | 24 – 96 hr after secondary dose** |
| | Secondary dose* | 0.4-0.6 µg/g | | |

*^If unsure about doses to start with, always begin with lowest dose*
**Secondary doses are usually administered in combination with metoclopramide a dopamine inhibitor.*
***Secondary doses may not be necessary if male becomes amplexed within 12 hr of first dose and female oviposits within 24 hr of priming dose.*

Physiological responses to GnRH may take longer than other hormone such as hCG because they act on the brain, not directly on the gonads. However, GnRH is considered the safest because it is conserved across all taxonomic groups in structure and function and is a naturally occurring hormone in amphibians. Unless other environmental issues are of concern, this protocol should enhance existing levels of systemic GnRH after hibernation in boreal toads. Amplectic behaviors, spermiation and oviposition will occur if the animals are at an optimum breeding capacity as GnRH will promote the LH surge required for appropriate breeding. This hormone can also be combined with other hormones such as, hCG and neurotrasmitters such as, metoclopramide and domperidone.

More detailed information on the action of GnRH can be found in Chapter 2: Assisted Reproductive technologies (ART) for Amphibians and in the Association of Zoo and Aquariums (AZA): Amphibian Husbandry Resource Guide. Vicky A. Poole, National Aquarium – Baltimore Shelly Grow, Association of Zoos & Aquariums Edition 2.0, 4 April 2012.

*Dopamine inhibition using metoclopramide (MET)*

Neuropeptides are involved in reproduction through the stimulation and inhibition of GnRH secretion. Dopamine secretion in fish (Popesku et al., 2011) and amphibians inhibits GnRH production and in turn ovarian and testicular function (Schueler et al., 2013; Sotowska-Brochocka, 1994; Trudeau et al., 2010; 2013). When administered with exogenous GnRH, dopamine antagonists such as metoclopramide and domperidone, release dopamine inhibition in the brain thereby increasing GnRH levels, which culminate in an LH surge. This LH surge then affects downstream secretion of steroid hormones in the gonad stimulating folliculogenesis and spermatogenesis. In 2010, Trudeau et al., reported the use of Amphiplex, a protocol that combined Gonadotropin releasing hormone (GnRHa) (specifically des-Gly$^{10}$, D-Ala$^{6}$, Pro-NHEt$^{9}$-GnRH) and a dopamine antagonist (metoclopramide) (Table 2) (Trudeau et al., 2010). Trudeau and colleagues (2010) demonstrated that Amphiplex, as part of a two injection priming protocol, could elicit ovulation, spermiation and reproductive behaviors in 4 species of anurans (*Lithobates pipiens, Ceratophrys ornate, Ceratophrys cranwelli and Odontophryns americanus*). Amphiplex has been used in amphibians in multiple zoos including, Omaha, Nashville, Vancouver Aquarium and National Zoo (Vance Trudeau pers. comm.). Since the initial study, Amphiplex has also been tested successfully in the Northern Cricket frog, *Acris crepitan,* (Snyder et al., 2013) the Hellbender, *Cryptobranchis alleganiensis* (Vance Trudeau, pers. comm.) and the Panamanian Golden frog, *Atelopus zeteki* (Gina Della Tonga, pers. comm). Amphiplex has also been tested in the boreal toad and has proved a safe hormone protocol, although it has not been thoroughly tested Amphiplex can elicit amplectic behaviors and spermiation in the boreal toad (Calatayud et al., unpublished); however, its effects on oviposition need further documentation.

**Table 2.** GnRH and Metaclopromide (Amphiplex) hormone protocols for inducing reproductive behaviors and gamete deposition in the boreal toad (see table 4 for product information)

| **Physiological response** | **Application** | **Number of doses (per body weight)** | | **Time between injections** | **Behavioral and physiological response time** |
|---|---|---|---|---|---|
| | | **GnRH** | **Metoclopramide** | | |
| Spermiation& Amplexus | Priming dose 1 | ^0.2-0.4 µg/g | none | 24 hr | 3-6 hr* |
| | Secondary dose | 0.4-0.6 µg/g | 10 µg/g | | |
| Oviposition | Priming dose 1 | ^0.2-0.4 µg/g | none | 24 hr | undetermined |
| | Secondary dose | 0.4-0.6 µg/g | 10 µg/g | | |

**Spermiation and amplexus may not occur simultaneously*

*Human Chorionic Gonadotropin (hCG)*

Human chorionic gonadotropin (hCG) is a mammalian hormone produced by the embryo following implantation. Its pituitary analog is LH and due to its similarity to LH, it can be used to induce ovulation as well as testosterone and sperm production. In *Xenopus*, hCG can induce oocyte maturation via increased progesterone production in the ovary, and at higher doses can also induce ovulation (Fortune et al., 1975). Using priming doses of hCG and an ovulatory dose of hCG and GnRH can induce oviposition in unamplexed female boreal toads (86%, Calatayud et al., 2015) after hibernation. The responses observed in boreal toad studies after injection of this hormone priming protocol may indicate a similar synergistic increase in progesterone in females to that described previously. The administration of hCG and GnRH, in combination as a final ovulatory dose, may provide the LH surge required to elevate progesterone to an ovulation-inducing threshold. In males, spermiation and amplectic behaviors occur 3-9 hr (spermiation and amplexus) post hCG injection (Calatayud et al., manuscript in preparation). In some cases spermiation post-injection can last between 24-96 hr (Calatayud et al., unpublished*).

**Table 3.** A combination protocol of hCG and GnRH for inducing reproductive behaviors and gamete deposition in the boreal toads using (see table 4 for product information)

| Physiological response | Application | Number of doses (per body weight) | | Time between injections | Behavioral and physiological response time |
|---|---|---|---|---|---|
| | | **hCG** | **GnRH** | | |
| Spermiation& amplexus | Single dose | 10 IU/g | 0.4 µg/g^ | 24 hr | 2-3 hr*<br>3-6 hr** |
| Oviposition | Priming dose 1 | 3 IU/g | None | Day 1 T=0 hr | |
| | Priming dose 2 | 3 IU/g | None | Day 3 T=72 hr | 18-48 hr*** |
| | Ovulatory dose | 13.5 IU/g | 0.4 µg/g | Day 4 T=96 hr | |

*^GnRH can be administered with hCG optionally. This combined protocol has resulted in longer amplexus times and enhanced spermiation responses (Calatayud et al., unpublished*).*
** Spermiation times*
*** Amplexus*
****The beginning of oviposition is usually detectable by pronounced abdominal contractions*

**Table 4.** Hormone information, suppliers, catalogue numbers and distributing countries

| Hormone | Synonym | Company | Catalogue number | Presentation | Country of origin |
|---|---|---|---|---|---|
| **GnRH** | Des-Gly$^{10}$,D-Ala$^{6}$,Pro-NHEt$^{9}$)-GNRH acetate salt (Alarelin, Dalarelin, Surfagon) | BACHEM | H-4070.005 or H-4070.0025 | 5 mg or 25 mg | USA |
| | GnRH analog ([des-Gly10], D-Ala6 ethylamide acetate salt hydrate), | Sigma-Aldrich | L4513 | 1 mg, 5 mg, 25 mg | USA |
| **hCG** | Human Chorionic Gonadotropin | Sigma-Aldrich | C1063 | 1 vial or 10 vial packs at 2500 IU / vial | USA |
| | | DRS labs | Look up hCG | 5 vials of 5,00 IU/ vial | United Kingdom |
| **Metoclopramide** | 4-Amino-5-chloro-*N*-[2-(diethylamino)ethyl]-2-methoxybenzamide, methoxychloroprocainamide | Sigma-Aldrich | M0763 | Powder: 10, 25 and 100 gr | USA |

93 Perry Street Error! **Bookmark not defined.**, Error! **Bookmark not defined.** Phone: Error! Bookmark **not defined.** Fax: Error! Bookmark **not defined.**
Error! Bookmark **not defined.**
E-Mail: Error! Bookmark **not defined.** Web: Error! Bookmark **not defined.**

**Figure 1**. Amplexed boreal toads breed 18-96 hr post hormone stimulation. Duration of time to oviposition will depend on the female's readiness and the hormone protocol used.

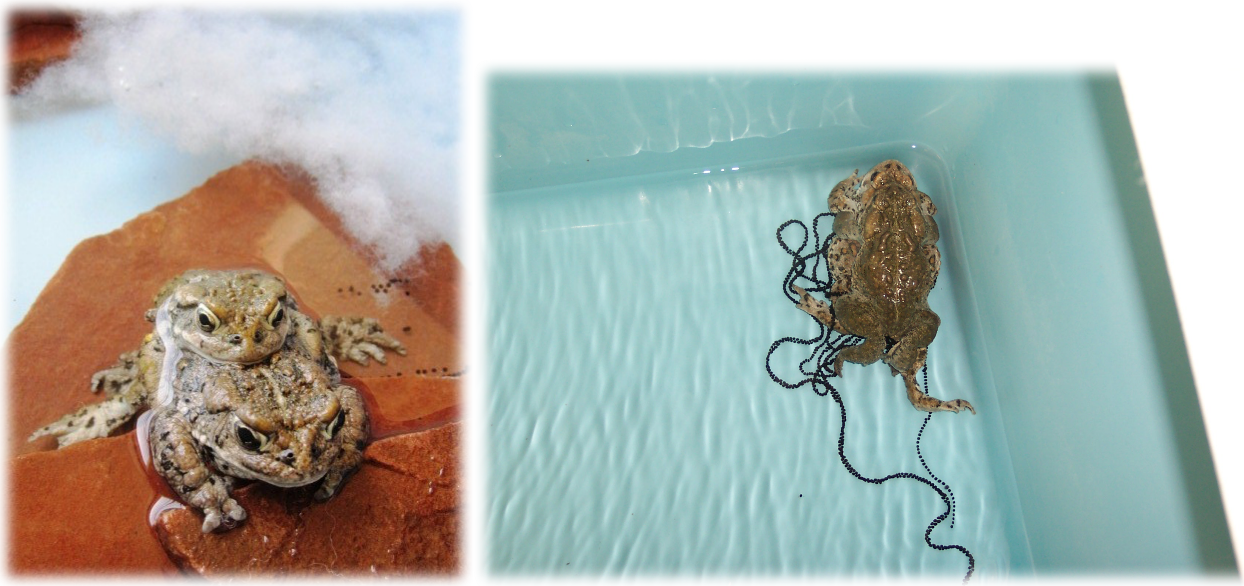

#### GENERAL GUIDELINES TO BREEDING

Using ultrasound and previous breeding history, which females are ready to breed can be determined. Female boreal toads in captivity have been bred every other year in the past, but based on recent data published by Muths et al., (2010), females should not be expected to breed more than every other year and in some cases every 3rd to 4th year. If proper identification of clutches and parents is required it is recommended that, once amplexed, any surplus animals be removed to allow the amplexed pair adequate space. Tanks should contain a platform to allow frogs to come out of the water as needed. Appropriate accessories, such as corrugated plastic tubes or stone slabs, can be placed in each tank to provide a surface for the deposition of eggs. Each pair should be maintained together for at least a week or until all breeding activity ceases. Breeding tanks should be monitored daily for the deposition of eggs. Security cameras may be used to record behaviors 24 hours per day during the breeding season if determination of egg deposition timing is important.
Frogs should be paired for breeding immediately after hibernation with a water temperature of approximately 40-50°F. Multiple males and female frogs should be introduced to each other and should be allowed to pick a mate. The ratio of males to females, if possible should be 3:1 to allow for selection and competition. Eggs should be removed from breeding enclosures soon after being laid. Frogs can be fed once they have finished breeding (marked by the termination of amplexus).

Our data suggests that during the breeding season, male boreals will respond to any of the previously listed hormone protocols. GnRH with hCG, hCG and Amphiplex all result in amplexus however, in 2012-13 experiments suggested that injecting the males is not required (Calatayud et al., manuscript in preparation). Instead, amplexus and spermiation appear to be correlated to the female's readiness to breed, therefore, stimulating only the females to oviposit may be the best way to promote breeding. Our studies also indicate that if females are to be treated, exposing the males to them from the time they are first prime elicits mating behaviors such as amplexus in the males (Calatayud et al., manuscript in preparation).

#### POST OVIPOSITION: EMBRYONIC ASSESMENTS

*Fertilization and embryo cleavage assessments*

In boreal toads the first signs of cleavage appear approximately 2-3 hr post-fertilization. Although cleavage rates are dependent on water temperature, at 40-50°F cleavage rates maybe first visible between 3-4 hr. Once a clutch has been deposited completely, carefully remove the eggs and transport them to their final rearing tank. To keep careful records of development, assess development every 24 hr where possible and remove dead or unfertilized eggs and embryos to avoid the spread of water born molds. If mold becomes a problem daily dips in amphibian ringers solution should help slow the spread. Salt solution dips are helpful in treating mold infections when embryos are exposed to warmer aquatic temperatures (65-85°F) during periods of artificially induced growth. However, if development does not need to be increased, cold temperatures (40-49°F) are the best preventative measure to help prevent mold growth.

Embryonic development in boreal toads is typical of amphibians (Figure 2; left to right). From the first division, (4 cell) at 3 hr post-fertilization there is approximately a 20-24 hr period before the embryo reaches gastrula (Gosner stage 11) (Gosner, 1960). Hatching occurs in 6-10 days. Development thereafter is temperature, food and density dependent.

**Figure 2**. Embryo at a) 4-cell (Gosner 4), b) 32-cell (Gosner 7), c) gastrula (Gosner 11), d) newly hatched tadpoles (Gosner ~23), e) late stage tadpoles (Gosner 26-30) and f) metamorph in tail reabsorption stage (Gosner 43-45) (Gosner, 1960).

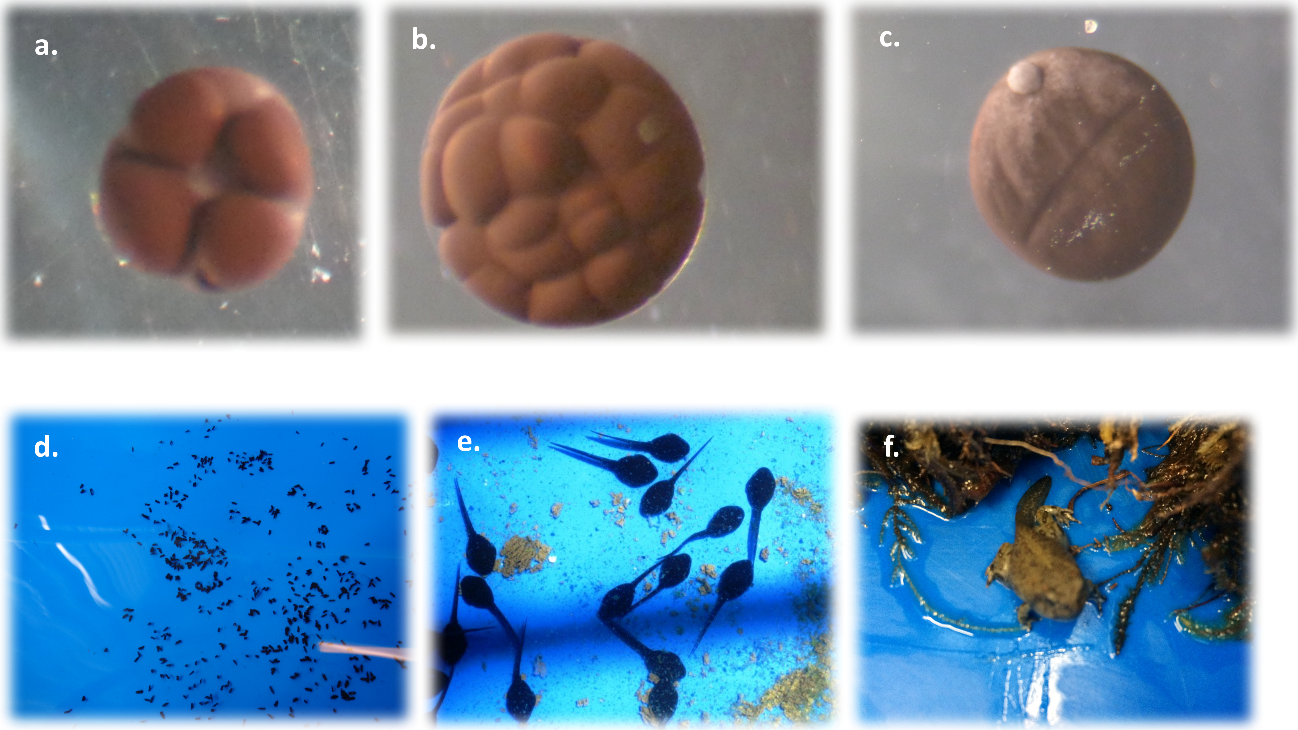


Tadpoles and sub-adult frogs produced from captive breeding may be reintroduced to the wild in alignment with conservation goals. A sufficient number of animals should remain within the captive colony to ensure future-breeding goals can be met.

#### COLLECTION AND ASSESSMENT OF BOREAL TOAD SPERM

*Sperm collection*

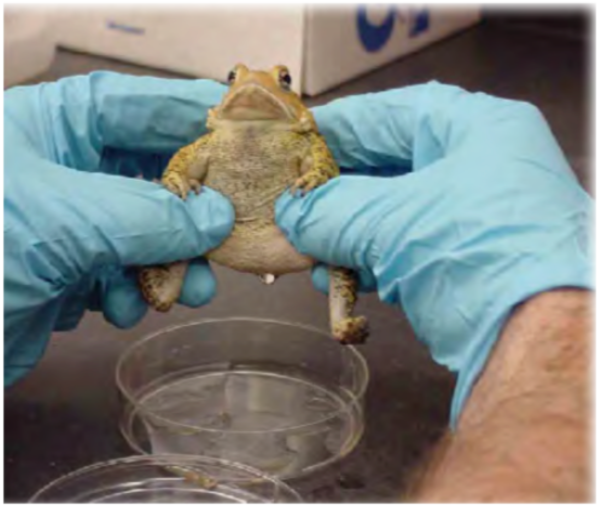

**Figure 3.** Spermic urine collection: The *cradle method* collection of spermic urine from a male Fowler toad. First, quickly dry the animal's ventral abdomen with a tissue to ensure that all the liquid collected is urine. Hold the male over a small Petri dish using the ring finger to spread his legs apart and induce urination. It is important that some movement is allowed, because this can help the animal to express urine in response to normal muscle contraction in the legs. Photo taken from Kouba et al., 2012.

To collect urine, males must be well hydrates in order to produce spermic urine. To ensure hydration, place males into small plastic container with 2-4 centimeters of water for at least 15 minutes prior to attempting to collect spermic urine. If the toad has been sitting in water prior to being caught, a period of hydration may not be necessary. Hold the toad using the cradling method outlined in AZA Amphibian Husbandry Resource Guide (Kouba et al., 2012), over a dry plastic container with the abdomen pointing down towards the dish (Figure 3). Softly massage the abdomen to encourage urination for no longer than 2 minutes, or less if urination occurs. Should the toad fail to urinate with this technique, spermic urine can be obtained by inserting a small disposable vinyl catheter (0.34mm x 0.052mm) (Scientific

**Figure 4.** Urine collection from a Mountain Yellow-legged frog using a catheter

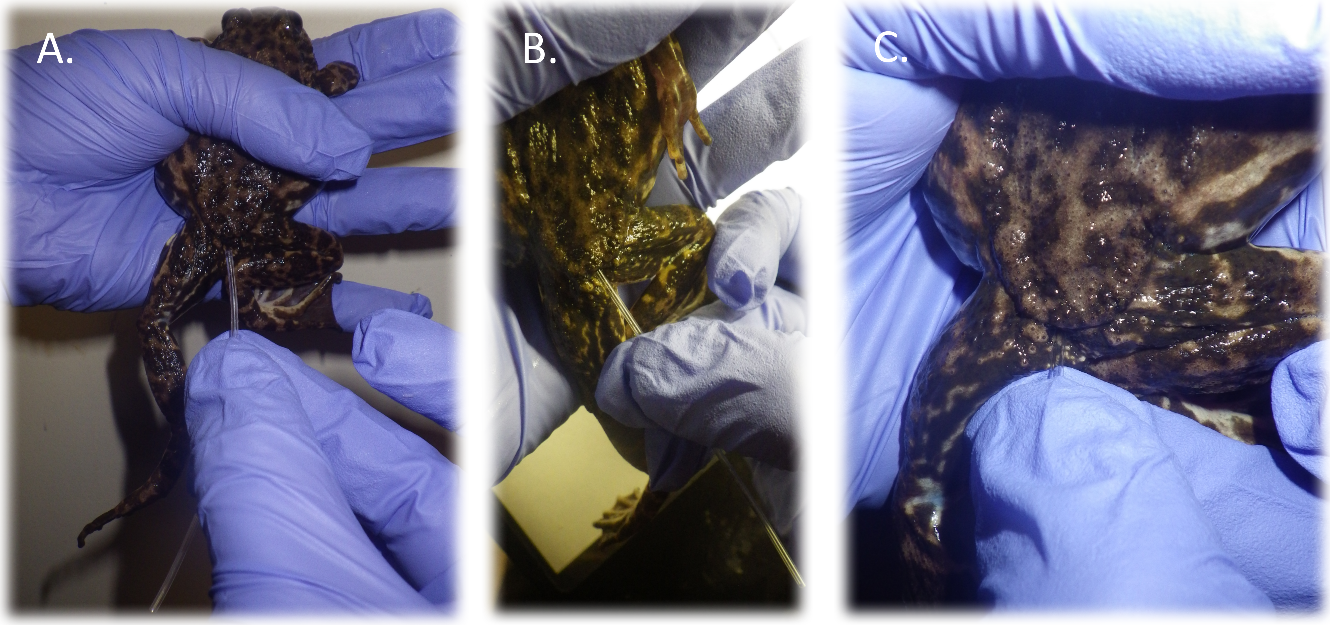


Commodities cat #: BB31785-V/5) into the cloaca to stimulate urination (as described in Kouba et al., 2012a). Catheter collections should not last longer than 30 seconds and the catheter should be inserted no more than 4 mm into the cloaca. This technique is successful in the boreal toad (Calatayud et al., unpublished; Calatayud et al., 2015; Kouba et al., 2012a). Place the catheter in a 2 mL tube before inserting the other tip into the toad. This will avoid loosing any sample if the catheter stimulates urination immediately. Urine samples can remain in this 2 mL centrifuge tube and stored at 4°C. boreal toads are avid urinators so collections do not usually require catheterization (Figure 4).

*Extracting eggs from a gravid female for IVF*
Before attempting to express eggs from a female check for egg jelly in the urine and signs of abdominal contractions. In boreal toads, females have begun expressing eggs within an hour or two of the first signs of abdominal contractions (Calatayud personal observation). If contractions are observed, you may not need to squeeze the female; instead wait for her to begin expressing eggs naturally. If a female is suspected of being close to oviposition but has not done so, eggs can be expressed by gently but firmly massaging the female's abdomen (Figure 3). To avoid struggling during gripping, a wet paper towel can be used to gently cover the female's head. In boreal toads, gravid females will easily express eggs if manipulated properly. Holding the toad in an encapsulating grip with the back end facing the researcher as shown in figure 2 for urine collection, and gently massaging the abdomen is usually enough to begin pushing eggs out of the cloaca.

*Sperm assessments*
Sperm concentration, motility and quality should be determined after collection to assess the sperms' suitability for in vitro fertilization (IVF) and as a tool for assessing the reproductive health of a male.
Concentrations: Counting a sample of sperm can be done using a haemocytometer (purchase from Fisher Scientific) (Figure 5) or a Makler cell counter (Fisher Scientific) (Figure 6). Ten microliters of sample containing sperm can be loaded onto either of these devices for counting. The haemocytometer and the Makler have special formulas that must be calculated after counting to obtain an accurate sperm concentration (Figure 3and 4). In boreal toads concentrations vary and we have counted ranges from 1 million up to >25 million during the breeding months. As summer progresses sperm counts may naturally decrease as will the males' production of sperm in response to hormone stimulation. If the spermic urine sample is very concentrated, a dilution of 1:10 (1 µL of sperm to 9 µL of water) can be mixed and loaded onto the Makler or haemocytometer. However, if counting a dilution, include the 1:10 into the calculation by multiplying the final concentration by 10.

**Figure 5.** Haemocytometer can be used to count sperm concentrations. It is not advisable to use it for motility since the cover glass is heavy and it can slow sperm down. However, since motility is scored subjectively, analyzing motility can be done on this device as long as the protocol remains consistent.

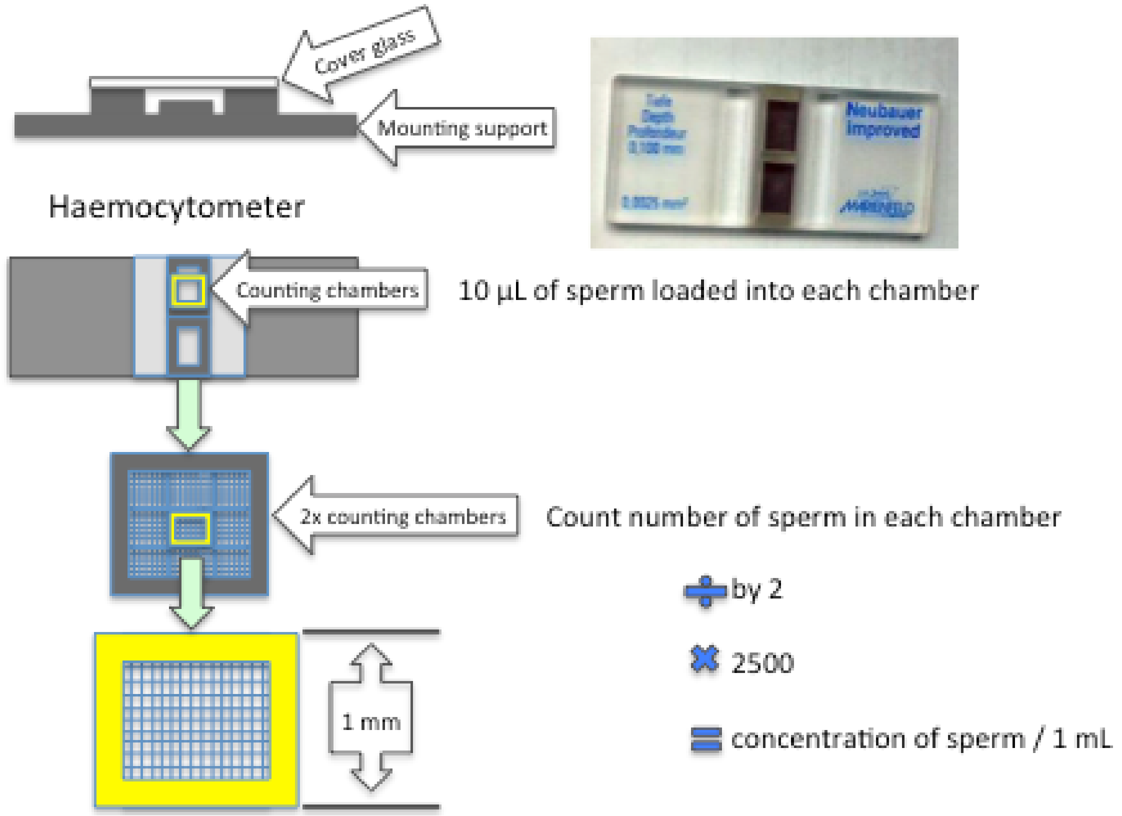


**Figure 6.** The Makler counting chamber is another instrument that can be used to count sperm cells. Its design allows motility and concentration assesments.

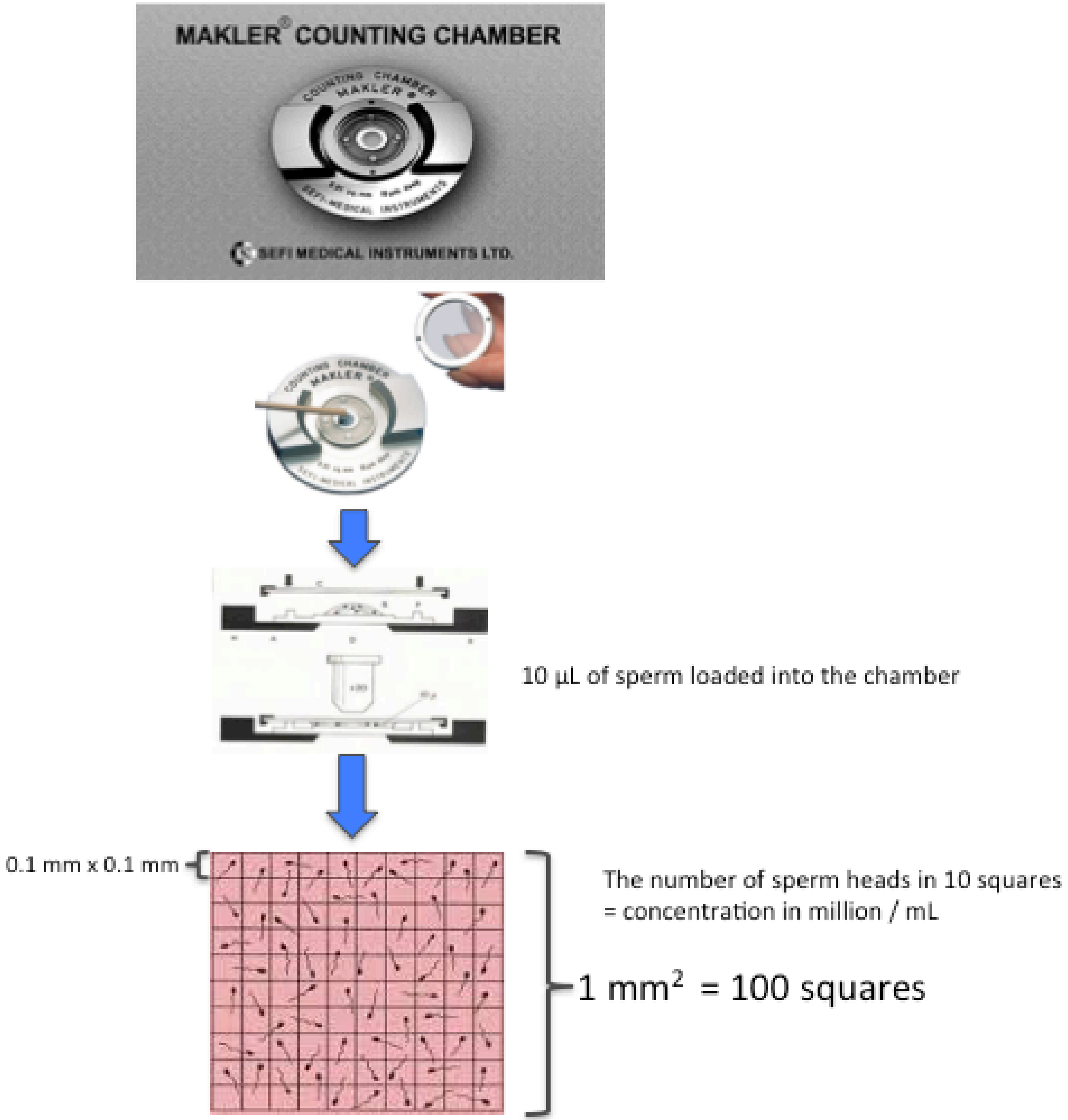


Motility: scoring motility is done on a scale of 0 to 5 with 0 equating to sperm that is not moving (non-motile), 1) sperm with tail movement ("twitching") but no forward progression, 2) some forward progression but slow and often swimming in circles, 3) tail moving and a moderate forward speed of progression (SOP), 4) tail moving vigorously and a good forward SOP, 5) sperm swimming very fast crossing the field of view quickly (this speed is rarely observed in boreal toad sperm) (Figure 7).

**Figure 7.** Motility scoring of boreal toad sperm, categories is subjectively scored from 0-5.

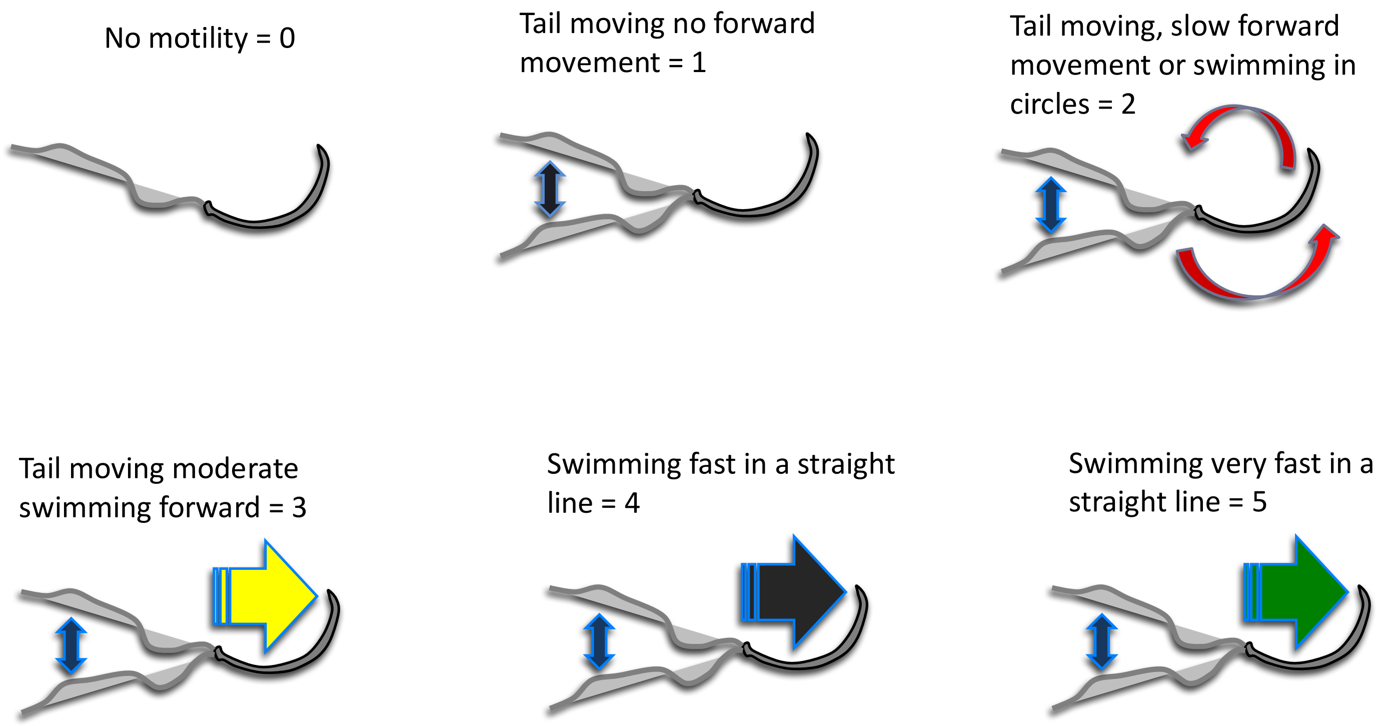


*Sperm morphology and live/dead staining.*

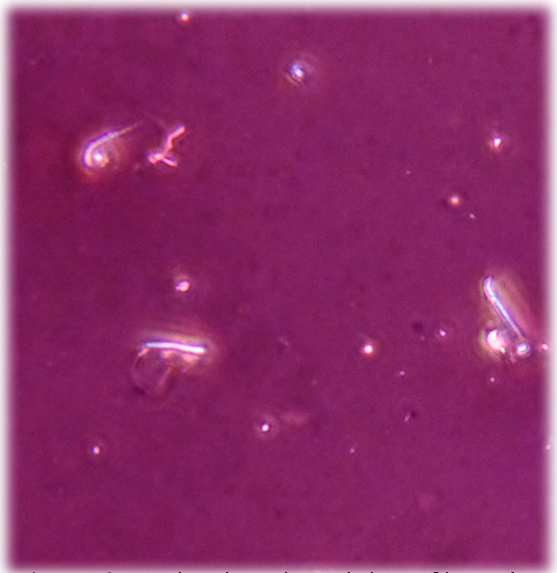

**Figure 8.** Eosin-nigrosin staining of boreal toad sperm with both live (unstained-white heads) and one dead sperm (pink stained head)

A method used to assess the quality of a sperm sample, is called live/dead staining. Live sperm can be differentiated from dead using Eosin / nigrosin, which relies on the ability of live sperm to exclude dyes (eosin-nigrosin) from entering the head and permeating the nucleus. A protocol for making eosin-nigrosin is in Appendix 1.

To stain sperm in a sample, 10 µL of eosin–nigrosin stain must be added to 10 µL of spermic urine on a glass slide and mixed well.

2. The slide is then allowed to air-dry but should be read within the hour, as dye will eventually permeate live sperm.

4. Sperm on the slide are then examined under ×40 with a bright field microscope. Live spermatozoa are left unstained (membrane-intact) and dead spermatozoa stain pink or red (membrane-damaged) (Figure 5.).

5. Count at least 100 sperm or as many as is possible, tally the numbers of live and dead spermatozoa using a laboratory counter, and calculate the percentage of live spermatozoa.

6. For boreal toads we use 80% as the lower reference limit for a good sperm sample (membrane-intact spermatozoa).

*In Vitro Fertilization (IVF)*

To conduct IVF in anurans, sperm is dropped/pipetted onto jellied eggs in a petri dish in a procedure known as *dry fertilization.* It is recommended that sperm be collected on the day oviposition is expected; however, good quality boreal toad sperm can be stored for up to 24 hr prior to performing IVF. Note that with increased storage time, motility and sperm quality can become depleted and may result in lower fertilization rates. Sperm concentration and motility should be determined before use to ensure the successful outcomes. Sperm concentrations of $1.0 \times 10^6$ are considered optimum for fertilizing 50-100 eggs, although this has not been experimentally confirmed. Motility is another important parameter to consider when performing IVF; however, less motile sperm do not necessarily reflect a sperms ability to fertilize. Sperm that show lower motility may still be able to fertilize eggs when pipetted directly onto them. Table 4 lists some basic parameters to consider when assessing sperm for IVF. To get a better assessment of sperm viability live/dead staining with Eosin-nigrosin is recommended (mentioned above). Eosin-nigrosin staining is an effective and simple way to assess the percentage of live sperm in a sample in the lab as well as the field.

**Table 5.** Spermic urine assessment parameters for performing IVF

| **Sperm urine collection times post hormone injection** | **Concentration million / mL** | **Motility** | | | **Viability staining** |
|---|---|---|---|---|---|
| | | **% moving** | **% forward progressing** | **speed of progression (SOP)** | **eosin/nigrosin % live sperm** |
| 2, 3, 5, 7, 9 & 12 hr post injection | 1,000,000 sperm or 100,000 / 100 µL | >70 | >40 | ≥3 | >80 |

Once spermic urine has been collected samples can be pooled if extra volume is needed. Preferably, pooled samples should be similar in motility and viability assessments.

*IVF protocol*

In a small petri dish or plastic container place freshly deposited or squeezed eggs. Add spermic urine at a concentration of 1 million per milliliter or 100,000 sperm / 100 µL and incubate 5-10 minutes without water ("dry fertilization") (Hollinger and Corton, 1980). Then eggs should be slowly flooded with tap or reconstituted distilled water and set aside for cleavage rate evaluation within 2-3 hours. Figure 3 shows an outline of the three steps involved in IVF. It is important to minimize disturbing the eggs once deposited into a Petri dish for fertilization as excess movement can cause the eggs to artificially cleave (auto- activate or parthonogenesis); These auto-activated eggs may appear to develop signs of early cleavage similar to a fertilized embryo, but are distinguishable by their asymmetrical shape and will not be fertilized.

**Figure 9.** Dry fertilization of eggs. Pipette spermic urine at concentrations of 100,000 sperm / 100 µL directly onto 50-100 "dry" eggs and incubate at room temperature for 5-10 minutes. Flood fertilized eggs with reconstituted or aged tap water.

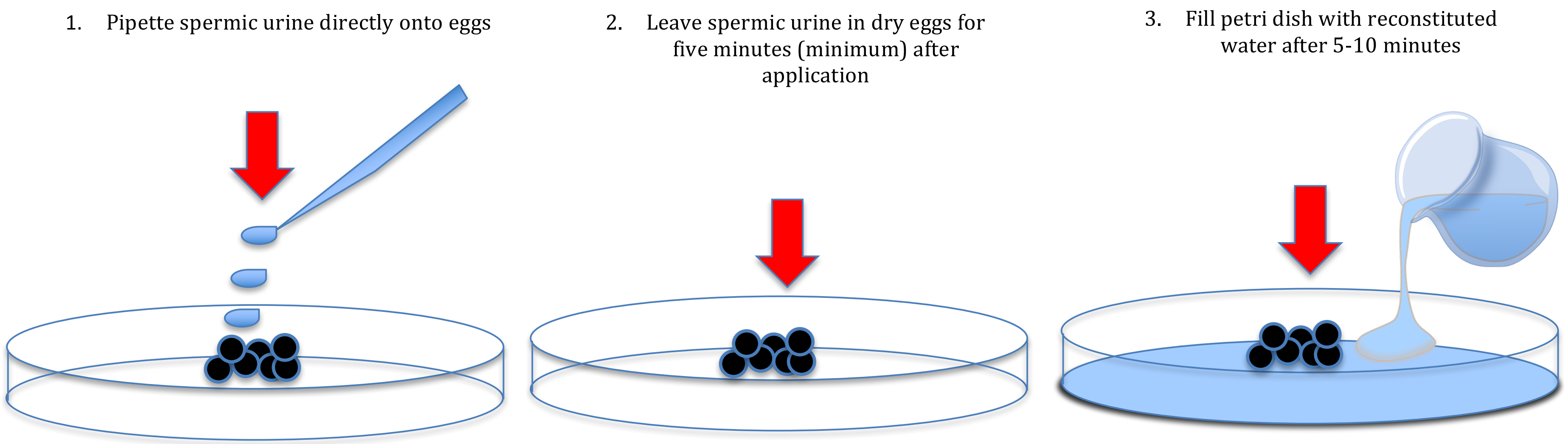

#### APPENDIX I

*Making Eosin-nigrosin dye (Mokovstev and Librach, 2013)*

1. Eosin–nigrosin stain: Dissolve 0.67 g eosin Y (color index 45380) and 0.9 g of sodium chloride in 100 mL of distilled water in a glass beaker placed on a stirring hot plate (see Note 1). Heat gently and add 10.0 g nigrosin (color index 50420) and dissolve it before bringing the stain to a boil. As soon as boiling is observed, remove the beaker from the hot plate and allow it to cool to room temperature. Strain the solution using filter paper, seal, and store at 4 °C in a dark glass bottle (see Note 2).
2. Phosphate-Buffered Saline (PBS): Dissolve one packet of Sigma PBS, pH 7.4 powder in 1,000 mL of distilled water to obtain a 0.01 M PBS solution (see Note 3). Store at room temperature for up to 6 months.
3. Microscope with bright field optics and ×100 oil immersion objective.
4. Stirring hot plate.
5. Microscope slides, 25 × 75 × 1 mm.

6. Coverslips, 22×50 mm, #1 thickness.
7. Disposable Pasteur pipettes.
8. Microtubes 1.5 mL.
9. Mounting medium.
10. Laboratory counter.
11. Laboratory timer.
12. Use a magnetic stir bar to dissolve ingredients for eosin– nigrosin staining.
13. Warm the stain to room temperature before use.
