## Supplementary Odds Ratio Tables for "Hormonal induction, quality assessments and the influence of seasonality on male reproductive viability in a long-term managed *ex situ* breeding colony of Southern Rocky Mountain Boreal toads, *Anaxyrus boreas boreas*"

Boreal Toad Manuscript for Animals supplementary tables.

**Supplementary Table 1**. Estimated marginal means for Boreal toad concentrations.

| Model | ZINB16b <- glmmTMB(sperm ~ month_f + offset(log_scale_ml_inv) + (1\|nasrfid), data=data5, ziformula=~hormone, family=nbinom2) |
| --- | --- |

| hormone | response | SE | df | lower.CL | upper.CL |
| --- | --- | --- | --- | --- | --- |
| Amphiplex | 9986 | 5301 | 3818 | 3527 | 28274 |
| GnRHA | 6033 | 3413 | 3818 | 1990 | 18293 |
| hCG | 40395 | 19987 | 3818 | 15312 | 106568 |
| hCG-GnRHA | 76077 | 38430 | 3818 | 28258 | 204819 |
| month_f = Jul: |  |  |  |  |  |
| hormone | response | SE | df | lower.CL | upper.CL |
| Amphiplex | 16573 | 8480 | 3818 | 6077 | 45194 |
| GnRHA | 10013 | 5886 | 3818 | 3163 | 31700 |
| hCG | 67043 | 33687 | 3818 | 25033 | 179553 |
| hCG-GnRHA | 126265 | 63684 | 3818 | 46971 | 339418 |
| month_f = Aug: |  |  |  |  |  |
| hormone | response | SE | df | lower.CL | upper.CL |
| Amphiplex | 5314 | 2826 | 3818 | 1874 | 15074 |
| GnRHA | 3211 | 1838 | 3818 | 1045 | 9864 |
| hCG | 21498 | 10660 | 3818 | 8132 | 56835 |
| hCG-GnRHA | 40489 | 20734 | 3818 | 14836 | 110499 |
| month_f = Sep: |  |  |  |  |  |
| hormone | response | SE | df | lower.CL | upper.CL |
| Amphiplex | 8644 | 4730 | 3818 | 2957 | 25273 |
| GnRHA | 5223 | 2966 | 3818 | 1715 | 15904 |
| hCG | 34969 | 17966 | 3818 | 12771 | 95750 |
| hCG-GnRHA | 65859 | 34809 | 3818 | 23366 | 185630 |

**Supplementary Table 2**. Concentration odds ratio according to hormone.

| contrast | ratio | SE | df | lower.CL | upper.CL |  |
| --- | --- | --- | --- | --- | --- | --- |
| (hCG-GnRHA) / GnRHA | 12.61 | 5.944 | 3818 | 5.005 | 31.77 | * |
| (hCG-GnRHA) / Amphiplex | 7.619 | 2.004 | 3818 | 4.549 | 12.76 | * |
| hCG / GnRHA | 6.695 | 2.929 | 3818 | 2.84 | 15.78 | * |
| hCG / Amphiplex | 4.045 | 1.112 | 3818 | 2.359 | 6.94 | * |
| (hCG-GnRHA) / hCG | 1.883 | 0.336 | 3818 | 1.328 | 2.67 | * |
| Amphiplex / GnRHA | 1.6551 | 0.8379 | 3818 | 0.6134 | 4.466 |  |

**Supplementary table 3.** Concentration odds ratio according to month.

| contrast | ratio | SE | df | lower.CL | upper.CL |  |
| --- | --- | --- | --- | --- | --- | --- |
| Jun / Aug | 3.119 | 0.468 | 3818 | 2.324 | 4.184 | * |
| Jul / Aug | 1.917 | 0.3639 | 3818 | 1.322 | 2.781 | * |
| Jun / Jul | 1.879 | 0.248 | 3818 | 1.451 | 2.434 | * |
| Sep / Aug | 1.66 | 0.235 | 3818 | 1.257 | 2.191 | * |
| Jul / Sep | 1.627 | 0.281 | 3818 | 1.159 | 2.284 | * |
| Jun / Sep | 1.155 | 0.1906 | 3818 | 0.836 | 1.596 | * |

**Supplementary table 4.** Estimated marginal means for Total motility.

| time_f | hormone | prob | SE | df | lower.CL | upper.CL |
| --- | --- | --- | --- | --- | --- | --- |
| 2 | Amphiplex | 0.729 | 0.0847 | 670 | 0.537 | 0.862 |
|  | GnRHA | 0.624 | 0.1018 | 670 | 0.415 | 0.796 |
|  | hCG | 0.785 | 0.0665 | 670 | 0.627 | 0.888 |
|  | hCG-GnRHA | 0.741 | 0.076 | 670 | 0.568 | 0.861 |
| 3 | Amphiplex | 0.75 | 0.0799 | 670 | 0.565 | 0.874 |
|  | GnRHA | 0.65 | 0.0982 | 670 | 0.443 | 0.813 |
|  | hCG | 0.803 | 0.0619 | 670 | 0.654 | 0.898 |
|  | hCG-GnRHA | 0.761 | 0.0715 | 670 | 0.596 | 0.874 |
| 5 | Amphiplex | 0.767 | 0.0769 | 670 | 0.586 | 0.884 |
|  | GnRHA | 0.67 | 0.0956 | 670 | 0.465 | 0.826 |
|  | hCG | 0.817 | 0.0586 | 670 | 0.674 | 0.906 |
|  | hCG-GnRHA | 0.777 | 0.0682 | 670 | 0.617 | 0.883 |
| 7 | Amphiplex | 0.787 | 0.0725 | 670 | 0.613 | 0.896 |
|  | GnRHA | 0.695 | 0.0913 | 670 | 0.495 | 0.842 |
|  | hCG | 0.834 | 0.0544 | 670 | 0.699 | 0.916 |
|  | hCG-GnRHA | 0.797 | 0.064 | 670 | 0.644 | 0.895 |
| 9 | Amphiplex | 0.786 | 0.0729 | 670 | 0.61 | 0.895 |
|  | GnRHA | 0.694 | 0.0924 | 670 | 0.491 | 0.842 |
|  | hCG | 0.833 | 0.0548 | 670 | 0.697 | 0.915 |
|  | hCG-GnRHA | 0.796 | 0.0642 | 670 | 0.642 | 0.894 |
| 12 | Amphiplex | 0.726 | 0.0857 | 670 | 0.532 | 0.861 |
|  | GnRHA | 0.621 | 0.1022 | 670 | 0.411 | 0.793 |
|  | hCG | 0.783 | 0.0668 | 670 | 0.625 | 0.886 |
|  | hCG-GnRHA | 0.738 | 0.0763 | 670 | 0.565 | 0.859 |
| 24 | Amphiplex | 0.621 | 0.1011 | 670 | 0.413 | 0.792 |
|  | GnRHA | 0.503 | 0.1088 | 670 | 0.301 | 0.704 |
|  | hCG | 0.69 | 0.0839 | 670 | 0.507 | 0.828 |
|  | hCG-GnRHA | 0.635 | 0.0916 | 670 | 0.445 | 0.791 |

**Supplementary table 5.** Motility odds ratios by Hormone.

| contrast | odds.ratio | SE | df | lower.CL upp | upper.CL |  |
| --- | --- | --- | --- | --- | --- | --- |
| hCG / GnRHA | 2.198 | 0.5242 | 670 | 1.376 | 3.511 | * |
| hCG / Amphiplex | 1.358 | 0.3072 | 670 | 0.871 | 2.118 | * |
| hCG / (hCG-GnRHA) | 1.279 | 0.156 | 670 | 1.006 | 1.625 | * |
| (hCG-GnRHA) / Amphiplex | 1.062 | 0.2325 | 670 | 0.691 | 1.632 | * |
| (hCG-GnRHA) / GnRHA | 1.718 | 0.4201 | 670 | 1.063 | 2.777 | * |
| Amphiplex / GnRHA | 1.618 | 0.499 | 670 | 0.883 | 2.965 | * |

**Supplementary table 6.** Total motility odds ratios by hormone.

| contrast | odds.ratio | SE | df | lower.CL upp | upper.CL |  |
| --- | --- | --- | --- | --- | --- | --- |
| hCG / GnRHA | 2.198 | 0.5242 | 670 | 1.376 | 3.511 | * |
| hCG / Amphiplex | 1.358 | 0.3072 | 670 | 0.871 | 2.118 |  |
| hCG / (hCG-GnRHA) | 1.279 | 0.156 | 670 | 1.006 | 1.625 | * |
| (hCG-GnRHA) / Amphiplex | 1.062 | 0.2325 | 670 | 0.691 | 1.632 |  |
| (hCG-GnRHA) / GnRHA | 1.718 | 0.4201 | 670 | 1.063 | 2.777 | * |
| Amphiplex / GnRHA | 1.618 | 0.499 | 670 | 0.883 | 2.965 |  |

**Supplementary table 7.** Motility odds ratios by time-point.

| contrast | odds.ratio | SE | df | lower.CL | upper.CL |  |
| --- | --- | --- | --- | --- | --- | --- |
| time_f2 / time_f12 | 1.014 | 0.1229 | 670 | 0.8 | 1.287 |  |
| time_f2 / time_f24 | 1.64 | 0.2002 | 670 | 1.291 | 2.084 | * |
| time_f3 / time_f2 | 1.118 | 0.133 | 670 | 0.886 | 1.413 |  |
| time_f3 / time_f12 | 1.134 | 0.1337 | 670 | 0.9 | 1.43 |  |
| time_f3 / time_f24 | 1.834 | 0.2165 | 670 | 1.455 | 2.313 | * |
| time_f5 / time_f12 | 1.242 | 0.1462 | 670 | 0.985 | 1.565 |  |
| time_f5 / time_f2 | 1.224 | 0.1471 | 670 | 0.967 | 1.55 |  |
| time_f5 / time_f3 | 1.095 | 0.1272 | 670 | 0.871 | 1.375 |  |
| time_f5 / time_f24 | 2.008 | 0.2378 | 670 | 1.591 | 2.534 | * |
| time_f7 / time_f2 | 1.375 | 0.1697 | 670 | 1.079 | 1.752 | * |
| time_f7 / time_f3 | 1.229 | 0.1471 | 670 | 0.972 | 1.555 |  |
| time_f7 / time_f5 | 1.123 | 0.1339 | 670 | 0.889 | 1.419 |  |
| time_f7 / time_f9 | 1.009 | 0.1247 | 670 | 0.791 | 1.286 |  |
| time_f7 / time_f12 | 1.395 | 0.1674 | 670 | 1.102 | 1.765 | * |
| time_f7 / time_f24 | 2.255 | 0.272 | 670 | 1.779 | 2.857 | * |
| time_f9 / time_f2 | 1.363 | 0.1702 | 670 | 1.067 | 1.742 | * |
| time_f9 / time_f3 | 1.219 | 0.1479 | 670 | 0.96 | 1.547 |  |
| time_f9 / time_f5 | 1.113 | 0.1352 | 670 | 0.877 | 1.413 |  |
| time_f9 / time_f12 | 1.382 | 0.1678 | 670 | 1.089 | 1.755 | * |
| time_f9 / time_f24 | 2.235 | 0.2739 | 670 | 1.757 | 2.843 | * |
| time_f12 / time_f24 | 1.617 | 0.1921 | 670 | 1.281 | 2.042 | * |

**Supplementary table 8.** Estimated Marginal means of forward progressive motility by hormone.

| Model | m14b=glmmTMB(p_fp ~ hormone + time_f + (1\|nasrfid)+(1\|season),  weights = sum_nm_fp, data=data7, family=betabinomial(link="logit")) |
| --- | --- |

| time | hormone | prob | SE | df | lower.CL | upper.CL |
| --- | --- | --- | --- | --- | --- | --- |
| 2 | Amphiplex | 0.464 | 0.1372 | 669 | 0.226 | 0.719 |
|  | GnRHA | 0.326 | 0.1225 | 669 | 0.139 | 0.591 |
|  | hCG | 0.628 | 0.1201 | 669 | 0.381 | 0.823 |
|  | hCG-GnRHA | 0.591 | 0.1246 | 669 | 0.344 | 0.799 |
| 3 | Amphiplex | 0.534 | 0.1366 | 669 | 0.281 | 0.771 |
|  | GnRHA | 0.391 | 0.1323 | 669 | 0.177 | 0.656 |
|  | hCG | 0.692 | 0.1091 | 669 | 0.451 | 0.86 |
|  | hCG-GnRHA | 0.658 | 0.1158 | 669 | 0.412 | 0.841 |
| 5 | Amphiplex | 0.574 | 0.1353 | 669 | 0.312 | 0.799 |
|  | GnRHA | 0.429 | 0.1359 | 669 | 0.202 | 0.691 |
|  | hCG | 0.724 | 0.1023 | 669 | 0.49 | 0.878 |
|  | hCG-GnRHA | 0.692 | 0.1096 | 669 | 0.45 | 0.861 |
| 7 | Amphiplex | 0.626 | 0.1299 | 669 | 0.36 | 0.833 |
|  | GnRHA | 0.483 | 0.1381 | 669 | 0.24 | 0.735 |
|  | hCG | 0.766 | 0.0918 | 669 | 0.545 | 0.899 |
|  | hCG-GnRHA | 0.737 | 0.0999 | 669 | 0.505 | 0.885 |
| 9 | Amphiplex | 0.615 | 0.1316 | 669 | 0.349 | 0.826 |
|  | GnRHA | 0.471 | 0.1385 | 669 | 0.23 | 0.727 |
|  | hCG | 0.757 | 0.0943 | 669 | 0.533 | 0.895 |
|  | hCG-GnRHA | 0.728 | 0.102 | 669 | 0.493 | 0.88 |
| 12 | Amphiplex | 0.522 | 0.1383 | 669 | 0.269 | 0.764 |
|  | GnRHA | 0.378 | 0.1309 | 669 | 0.17 | 0.645 |
|  | hCG | 0.681 | 0.1115 | 669 | 0.438 | 0.854 |
|  | hCG-GnRHA | 0.646 | 0.1178 | 669 | 0.399 | 0.834 |
| 24 | Amphiplex | 0.402 | 0.1335 | 669 | 0.185 | 0.667 |
|  | GnRHA | 0.273 | 0.111 | 669 | 0.111 | 0.53 |
|  | hCG | 0.568 | 0.126 | 669 | 0.324 | 0.783 |
|  | hCG-GnRHA | 0.53 | 0.1286 | 669 | 0.29 | 0.756 |

**Supplementary Table 9.** Forward progressive motility odds ratios by hormone.

| contrast | odds.ratio | SE | df | lower.CL | upper.CL |  |
| --- | --- | --- | --- | --- | --- | --- |
| hCG / GnRHA | 3.5 | 0.959 | 669 | 2.044 | 5.99 | * |
| hCG / Amphiplex | 1.954 | 0.513 | 669 | 1.167 | 3.27 | * |
| hCG / (hCG-GnRHA) | 1.168 | 0.1639 | 669 | 0.887 | 1.538 |  |
| (hCG-GnRHA) / GnRHA | 2.997 | 0.846 | 669 | 1.722 | 5.22 | * |
| (hCG-GnRHA) / Amphiplex | 1.674 | 0.429 | 669 | 1.012 | 2.77 | * |
| Amphiplex / GnRHA | 1.791 | 0.6434 | 669 | 0.884 | 3.626 |  |

**Supplementary Table 10.** Forward progressive motility odds ratios by time-point.

| contrast | odds.ratio | SE | df | lower.CL | upper.CL |  |
| --- | --- | --- | --- | --- | --- | --- |
| time_f2 / time_f24 | 1.284 | 0.1899 | 669 | 0.961 | 1.717 |  |
| time_f3 / time_f2 | 1.327 | 0.1874 | 669 | 1.006 | 1.752 | * |
| time_f3 / time_f12 | 1.052 | 0.146 | 669 | 0.801 | 1.382 |  |
| time_f3 / time_f24 | 1.705 | 0.2418 | 669 | 1.29 | 2.252 | * |
| time_f5 / time_f2 | 1.556 | 0.2201 | 669 | 1.179 | 2.054 | * |
| time_f5 / time_f3 | 1.172 | 0.1582 | 669 | 0.899 | 1.528 |  |
| time_f5 / time_f12 | 1.233 | 0.1695 | 669 | 0.942 | 1.615 |  |
| time_f5 / time_f24 | 1.998 | 0.2823 | 669 | 1.514 | 2.637 | * |
| time_f7 / time_f2 | 1.937 | 0.2803 | 669 | 1.458 | 2.574 | * |
| time_f7 / time_f3 | 1.459 | 0.2018 | 669 | 1.112 | 1.914 | * |
| time_f7 / time_f5 | 1.245 | 0.1696 | 669 | 0.953 | 1.627 |  |
| time_f7 / time_f9 | 1.048 | 0.1463 | 669 | 0.797 | 1.379 |  |
| time_f7 / time_f12 | 1.535 | 0.2137 | 669 | 1.168 | 2.018 | * |
| time_f7 / time_f24 | 2.488 | 0.3558 | 669 | 1.879 | 3.295 | * |
| time_f9 / time_f2 | 1.848 | 0.2682 | 669 | 1.39 | 2.457 | * |
| time_f9 / time_f3 | 1.392 | 0.194 | 669 | 1.059 | 1.83 | * |
| time_f9 / time_f5 | 1.188 | 0.1639 | 669 | 0.906 | 1.557 |  |
| time_f9 / time_f12 | 1.465 | 0.2054 | 669 | 1.112 | 1.929 | * |
| time_f9 / time_f24 | 2.373 | 0.343 | 669 | 1.787 | 3.152 | * |
| time_f12 / time_f2 | 1.262 | 0.1821 | 669 | 0.95 | 1.675 |  |
| time_f12 / time_f24 | 1.62 | 0.233 | 669 | 1.222 | 2.149 | * |

**Supplementary Table 11.** Estimated marginal means of live-dead cell (viability).

| Time | hormone | prob | SE | df | lower.CL | upper.CL |
| --- | --- | --- | --- | --- | --- | --- |
| 2 | Amphiplex | 0.894 | 0.0267 | 677 | 0.829 | 0.937 |
|  | GnRHA | 0.854 | 0.031 | 677 | 0.782 | 0.905 |
|  | hCG | 0.878 | 0.0177 | 677 | 0.839 | 0.909 |
|  | hCG-GnRHA | 0.888 | 0.0167 | 677 | 0.85 | 0.916 |
| 3 | Amphiplex | 0.881 | 0.0294 | 677 | 0.811 | 0.928 |
|  | GnRHA | 0.837 | 0.0338 | 677 | 0.76 | 0.893 |
|  | hCG | 0.863 | 0.0185 | 677 | 0.822 | 0.895 |
|  | hCG-GnRHA | 0.874 | 0.0179 | 677 | 0.834 | 0.905 |
| 5 | Amphiplex | 0.884 | 0.0292 | 677 | 0.813 | 0.93 |
|  | GnRHA | 0.841 | 0.0329 | 677 | 0.765 | 0.895 |
|  | hCG | 0.866 | 0.018 | 677 | 0.827 | 0.898 |
|  | hCG-GnRHA | 0.877 | 0.0176 | 677 | 0.838 | 0.907 |
| 7 | Amphiplex | 0.887 | 0.0283 | 677 | 0.819 | 0.932 |
|  | GnRHA | 0.845 | 0.0322 | 677 | 0.771 | 0.898 |
|  | hCG | 0.87 | 0.0177 | 677 | 0.831 | 0.901 |
|  | hCG-GnRHA | 0.88 | 0.0174 | 677 | 0.842 | 0.91 |
| 9 | Amphiplex | 0.894 | 0.027 | 677 | 0.828 | 0.936 |
|  | GnRHA | 0.853 | 0.0313 | 677 | 0.781 | 0.905 |
|  | hCG | 0.877 | 0.0172 | 677 | 0.839 | 0.907 |
|  | hCG-GnRHA | 0.887 | 0.0167 | 677 | 0.85 | 0.916 |
| 12 | Amphiplex | 0.88 | 0.0299 | 677 | 0.808 | 0.928 |
|  | GnRHA | 0.836 | 0.0346 | 677 | 0.757 | 0.893 |
|  | hCG | 0.862 | 0.0187 | 677 | 0.821 | 0.895 |
|  | hCG-GnRHA | 0.873 | 0.0183 | 677 | 0.832 | 0.905 |
| 24 | Amphiplex | 0.833 | 0.0389 | 677 | 0.743 | 0.897 |
|  | GnRHA | 0.776 | 0.0434 | 677 | 0.679 | 0.85 |
|  | hCG | 0.809 | 0.0243 | 677 | 0.757 | 0.853 |
|  | hCG-GnRHA | 0.823 | 0.0238 | 677 | 0.772 | 0.865 |

**Supplementary Table 12.** Vitality odds ration by hormone.

| contrast | odds.ratio | SE | df | lower.CL | upper.CL |
| --- | --- | --- | --- | --- | --- |
| Amphiplex / GnRHA | 1.445 | 0.468 | 677 | 0.766 | 2.73 |
| Amphiplex / hCG | 1.178 | 0.325 | 677 | 0.685 | 2.03 |
| Amphiplex / (hCG-GnRHA) | 1.073 | 0.294 | 677 | 0.626 | 1.84 |
| (hCG-GnRHA) / GnRHA | 1.347 | 0.327 | 677 | 0.836 | 2.17 |
| (hCG-GnRHA) / hCG | 1.098 | 0.156 | 677 | 0.831 | 1.45 |
| hCG / GnRHA | 1.227 | 0.288 | 677 | 0.773 | 1.95 |

**Supplementary Table 13.** Vitality odds ration by time point.

| contrast | odds.ratio | SE | df | lower.CL | upper.CL |  |
| --- | --- | --- | --- | --- | --- | --- |
| time_f2 / time_f3 | 1.141 | 0.139 | 677 | 0.898 | 1.45 |  |
| time_f2 / time_f5 | 1.112 | 0.136 | 677 | 0.875 | 1.41 |  |
| time_f2 / time_f7 | 1.074 | 0.134 | 677 | 0.841 | 1.37 |  |
| time_f2 / time_f9 | 1.008 | 0.128 | 677 | 0.785 | 1.29 |  |
| time_f2 / time_f12 | 1.15 | 0.144 | 677 | 0.899 | 1.47 |  |
| time_f2 / time_f24 | 1.693 | 0.212 | 677 | 1.323 | 2.17 | * |
| time_f3 / time_f12 | 1.007 | 0.117 | 677 | 0.801 | 1.27 |  |
| time_f3 / time_f24 | 1.484 | 0.173 | 677 | 1.181 | 1.86 | * |
| time_f5 / time_f3 | 1.026 | 0.1152 | 677 | 0.823 | 1.279 |  |
| time_f5 / time_f12 | 1.034 | 0.119 | 677 | 0.825 | 1.29 |  |
| time_f5 / time_f24 | 1.522 | 0.175 | 677 | 1.215 | 1.91 | * |
| time_f7 / time_f3 | 1.062 | 0.1223 | 677 | 0.848 | 1.332 |  |
| time_f7 / time_f5 | 1.035 | 0.1174 | 677 | 0.829 | 1.293 |  |
| time_f7 / time_f12 | 1.07 | 0.125 | 677 | 0.851 | 1.35 |  |
| time_f7 / time_f24 | 1.576 | 0.184 | 677 | 1.253 | 1.98 | * |
| time_f9 / time_f3 | 1.133 | 0.1336 | 677 | 0.898 | 1.428 |  |
| time_f9 / time_f5 | 1.104 | 0.1283 | 677 | 0.878 | 1.387 |  |
| time_f9 / time_f7 | 1.066 | 0.1257 | 677 | 0.846 | 1.344 |  |
| time_f9 / time_f12 | 1.141 | 0.137 | 677 | 0.902 | 1.44 |  |
| time_f9 / time_f24 | 1.68 | 0.2 | 677 | 1.33 | 2.12 | * |
| time_f12 / time_f24 | 1.473 | 0.173 | 677 | 1.169 | 1.86 | * |

**Supplementary Table 14.** Estimated Marginal Means of hormone by month.

Model: m4=glmer(p_mot ~ hormone + month + (1|olre), weights = spermtot,data=data6,family=binomial)

|  | hormone | prob | SE | df | asymp.LCL | asymp.UCL |
| --- | --- | --- | --- | --- | --- | --- |
| Aug | Amphiplex | 0.627 | 0.0686 | Inf | 0.486 | 0.749 |
|  | GnRHA | 0.645 | 0.0761 | Inf | 0.486 | 0.777 |
|  | hCG | 0.744 | 0.0271 | Inf | 0.687 | 0.794 |
|  | hCG-GnRHA | 0.663 | 0.0353 | Inf | 0.59 | 0.728 |
| Jul | Amphiplex | 0.642 | 0.0613 | Inf | 0.515 | 0.752 |
|  | GnRHA | 0.66 | 0.0704 | Inf | 0.512 | 0.782 |
|  | hCG | 0.757 | 0.0214 | Inf | 0.712 | 0.796 |
|  | hCG-GnRHA | 0.677 | 0.0255 | Inf | 0.625 | 0.725 |
| Jun | Amphiplex | 0.694 | 0.0574 | Inf | 0.572 | 0.794 |
|  | GnRHA | 0.71 | 0.0659 | Inf | 0.567 | 0.821 |
|  | hCG | 0.797 | 0.0199 | Inf | 0.755 | 0.833 |
|  | hCG-GnRHA | 0.726 | 0.0306 | Inf | 0.663 | 0.782 |
| Sep | Amphiplex | 0.611 | 0.0732 | Inf | 0.462 | 0.741 |
|  | GnRHA | 0.629 | 0.0739 | Inf | 0.477 | 0.759 |
|  | hCG | 0.731 | 0.0328 | Inf | 0.662 | 0.79 |
|  | hCG-GnRHA | 0.647 | 0.0386 | Inf | 0.568 | 0.719 |

**Supplementary Table 15.** Motility Odds Rations by hormone.

| contrast | odds.ratio | SE | df | LCL | UCL |  |
| --- | --- | --- | --- | --- | --- | --- |
| hCG / Amphiplex | 1.733 | 0.4702 | Inf | 1.018 | 2.949 | * |
| hCG / GnRHA | 1.603 | 0.5023 | Inf | 0.867 | 2.963 |  |
| (hCG-GnRHA) / Amphiplex | 1.171 | 0.3264 | Inf | 0.678 | 2.022 |  |
| hCG / (hCG-GnRHA) | 1.48 | 0.192 | Inf | 1.147 | 1.91 | * |
| (hCG-GnRHA) / GnRHA | 1.083 | 0.3439 | Inf | 0.581 | 2.018 |  |
| GnRHA / Amphiplex | 1.081 | 0.4324 | Inf | 0.493 | 2.368 |  |

**Supplementary Table 16.** Motility Odd ratios by month.

| contrast | odds.ratio | SE | df | LCL | UCL |  |
| --- | --- | --- | --- | --- | --- | --- |
| Jun / Aug | 1.352 | 0.242 | Inf | 0.953 | 1.92 | * |
| Jun / Jul | 1.265 | 0.194 | Inf | 0.937 | 1.71 | * |
| Jun / Sep | 1.446 | 0.283 | Inf | 0.986 | 2.12 | * |
| Jul / Sep | 1.144 | 0.207 | Inf | 0.802 | 1.63 |  |
| Aug / Sep | 1.07 | 0.219 | Inf | 0.716 | 1.6 |  |
| Jul / Aug | 1.069 | 0.177 | Inf | 0.773 | 1.48 |  |

**Supplementary table 17.** Estimated marginal means of vitality by month.

|  | hormone | prob | SE | df | asymp.LCL | asymp.UCL |
| --- | --- | --- | --- | --- | --- | --- |
| Aug | Amphiplex | 0.812 | 0.0437 | Inf | 0.711 | 0.883 |
|  | GnRHA | 0.885 | 0.03154 | Inf | 0.807 | 0.934 |
|  | hCG | 0.819 | 0.02052 | Inf | 0.776 | 0.856 |
|  | hCG-GnRHA | 0.831 | 0.02242 | Inf | 0.783 | 0.871 |
| Jul | Amphiplex | 0.893 | 0.02477 | Inf | 0.834 | 0.933 |
|  | GnRHA | 0.937 | 0.01691 | Inf | 0.895 | 0.963 |
|  | hCG | 0.898 | 0.01008 | Inf | 0.876 | 0.916 |
|  | hCG-GnRHA | 0.905 | 0.00945 | Inf | 0.885 | 0.922 |
| Jun | Amphiplex | 0.91 | 0.02153 | Inf | 0.858 | 0.944 |
|  | GnRHA | 0.948 | 0.01468 | Inf | 0.91 | 0.97 |
|  | hCG | 0.914 | 0.00913 | Inf | 0.894 | 0.93 |
|  | hCG-GnRHA | 0.92 | 0.01094 | Inf | 0.896 | 0.939 |
| Sep | Amphiplex | 0.856 | 0.03703 | Inf | 0.768 | 0.915 |
|  | GnRHA | 0.914 | 0.02316 | Inf | 0.857 | 0.95 |
|  | hCG | 0.863 | 0.01966 | Inf | 0.819 | 0.897 |
|  | hCG-GnRHA | 0.872 | 0.01865 | Inf | 0.831 | 0.904 |

**Supplementary table 18.** Vitality odd ratios by hormone.

| contrast | odds.ratio | SE | df | asymp.LCL | asymp.UCL |
| --- | --- | --- | --- | --- | --- |
| GnRHA / Amphiplex | 1.788 | 0.674 | Inf | 0.855 | 3.74 |
| GnRHA / hCG | 1.699 | 0.491 | Inf | 0.964 | 2.99 |
| GnRHA / (hCG-GnRHA) | 1.565 | 0.459 | Inf | 0.881 | 2.78 |
| hCG / Amphiplex | 1.053 | 0.277 | Inf | 0.628 | 1.76 |
| (hCG-GnRHA) / Amphiplex | 1.143 | 0.31 | Inf | 0.671 | 1.95 |
| (hCG-GnRHA) / hCG | 1.086 | 0.137 | Inf | 0.847 | 1.39 |

**Supplementary table 19.** Vitality odds ratio by month.

| contrast | odds.ratio | SE | df | asymp.LCL | asymp.UCL | |
| --- | --- | --- | --- | --- | --- | --- |
| Jun / Aug | 2.349 | 0.408 | Inf | 1.671 | 3.301 | * |
| Jun / Sep | 1.696 | 0.3255 | Inf | 1.165 | 2.471 | * |
| Jun / Jul | 1.209 | 0.176 | Inf | 0.91 | 1.608 |  |
| Jul / Aug | 1.942 | 0.314 | Inf | 1.414 | 2.667 | * |
| Jul / Sep | 1.403 | 0.2478 | Inf | 0.992 | 1.983 |  |
| Sep / Aug | 1.385 | 0.283 | Inf | 0.928 | 2.066 |  |
